# Antibiotics that Kill Gram-negative Bacteria by Restructuring the Outer Membrane Protein BamA

**DOI:** 10.1101/2024.12.16.628070

**Authors:** Seyed Majed Modaresi, Ryosuke Sugiyama, Nhan Dai Thien Tram, Roman P. Jakob, Chin-Soon Phan, Amir Ata Saei, Yohei Morishita, Tobias Mühlethaler, Joel Lim, Danilo Ritz, Preston Shi Yang Long, Phillipe A Lehner, Zhen Heng Lim, Morris Degen, Ziwei Yao, Timm Maier, Yuxin Hou, Jia Ying Lee, Jian Xu, Andrew Yeo Jung Yeat, Kenny Ting Sween Koh, Wei Yang Goh, Sharon Y. H. Ling, Patrina W. L. Chua, Mami Yamazaki, Pui Lai Rachel Ee, Sebastian Hiller, Brandon I Morinaka

**Author notes:** To whom correspondence may be addressed. (PLRE), (SH), or (BIM). These authors contributed equally (Seyed Majed Modaresi, Ryosuke Sugiyama, Nhan Dai Thien Tram). Present address: Institute for Medical Engineering & Science, Massachusetts Institute of Technology, Cambridge, MA, 02139, USA. Infectious Disease and Microbiome Program, Broad Institute of MIT and Harvard, Cambridge, MA, 02142, USA. Present address: Centre for Translational Microbiome Research, Department of Microbiology, Tumor and Cell Biology, Karolinska Institute, Stockholm 17165, Sweden (Amir Ata Saei). Present address: Latvian Institute of Organic Synthesis, Aizkraukles Street 21, LV-1006 Riga, Latvia.

## Abstract

The essential outer membrane protein insertase BamA has recently emerged as a valid target for killing Gram-negative bacteria. Bamabactins, competitive inhibitors targeting the lateral gate of BamA, disrupt the substrate folding process, compromise the outer membrane integrity, and lead to bacterial cell death. Despite their promise, the full pharmacological potential of bamabactins remains underexploited. We applied phylogenetic genome mining and synthetic biology to identify xenorceptides which selectively kill *Enterobacteriaceae*. Mode of action studies show that xenorceptide A2 integrates itself into BamA as an additional β-strand between β1 and β16 at the lateral gate, inducing a conformation of BamA that has not been observed before. Biological evaluation of xenorceptide A2 shows promising activity *in vitro* and *in vivo*, and limited resistance which differentiates it from other bamabactin antibiotics. Our data show that the chemical diversity of bamabactins is far greater than previously recognized and thus an attractive source for antibiotic discovery.

## Introduction

The Centers for Disease Control and Prevention and the World Health Organization have classified Carbapenem-resistant *Enterobacteriaceae* (CRE), including the Gram-negative *Klebsiella pneumoniae* and *Escherichia coli*, as top priority pathogens for which new antibiotics are urgently needed^1,2^. CRE pose an immediate threat due to their resistance to most antibiotic classes, as hospital-acquired CRE infections have increased by 35% in 2020 compared to 2019^3^. The augmented use of carbapenems and the spread of resistance mechanisms are major factors driving this rise^4,5^. Infections can be severe and often deadly by causing pneumonia, bloodstream infections, urinary tract infections, wound infections, and meningitis.^6^ Developing narrow-spectrum antibiotics is therefore crucial for reducing mortality and minimizing the risk of cross-resistance, providing a targeted approach to combat infections.

Microbial natural products have long been a vital source of antibiotics.^7^ In 2019, Lewis and coworkers reported their discovery of utilizing strains of *Xenorhabdus* and *Photorhabdus* – bacterial symbionts in nematode gut microbiomes – for their antibiotic potential^8^. Screening a *Photorhabdus* library revealed darobactin A from extracts of *Photorhabdus khaini*. Darobactin A is a 7-mer peptide with a fused bicyclic structure, and the founding member of a new antibiotic class we term here as bamabactins. Bamabactins are cyclophane-containing peptides that target the essential outer membrane protein (OMP) BamA.^9^ Their mechanism of action is a β-β interaction with BamA’s lateral gate, displacing native substrates. In 2022, dynobactin A was identified as a 10-mer bamabactin from *Photorhabdus australis* with two unlinked cyclophane rings^10^. Unlike darobactin A, dynobactin A extends three residues into the BamA β-barrel (BamA-β) lumen, this suggested that the active chemical space might extend much beyond these two compounds^11^. Despite these two interesting lead compounds for antibiotic development, BamA is not targeted by any clinically approved drug so far, which necessitates further exploration of this chemical space.

Expanding the chemical landscape based on phylogeny is instrumental in antibiotic discovery and development processes. Phylogenetically related antibiotics often possess unique scaffolds leading to optimized biological activity via various modes of action^12,13^. In parallel, synthetic biology offers a complementary approach to access targeted natural products through phylogeny-based genome mining^14–16^. In this approach, biosynthetic gene clusters (BGCs) are identified by phylogeny, and then promising BGCs are selected for compound production. Systematic heterologous expression in a uniform chassis, such as *E. coli,* can rapidly produce entire families of natural products. This strategy eliminates the need to culture diverse bacteria with specific growth conditions or genetic manipulation. In contrast, bioactivity-guided fractionation from crude extracts has several limitations, such as the frequent rediscovery of known compounds and challenges in accurately assessing the yield and potency of the produced natural product^17^. Genome mining and synthetic biology overcome these limitations by enabling the prediction of novel compounds and direct production at defined concentrations, allowing for accurate biological assessment of the novel compounds.

Here we used a combination of phylogeny-guided genome mining, synthetic biology, and structural biology to identify xenorceptides, peptide natural products, that kill Gram-negative bacteria by blocking the outer membrane protein BamA. Our findings reveal that xenorceptide A2 (hereafter called **2**) induces a complete restructuring of the transmembrane domain of BamA and is not susceptible to most of the mutations conferring resistance to formerly described bamabactins. Mode of action of **2** unveils an unprecedented phenomenon where our atomic model guides design of narrow spectrum antibiotics targeting BamA. Our results expand the bamabactin class to include xenorceptides that have a distinct mode of action to combat multi-drug resistant *Enterobactericeae*.

## Results

### Bioinformatic analysis show xenorceptides are prevalent within Enterobacteriaceae

Xenorceptide A1 (hereafter referred to as **1**), is a 12-mer cyclophane ribosomally synthesized and post-translationally modified peptide (RiPP) isolated from *Xenorhabdus nematophila* (Fig. 1a). **1** resulted from an effort to characterize and describe a new subclass of cyclophane RiPPs, *triceptides* (Extended data Fig. 1), but did not show antibacterial activity. The biosynthetic gene clusters (BGCs) that encode the xenorceptide natural product family were catalogued in the TIGRFAMs database as XYE (*<u>X</u>enorhabdus*, *<u>Y</u>ersinia*, and *<u>E</u>rwinia*) maturase systems.^18,19^ These BGCs encode a precursor peptide (A), radical SAM cyclophane synthase (B), aspartic acid protease (C), HlyD transporter (D), and protease-transporter (E) (Fig. 1a). The precursor peptides contain a conserved Gly-Gly motif as the target of the protease to release the modified core peptide.

**Figure 1.**
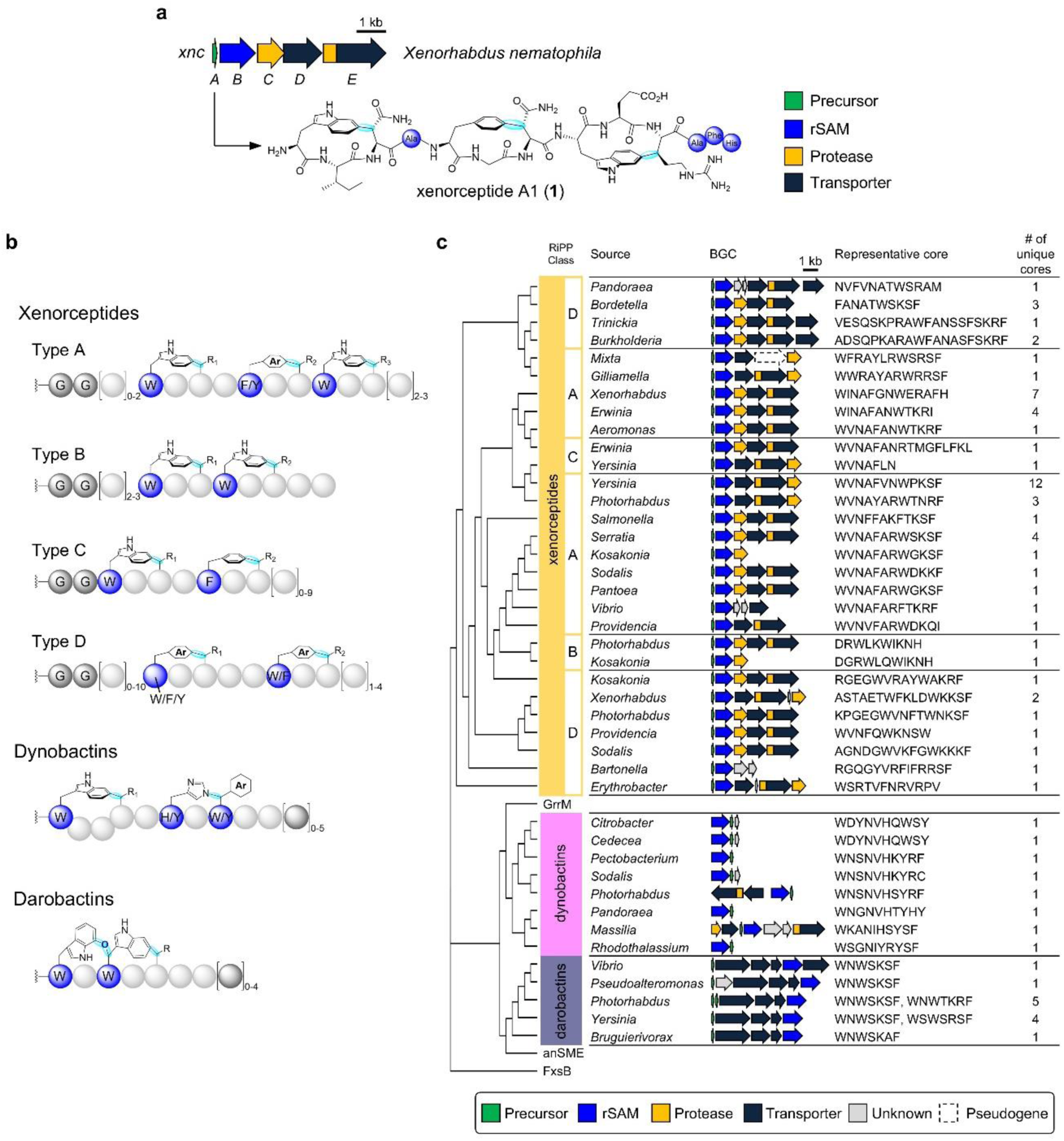
Bioinformatic mapping of xenorceptides. (a) Biosynthetic gene cluster for xenorceptide A1 (**1**). (b) Bamabactins (cyclophane RiPP antibiotics) that target BamA include xenorceptides, darobactins, and dynobactins. (c), A phylogenetic tree made by Clustal Omega summarizing gene sequences encoding rSAM/SPASM XyeB proteins associated with a XyeA precursor peptides (types I-IV), DarA, and DynE. The table summarizes the genera of source bacteria, representative BGC diagrams with core sequences, and the number of unique core sequences found in each genus.

Previously we identified 56 xenorceptide family precursor peptides^20^ with unique core peptide sequences and now classify them into four types based on the conserved core peptide motifs: type A, Ωx_4_Ωx_2_Ωx_4-5_ (n = 37, n: unique core peptide sequences); type B, Ωx_2_Ωx_2_ (n = 2); type C, Ωx_3_Ωx_2_ (n = 2); and type D, Ωx_4_Ωx_2_ (n = 15) where Ω stands for the aromatic amino acid residues Trp, Phe, or Tyr (Fig. 1b). Types A, B, C, and D precursor peptides, lead to xenorceptides Ax, Bx, Cx, and Dx (x: sequential number given in the order of production), respectively. The core peptide sequences, source bacteria, and representative BGCs are shown in Fig. 1c. The phylogenetic tree was constructed based on gene sequences encoding the radical SAM enzyme in the respective BGCs. The 5 predominant genera that encode type A precursor peptides are *Yersinia*, *Xenorhabdus*, *Serratia*, *Erwinia*, and *Photorhabdus*, all of which belong to the order *Enterobacterales*. Type A xenorceptides were predicted to contain three cyclophane rings, and a conserved C-terminal 5-mer (W-T/S-K/R-S-F) comparable to darobactin A (Fig. 1c). While the discovery of darobactin A and dynobactin A relied on production from host bacteria, identifying antibacterial xenorceptides required an on-demand approach to screen natural products for activity. Therefore, we turned toward synthetic biology for production.

### Synthetic biology facilitates production of the xenorceptide natural product family

To produce xenorceptides systematically, we used two different strategies with *E. coli* as a host. In strategy 1, Vector 1 contains N-His_6_ precursor peptide (A) paired with its cognate rSAM enzyme (B), and Vector 2 contains *xncCDE*, the machinery for cleavage and transport from the *xnc* BGC (Fig. 2a). We found that the export proteins XncCDE are promiscuous and carry out more efficient cleavage and transport than those from native BGCs. As an example of strategy 1, we produced **2** encoded by the *smc* BGC from a clinical isolate of *Serratia marcescens* (Supplementary Figure S1). After expression of His_6_-smcAB, Ni-affinity purification and trypsin digestion, we detected a double-charged fragment at *m/z* 1389.7, corresponding to -6 Da mass loss from the C-terminal region of SmcA (ALAQSMLDSVSGGWVNAFARWSKSF, *m/z* 903.8 [M+3H]^3+^). This showed that the rSAM cyclophane synthase SmcB efficiently modified the core peptide as expected. Next, we co-expressed Vector 1 (His_6_-SmcAB) with Vector 2 (XncCDE). This combination drastically increased the yield of **2** compared to the native His_6_-SmcAB + SmcCDE (Supplementary Figure S1). Using strategy 1, we tested 10 AB constructs with XncCDE, and detected the production of 6 end products, and obtained xenorceptides A2–A4 (**2**– **4**) in sufficient amounts for antibacterial assays (Fig. 2b and Supplementary Fig. S2).

**Figure 2.**
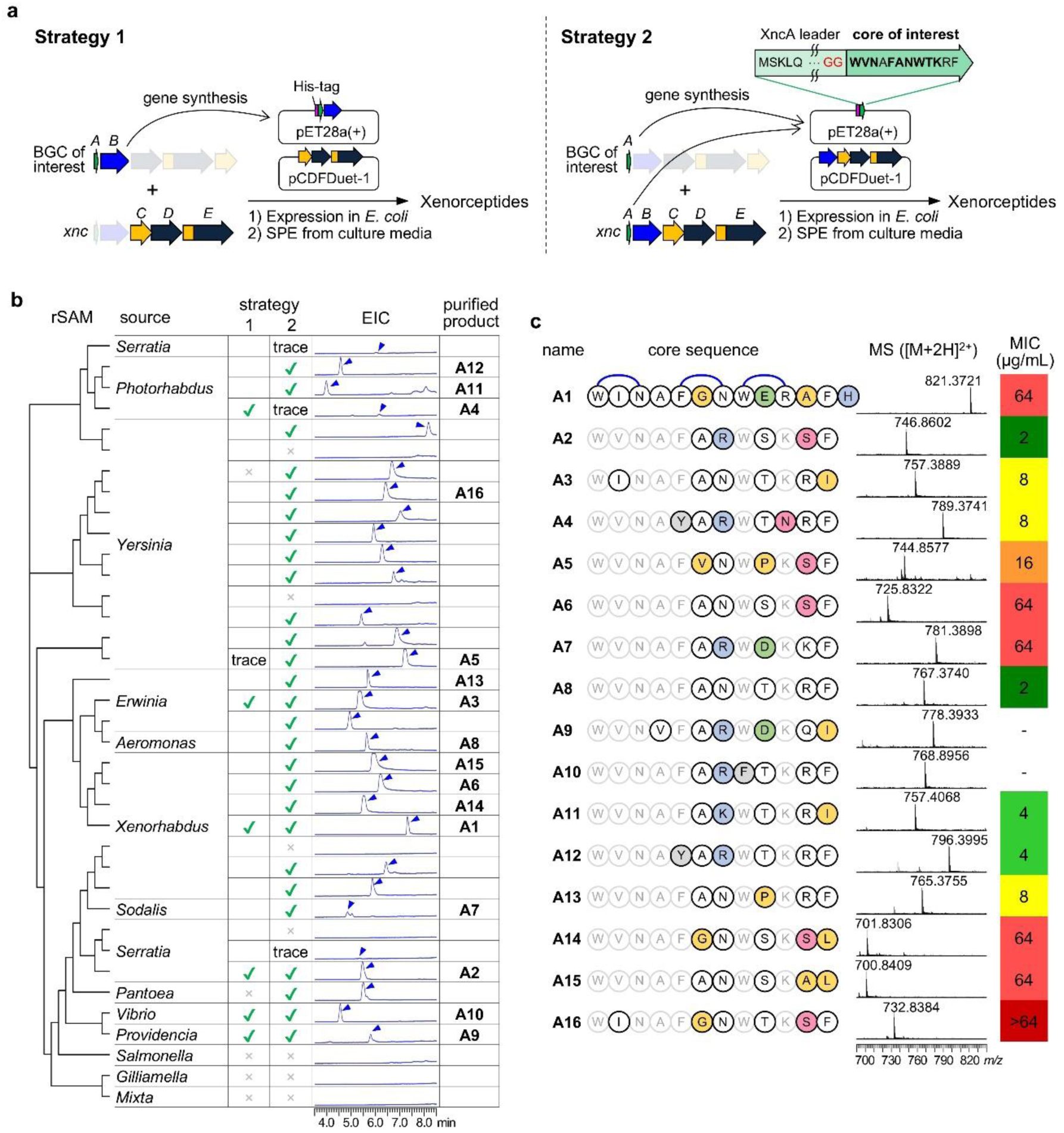
Production and antibacterial activity for xenorceptides. (a) Production of natural products using 2-vector systems. Strategy 1 employs His6-AB/pET28a(+) and CDE/pCDFDuet-1 while in strategy 2 the precursor constituted of His-tagged XncA leader and Xye triceptide core sequence is encoded in pET28a(+) and co-expressed with XncBCDE/pCDFDuet-1. (b) Production of Type A xenorceptides with 37 unique core sequences in two strategies. The phylogenetic tree made by Clustal Omega summarizes gene sequences encoding rSAM/SPASM XyeB proteins associated with a XyeA precursor peptide with a unique core. Check marks represent the strategies successfully producing the natural products. Extracted ion chromatograms (EICs) detect natural products whose *m/z* values correspond to -6 Da mass loss from those of unmodified core peptides. More detailed information can be found in Extended Data Fig. 1. (c) Summary of xenorceptides A1–A16 (**1**–**16**) purified in this study. Blue connectors indicate the residues forming a cyclophane ring while characteristic residues are highlighted in colors. HR-MS data and minimal inhibitory concentrations (MICs) against *E. coli* ATCC 25922 are shown. Products A9 and A10 could not be tested in the subsequent bioassays due to the low yield.

For strategy 2, the precursor peptide genes encoding His_6_-XncA leader fused to the desired core peptides were synthesized and inserted into pET28a(+) vector (Vector 1). These constructs were prepared for all 37 unique core peptide sequences found in 13 genera. The chimeric precursor was co-expressed with XncBCDE encoded in pCDFDuet-1 (Vector 2), and the end products were obtained from the culture media (Fig. 2a). This strategy allowed us to detect 30 end products corresponding to -6 Da loss from the *m/z* values of respective core peptide sequences (Fig. 2b and Extended Data Fig. 1). Tandem MS analysis confirmed the amino acid sequences and -2 Da losses localized to each of the three expected motifs (Supplementary Fig. S3–S7 and Table S1). These results demonstrate that the *xnc* BGC chassis is applicable to produce a broad range of type A xenorceptides in *E. coli*. We selected 12 constructs encoding characteristic core peptide sequences and unique bacterial genera for larger scale fermentation (Fig. 2c). These natural products were named xenorceptides A5–A16 (**5**–**16**) in the order of isolation and were subjected to minimal inhibitory concentration (MIC) assays for antibacterial activity assessments.

### Xenorceptides target Enterobacteriaceae and overcome antimicrobial resistance

Xenorceptides A1–A16 (**1**–**16**) except **9** and **10** along with selected synthetic versions of the unmodified peptide sequences were screened for antibacterial activity. Our initial panel consisted of quality control strains representing Gram-positive and Gram-negative bacteria. Minimal inhibitory concentration (MIC) values were obtained using broth microdilution assays (Fig. 2c and Extended Data Table 1). While **1** showed weak or no activity as before^21^, **2**-**4**, **8**, and **11**-**13** showed selective activity against Gram-negative pathogens (*E. coli* ATCC 25922 and *K. pneumoniae* ATCC 700603). All tested products were not active against Gram-positive bacteria (*B. subtilis* ATCC 6633 and *S. aureus* ATCC 29737) proving selective activity against Gram-negative bacteria. The unmodified synthetic peptides representing the core peptide sequences from **2**-**4** did not show any activity against neither Gram-negative nor Gram-positive bacteria, which confirms that the post-translational modification is critical for the antibacterial activity. We shifted our focus to further biological evaluation and mode of action studies on the most potent analog, xenorceptide A2 (**2**). **2** was tested against a larger panel of clinical drug-resistant isolates. These results are summarized in Table 1 and confirm the selective activity (2–8 μg/ml MICs) against Gram-negative Enterobacteriaceae, several of which are CRE pathogens.

**Table 1.** MIC values for 2.

| Species | Strain | MIC<br>($\mu\text{g/mL}$) |
| --- | --- | --- |
| **Gram-negative bacteria** |  |  |
| *Escherichia coli* | M2 | 16 |
|  | M6 | 8 |
|  | M10 | 4 |
|  | M11 | 4 |
|  | CRE1006 | 4 |
|  | ATCC 25922<sup>a</sup> | 2 |
|  | *bamA*<sup>G807D</sup> | 2 |
|  | *bamA*<sup>G807R</sup> | 8 |
|  | MG1655<sup>b</sup> | 2 |
|  | strain 1 | 8 |
|  | strain 2 | 8 |
|  | *bamA*<sup>G429C</sup> | 32 |
| *Klebsiella pneumoniae* | M7 | 64 |
|  | CRE1007 | 8 |
|  | CRE1008 | 8 |
|  | CRE1011 | 8 |
|  | CRE1012 | 8 |
|  | ATCC 700603 | 8 |
| *Enterobacter cloacae* | CRE1010 | 4 |
|  | CRE1014 | 16 |
|  | CRE1015 | 32 |
|  | CRE1016 | 16 |
|  | CRE1017 | 32 |
| *Salmonella typhimurium* | ATCC 14028 | 8 |
| *Salmonella enteritidis* | ATCC 13076 | 8 |
| *Shigella flexneri* | M90T | 2 |
| *Acinetobacter baumannii* | ACBA1001 | 32 |
|  | ACBA1002 | 32 |
|  | ACBA1003 | 32 |
|  | ACBA1004 | 64 |
|  | ATCC 19606 | >64 |
| *Pseudomonas aeruginosa* | DR4877/07 | 64 |
|  | DR5790/07 | 64 |
|  | DM4150R | 64 |
|  | DM23376 | >64 |
|  | ATCC 9027 | 64 |
| *Morganella morganii* | CRE1001 | 32 |
|  | ATCC 25830 | 32 |
| **Gram-positive bacteria** |  |  |
| *Staphylococcus aureus* | ATCC 29737 | >64 |
|  | ATCC 43300 | >64 |
| *Bacillus cereus* | ATCC 11778 | >64 |
| *Bacillus subtilis* | ATCC 6633 | >64 |

<sup>a</sup>darobactin- or dynobactin-resistant *bamA* mutants previously reported.168 <sup>b</sup>darobactin-resistant strains previously reported. *bamA*<sup>G429C</sup> was generated in the present study.

### NMR structural analysis of 2 shows unique cyclophane topology

Cyclization of **2** was characterized by NMR spectroscopy to understand whether the cyclophane synthase SmcB from *S. marcescens* catalyzes similar cyclophane topology compared to XncB from *X. nematophila*. Structural analysis was carried out similar to **1**, and the NMR spectra and data are shown in Extended Data Fig. 2, Supplementary Fig. S8–S12 and Table S2. The first (WVN) and third (WSK) Trp-containing cyclophanes were identified to contain a crosslink between Trp-C6 to Cβ and are thus analogous to **1**. The FAR cyclophane in **2** was elucidated as *meta*-substituted based on 2D NMR. Phe5-H2 was assigned to a singlet (δ 6.91 ppm) while the remaining three aromatic protons (H4, δ 7.17 ppm; H5, δ 7.25 ppm; H6, δ 7.09 ppm) were within the same spin system. The metacyclophane in **2** has not been observed in a peptide natural product, but bears some resemblance to those found in plant-derived burpitide family RiPPs.^22^ The relative and absolute configurations of the cyclophane rings were assigned by NMR and Marfey’s analysis (Extended Data Fig. 2 and Supplementary Table S3).^23^ This showed the planar or axial chiralities of the WVN, FAR, and WSK cyclophanes are ^3^*M,* ^1^*P*, and ^3^*M* respectively.

### Cell-wide exploration for the mode of action of 2 highlights outer membrane involvement

The chemical structure of **2** shares similar structural features to darobactin A and dynobactin A, therefore we reasoned that **2** targets BamA’s lateral gate. We employed Thermal Proteome Profiling (TPP)^24^ in a high-throughput format, known as Protein Integral Solubility Alteration (PISA) assay^25^. PISA was performed on lysates (for both dynobactin A and **2**) and on whole cells (for **2** only) of *E. coli* K12 MG1655. The lysate experiment revealed significant solubility enhancement (cutoff: p-value = 0.05, arbitrary, log2FC = 0.8) in outer membrane proteins and associated lipoproteins upon exposure to dynobactin A and **2** (Fig. 3a and 3b and Extended data Fig. 3a), indicating a commonality of 68% across the identified hits on the lysate datasets. The high commonality in identified hits across the lysate datasets provided strong converging evidence on a similar mode of action for both ligands, suggesting a targeted disruption of outer membrane as a result of drug binding. Moreover, a pair-wise analysis of cell and lysate experiment done for **2** suggested BamA as a significant hit among proteins with altered solubility (Extended data Fig. 3b, c). Additionally, a growth inhibition assay was conducted to compare susceptibility of wild-type *E. coli* with a *ΔbamB* knockout. The knockout cells exhibited increased sensitivity to both dynobactin A and **2** when exposed to sub-MIC concentrations, mirroring an effect previously reported for the BamA-targeting small molecule MRL-494^26^ (Fig. 3c, Extended data Fig. 4). The increased sensitivity in *ΔbamB* cells further supported that BamA is a likely target for **2**, as the absence of BamB may enhance compound access to BamA, amplifying its antimicrobial effect.

**Figure 3.**
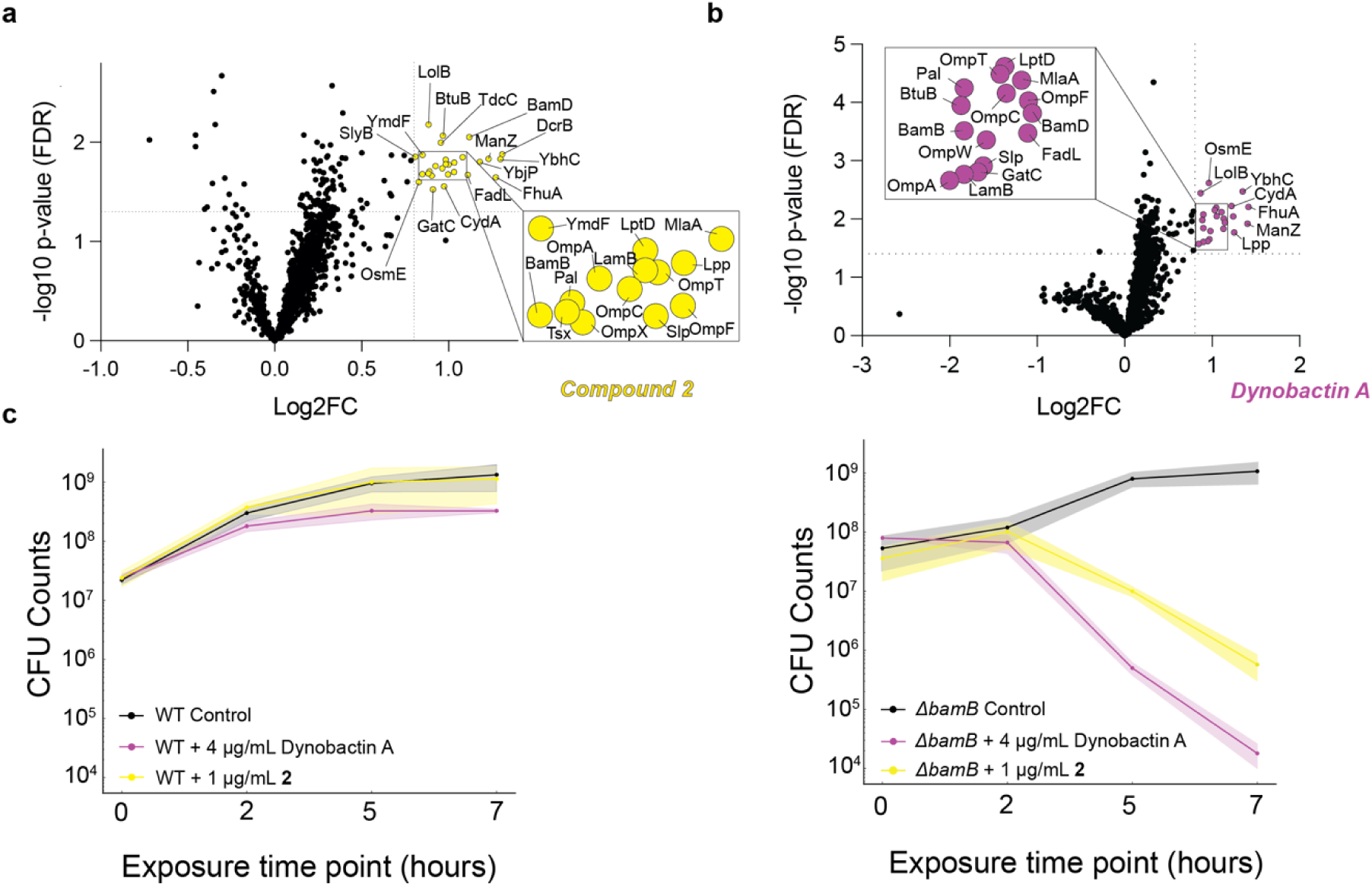
Cell-wide target deconvolution of dynobactin A and 2. (a) PISA assay performed on *E. coli* K12 MG1655 lysates with **2** and (b) dynobactin A, significant solubility alterations observed in outer membrane proteins and outer membrane-associated lipoproteins, including BamD and BamB (number of detected unique peptides > 5). (c) Effect of *bamB* deletion on CFUs (Colony Forming Units) of *E. coli* K12 MG1655 in the growth inhibition assay, demonstrating increased susceptibility of *ΔbamB* cells to either of dynobactin A or **2**.

**Figure 4.**
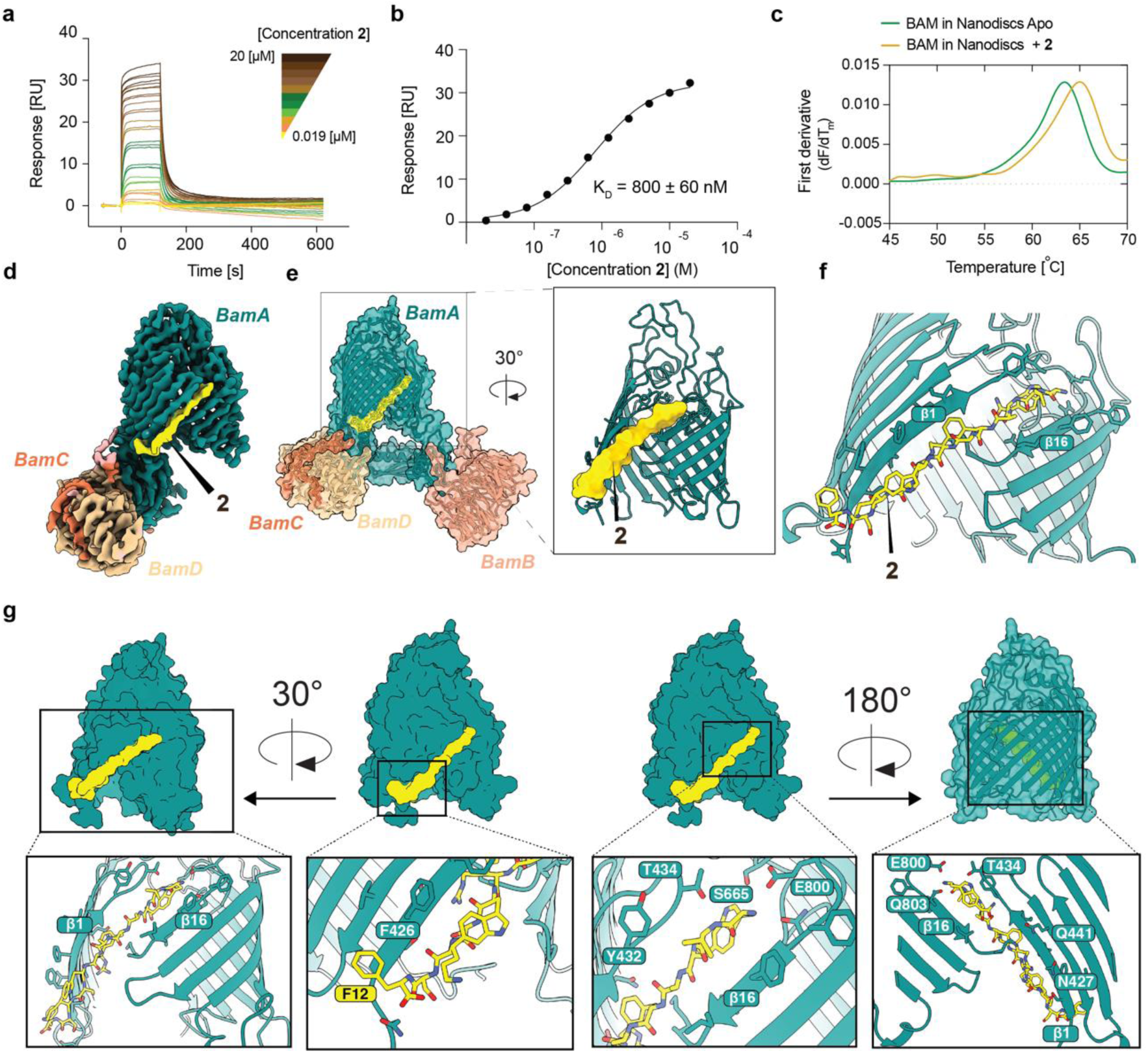
Biophysical Characterization and Structure Determination of the BAM-compound 2 Complex. (a) Sensorgrams for affinity measurements of **2** to BamA-β via Surface Plasmon Resonance and (b) the corresponding steady-state affinity plots show **2** binding to BamA-β within the concentration range of 0.019 µM to 20 µM, with the dissociation constant (KD ≈ 0.8 µM) calculated from the steady state response. (c) DSF of the full-length BAM complex in nanodiscs, illustrating that **2** binding increases the melting temperature of the complex, indicating drug-induced protein stabilization. (d) Cryo-EM map of BAM in MSPdH5 nanodiscs, with the density from **2** colored in yellow. (e) Three-dimensional structural model resolved from the 3.0 Å map of BAM complex (POTRA domain is not shown) obtained in MSPdH5 nanodiscs bound to **2**, showing **2** binds to BAM by introducing a new β strand in the transmembrane region of BamA. (f) Close-up view of the **2** binding site. (g) Binding pocket details where in top BamA is shown as surface and zoomed-in panels with BamA shown as ribbon detailing the BamA-**2** binding interface.

### Compound 2 unveils a novel mechanism of action via BamA

We followed up by measuring affinity of **2** to BamA-β (transmembrane β-barrel domain of BamA) in lauryl dimethylamine oxide (LDAO) micelles using Surface Plasmon Resonance (SPR). The dissociation constant K_D_ was determined from steady-state measurements to K_D_ = 800 ± 60 nM and was consistent with the value derived from kinetics analysis (K_D_ = 676 nM) (Fig. 4a and 4b, Extended data Fig. 5). Next, to test binding in a native-like environment, we reconstituted BAM complex in MSP1D1 nanodiscs (DMPC 14:0 lipids). Differential Scanning Fluorimetry (DSF) was applied in the absence and presence of **2**, demonstrating a 3.8 °C increase in melting temperature (T_m_) of BAM complex upon drug binding (Fig. 4c). Accordingly, BamA was validated as the target for **2**, evidenced by its strong binding affinity and thermal stabilization of the complex. To resolve interaction of **2** to BamA at the atomic level, we employed cryo-electron microscopy (cryo-EM) on purified BAM complex reconstituted in three different membrane mimetics and incubated with excess of **2**. The 3D reconstruction of BAM with **2** in DDM micelles revealed that BAM adopted the outward-closed conformation, but the resolution was insufficient to build an atomic model (Extended data Fig. 6c). For BAM with **2** reconstituted in MSP1D1 nanodiscs, the overall resolution was better, but the lateral gate of BamA could still not be resolved (Extended data Fig. 6d). For BAM in MSPdH5 nanodiscs, however, a map at 3.0 Å resolution was successfully reconstructed, and this map featured density for **2** that was clearly detectable and interpretable with an all-atom model (Fig. 4d, Extended data Fig. 6f). Compound **2** targeted BamA at the lateral gate, as expected, but strikingly, it formed a binding interface that was fundamentally different from the previously characterized bamabactins. Structural analysis revealed that **2** binds to BamA by inserting itself as an extra β-strand between strands β1 and β16 in the transmembrane region (Fig. 4d-g).

**Figure 5.**
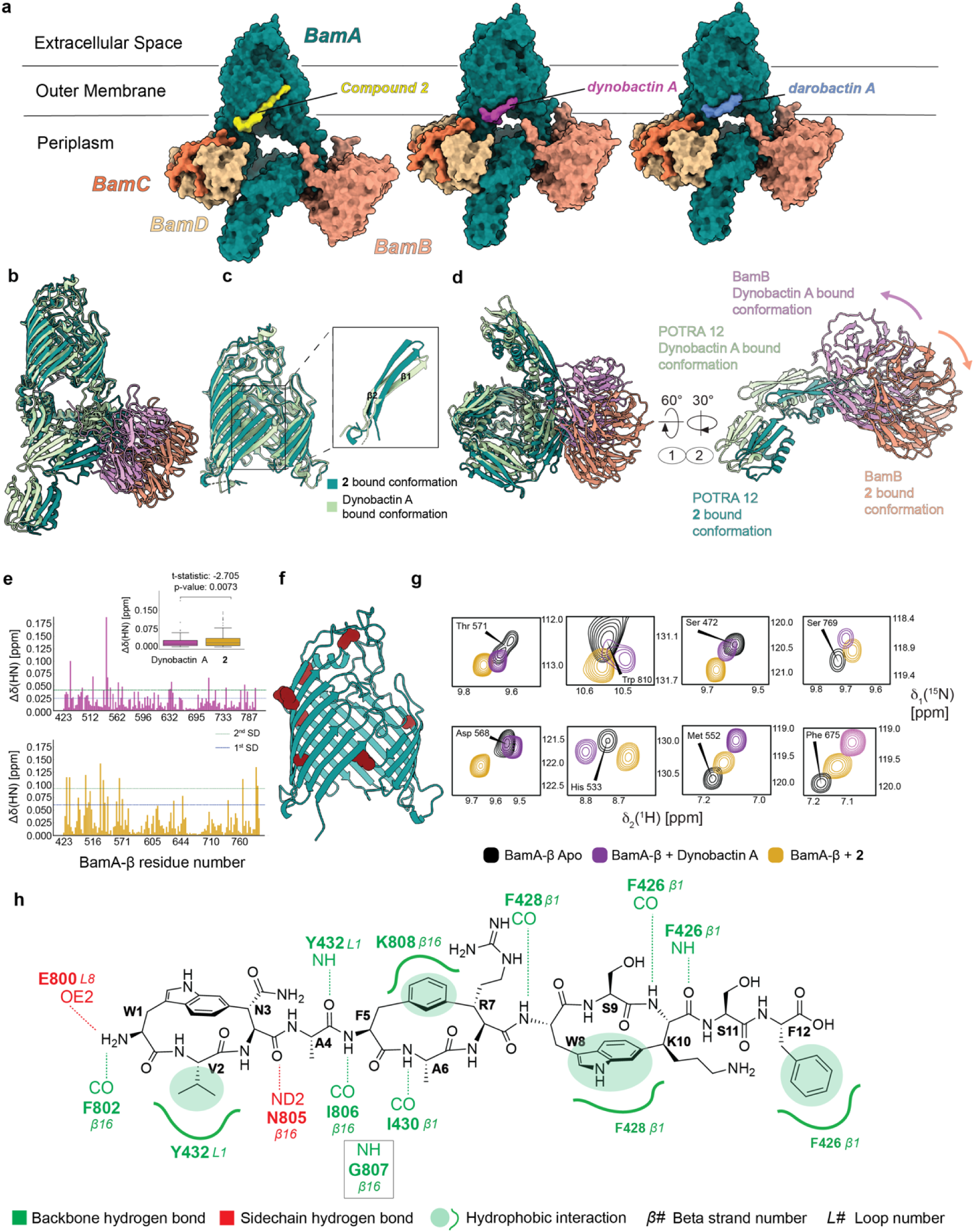
Atomic Details for Mode of Action of compound 2. (a) Structural comparison of BAM-targeting natural peptides (bamabactins). (b) Overlaid structures of dynobactin A bound BAM complex (PDB:7R1W) and **2** bound (PDB: **9HE1**) and (c) Conformation of the lateral gate from cryo-EM structural models, β-strands β1 and β2 of BamA tilts counterclockwise to accommodate **2**. (d) Bottom and side view of the same structural comparison showing BamB and BamA’s POTRA 1 and 2 conformational differences when bound to dynobactin A or **2**. (e) Chemical Shift Perturbation plots for [15N, 1H]-TROSY spectra of BamA induced by addition of two equivalents of **2** (yellow) and dynobactin A (purple) as well as a t-test performed on average CSPs from two datasets indicating a significant variation comparing the two datasets. Blue and green dashed lines on the plot, represent the first and second standard deviation. (f) Representative residues from the TROSY spectra of BamA-β corresponding to the (g) Assigned [15N, 1H] amide NMR resonances of BamA-β in LDAO detergent micelles, obtained from a TROSY experiment conducted in solution. NMR data suggest presence of two different conformations of BamA-β in solution, each induced by interaction with either **2** or dynobactin A. (h) Contact map illustrating intermolecular interactions between **2** and BamA.

**Figure 6.**
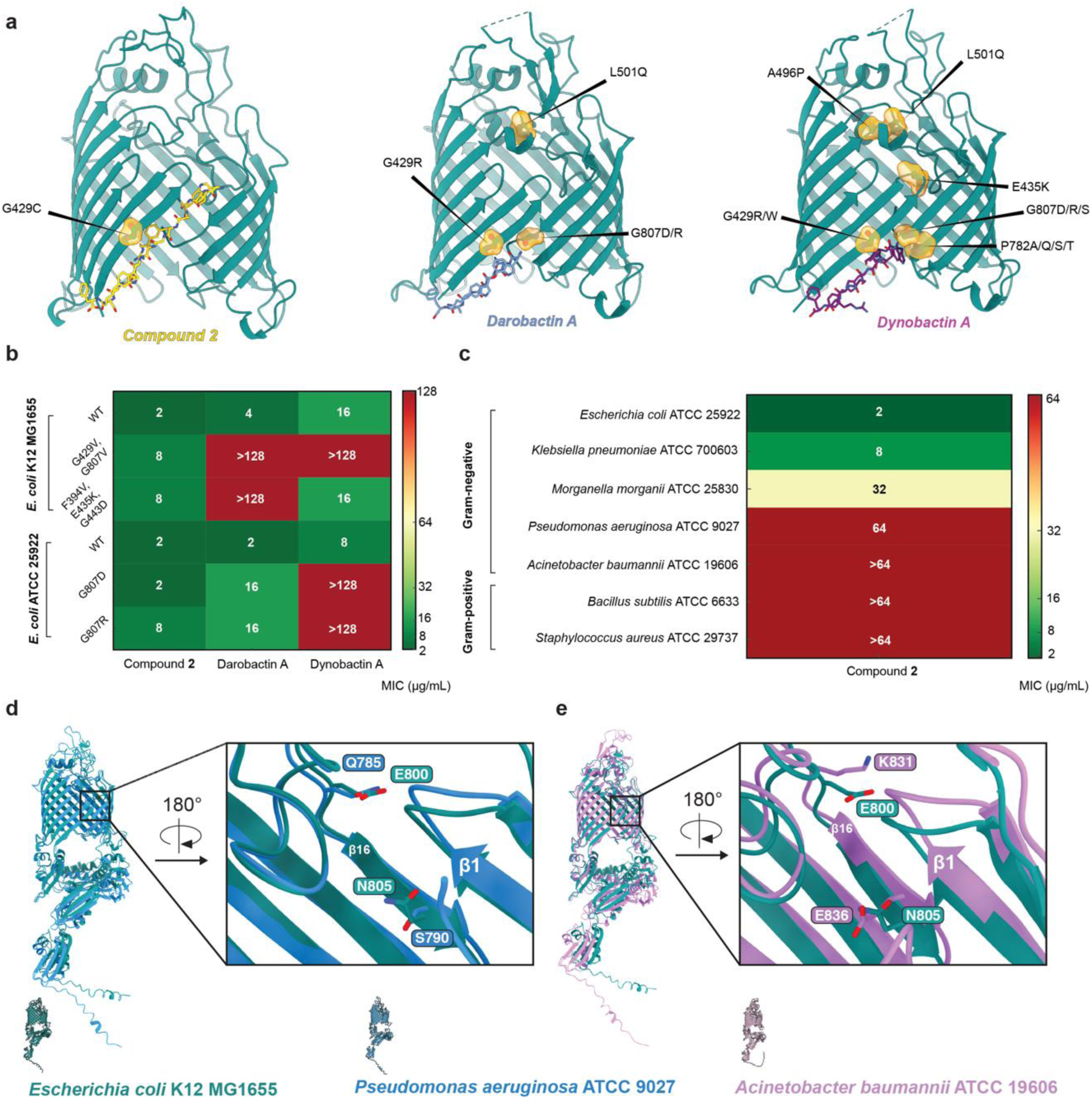
Structure activity relationship and the effect of BamA mutantions on the activity of bamabactins. (a) Transmembrane domain of BamA is shown as a cartoon representation and colored in teal. Mutated residues associated with resistance (MIC > 16 µg.ml-1) are shown as sticks along their corresponding surface view (orange) for BamA in complex with compound **2**, darobactin A (PDB: 7NRF) and dynobactin A (PDB: 7R1V), respectively, with bound molecules also depicted as sticks. (b) MIC values of bamabactins targeting BamA of *E.coli* K12 MG1655 and *E. coli* ATCC 25922 strains showed as heatmap. Compound **2** remains active against *E. coli* strains with mutations conferring resistance to darobactin A and dynobactin A. (c) MIC values of compound **2** against a panel of Gram-negative and Gram-positive bacteria where **2** is active against *Enterobacteriaceae* only. (d) AlphaFold predictions of BamA from *E. coli*, *P. aeruginosa*, and *A. baumannii*, highlighting structural variations that explain the inactivity of compound **2** on these pathogens. In *P. aeruginosa*, the N805 residue in *E. coli* is replaced by a shortened side chain (S790), and in *A. baumannii*, the E800 residue in *E. coli* undergoes charge inversion (K831), disrupting key interactions of **2** with BamA. These alterations explain the loss of activity of compound **2** against *P. aeruginosa* and *A. baumannii*.

Compared to canonical β-barrels, BamA is characterized by its lateral gate, where hydrogen bonds remain incompletely formed. Darobactin and dynobactin interact with BamA through a β-augmentation mechanism, effectively filling the open space at the bottom of the β16 strand. This interaction thus closes the lateral gate by extending the existing hydrogen-bond network and stabilizing the barrel structure^9^. In contrast, integration of **2** into BamA forms an additional β-strand, thus converting the β-barrel structure of BamA from a 16-stranded into a 17-stranded hybrid barrel. This unique interaction stabilizes the lateral gate not just by filling gaps in the hydrogen-bonding network, but by significantly enlarging the structural interface, resulting in a tighter closure of the BamA-β (Fig 5a, Extended data Fig.7). Analysis of the BamA lateral gate structure revealed that β1 and β2 tilt counterclockwise to accommodate **2**. The periplasmic BamB as well as POTRA 1 and 2 reorient to create a new binding interface. (Fig. 5b-d). This space was previously unexploited by bamabactins and thus highlights both the enormous plasticity of the BAM complex in the gate region, as well as the unique interaction mechanism of our new compound.

**Figure 7.**
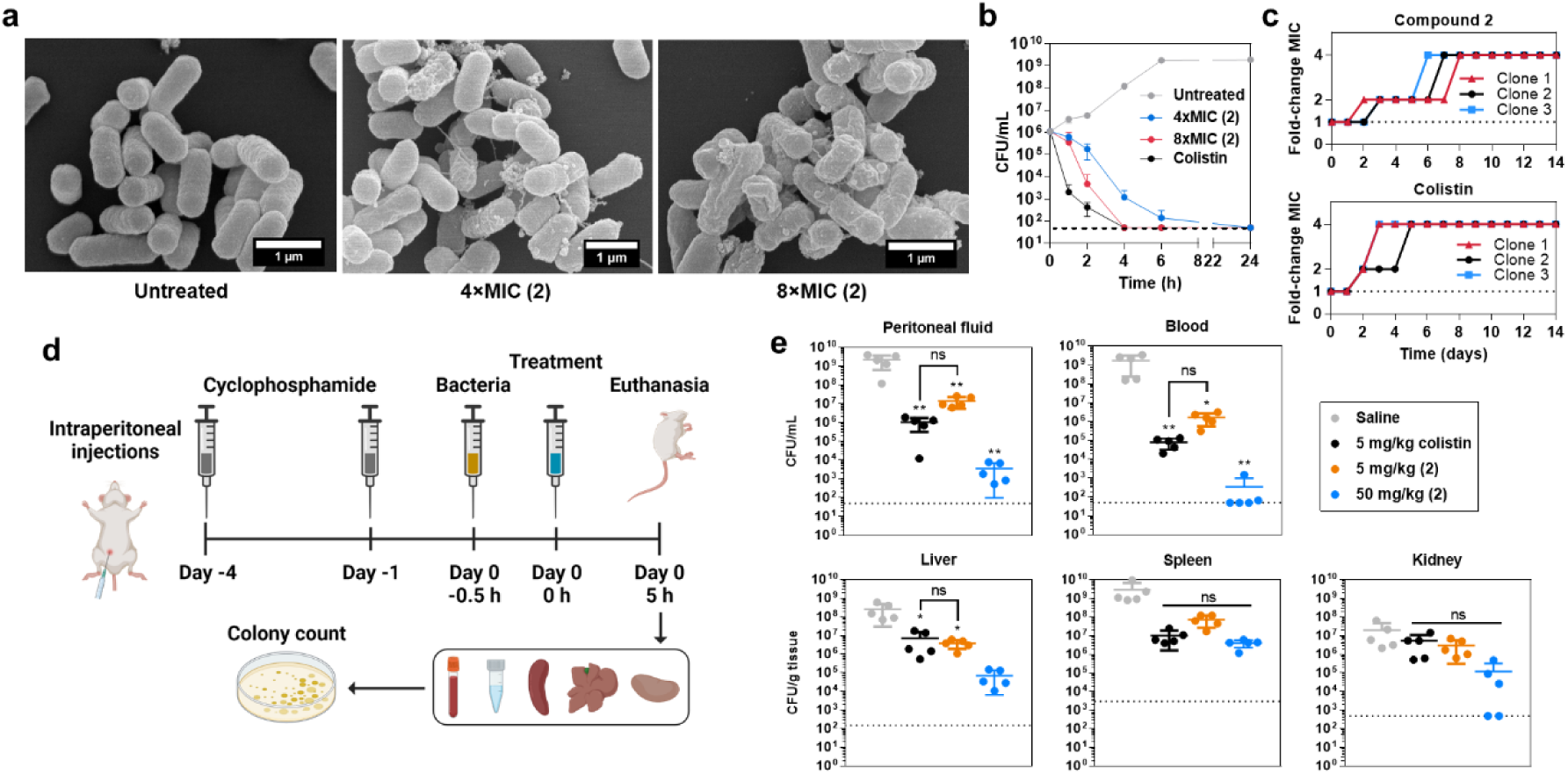
Biological evaluation of compound 2. (a) SEM images of *E. coli* M6 after treatment with **2** at 4× or 8×MIC for 2 h. For each sample slide, at least five independent fields were imaged to ensure representativeness. Magnification = 25,000×. Scale bar = 1 μm. (b) Time-kill kinetics of **2** against *E. coli* M6 over 24 h was determined by agar colony count. Colistin at 2×MIC was tested as a positive control. Black dotted lines indicate the limit of detection (50 CFU/mL). Experiments were repeated on three biologically independent samples. Data are presented as geometric mean ± SE. (c) The development of resistance of *E. coli* M6 against **2** was monitored using serial passage over 14 days. Experiments were repeated three times with different starting bacterial cultures. (d) Experimental schematics of the mouse peritonitis model infected with *E. coli* M6 for evaluating the *in vivo* efficacy of **2**. (**e**) Bacterial burden in the peritoneal fluid, blood, liver, spleen, and kidney of C57BL/6NTac mice (n = 5 mice per treatment group) collected 5 h after treatment with 5 mg/kg **2**, 50 mg/kg **2**, 5 mg/kg colistin, or saline (vehicle control). Samples were plated onto LB agar and incubated for 18-20 h at 37 °C before colony count. Colony counts of organ tissues were normalized against the average mass of the respective mouse organs. Statistical significance of differences between data groups were evaluated using one-way analysis of variance (ANOVA) followed by Turkey post-hoc test (ns: p > 0.05, *: p ≤ 0.05, **: p ≤ 0.01).

We then assessed the conformation of BamA-β in solution with NMR spectroscopy by recording 2D [^15^N,^1^H]-TROSY experiments on spin labelled BamA-β in LDAO micelles. Addition of two equivalents of either dynobactin or **2** induced large chemical shift perturbations (Extended data Fig. 9). Evidently, binding of **2** resolved the two-state conformational equilibrium that is present in apo BamA-β, into a single conformation, just as the other bamabactins. However, the resulting single set of amide resonances was detected at substantially different positions than for dynobactin, demonstrating that **2** induces a particular conformation in solution of BamA-β that is not observed for the other bamabactins (Fig. 5e-g). The behavior of isolated BamA-β thus correlates well with the structural rearrangements observed in the cryo-EM models.

Interaction of **2** with BamA is established through multiple intermolecular contacts including hydrophobic interactions and hydrogen bonds while the dominant series of interactions are the canonical backbone-backbone hydrogen bonds with strands β1 and β16 of BamA. Backbone amine of F5 forms a new hydrogen bond with the carbonyl of I806 on BamA. The methylene moieties of lysine 808 located on β1 of BamA participate in a hydrophobic interaction with F5 on **2**, establishing a novel binding interface. Interestingly, F5 is positioned near G807 on β16 of BamA, suggesting a possible interaction that may explain insensitivity of **2** to mutations at this site (Fig. 5h, Fig 7b, Extended Data Fig. 8). The uncovered targetable binding site of BamA involves a hydrogen bond between backbone amine of Y432 located on loop 1 of BamA with carbonyl of A4 on **2** as well as a hydrophobic interaction between the same tyrosine with V2 on **2**. Additionally, terminal amine of **2** forms a non-canonical bifurcated hydrogen bond with backbone carbonyl of F802 (β16) and Oε side chain of E800 (β16), simultaneously, thus further strengthening β-augmentation mechanism for **2** to be integrated to the BamA β-barrel as an extra β-strand (Fig. 5h, Extended Data Fig. 8).

### Resistance evolution and cross-resistance to bamabactins

To study resistance patterns against **2**, we monitored the *bamA* mutations emerging from a resistance-experiment against **2**. The *E. coli* MG1655 cells at mid-log phase were streaked onto Muller-Hinton II B agar containing **2** at 2x MIC (4 μg/mL). The resistant colonies were further screened in the MHIIB liquid broth containing different concentrations of **2** to determine MICs. 20 strains with a range of resistance were selected for PCR amplification and sequencing of the *bamA* locus. Unexpectedly, 18 strains with lower resistance (2x ∼ 8x MIC) had no detectable mutation on *bamA.* In contrast, two strains with the highest resistance (16x MIC) showed the same single amino acid mutation: *bamA* G429C (Fig. 6a). Genomic recombination of wild-type MG1655 by this *bamA^G429C^* gene fragment provided 16x higher resistance to **2**, as the original resistant strains (Fig. 6a and Table 1). This allelic mutation experiment revealed the single *bamA* mutation completely accounted for the change in activity. Next, we tested efficacy of **2** against reported *E. coli* strains harboring BamA mutations and resistance to other bamabactins. The BamA mutations in these strains lead to an MIC of >128 μg/mL for darobactin and/or dynobactin^ref^. Surprisingly, **2** was effective against the four resistant strains (MIC = 2 or 8 μg/mL), demonstrating that xenorceptides are not susceptible to cross-resistance against both darobactin and dynobactin (Fig. 7b). Collectively, these results confirmed BamA as the target of **2** but its mutations are not as critical for antibacterial activity of **2** compared to other bamabactins.

### AlphaFold predictions explain narrow spectrum activity of xenorceptides

Given the observed narrow-spectrum of activity for xenorceptides, we compared the interaction site on BamA for susceptible and resistant strains. AlphaFold predictions for the BamA structure in *A. baumannii* revealed a lysine residue (K831) at the position where glutamate (E800) resides in *E. coli* K12, interacting with the N-terminus of **2**. This charge inversion likely disrupts the binding of **2**, and might explain its lack of activity against *A. baumannii* (Fig. 7c and e, Extended data Fig. 10). Furthermore, AlphaFold predictions for the BamA structure in the **2**-resistant strain *P. aeruginosa* revealed a shorter side chain (S790), corresponding to the position of N805 in *E. coli*, where it interacts with the N3 group of **2**. This variation in residues among strains likely reduces the binding affinity of **2** to BamA, contributing to the strain’s resistance (Fig. 7 c and d, Extended data Fig. 10). These findings highlight a novel BamA binding site that can be exploited to design narrow-spectrum antibiotics.

### *In vitro* and *in vivo* biological evaluation of 2

Using scanning electron microscopy (SEM), we further imaged the surface morphology of *E. coli* M6, a carbapenem- and colistin-resistant clinical isolate, in the presence of **2**. Within 2 h of treatment, the cells showed clear membrane damage and surface blebbing, followed by cell lysis and death (Fig. 7a). Time-kill assays against *E. coli* M6 showed that **2** displayed bactericidal effect over 24 h at 8 x MIC, causing 3-log reduction in bacteria count (Fig. 7b). In addition, **2** demonstrated no detectable cytotoxicity against human cell lines (HepG2, HK-2, IMR90, and HIEC-6) up to a concentration of 256 μg/ml, highlighting its superiority over colistin. We then performed serial passage assay where *E. coli* M6 was continually exposed to **2** at sub-inhibitory concentrations to evaluate resistance development compared to colistin. After 5-7 days, *E. coli* M6 developed approximately 4-fold resistance to **2** with an MIC of 32 µg/ml, whereas a faster development of resistance to colistin was observed after 3 days (Fig. 7c). *In vivo* antimicrobial efficacy of **2** was next assessed in mouse models. A single-dose pharmacokinetic study revealed that a 50 mg/kg intraperitoneal injection of **2** resulted in good systemic exposure, reaching a peak blood concentration of 61 μg/mL and a half-life of 2.4 hours (Extended Data Fig. 11a). Importantly, blood concentrations remained above the MIC for *E. coli* for 8.8 hours, indicating strong potential for anti-infective efficacy. **2** was next assessed using a peritonitis model in neutropenic mice (Fig. 7d). After 30 min of injection with *E. coli* M6, mice (n = 5 per group) were given a single intraperitoneal injection of treatment or saline. At 5 h post-treatment, the mice were euthanized for collection of peritoneal fluid, blood, and organs to quantify the bacterial burden using the colony counting method^27^. **2** displayed a concentration-dependent antimicrobial effect in peritoneal fluid, blood, and liver, where 50 mg/kg dose caused a 6-, 7-, and 4-log decrease in colony count relative to saline control results, respectively (Fig. 7e). While a weaker effect was observed in the spleen and kidney, 50 mg/kg of **2** still achieved a 2-log reduction in bacterial burden. At the same 5 mg/kg dose, **2** displayed comparable efficacy to colistin. These experiments show that **2** is a suitable antibiotic lead for further development against Enterobacteriaceae, offering the potential to overcome the toxicity and resistance challenges associated with carbapenems and colistin.

## Discussion

The discovery of antibiotics with novel modes of action is an inevitable need, particularly in response to the growing resistance against clinically used antibiotics.^28^ While natural products have historically been a rich source of antibiotics, few new compounds with alternative modes of action have emerged since the golden era.^7^ Bamabactins are a class of promising lead antibiotics that have a selective activity profile and unique mode of action. The key structural features in these antibiotics are 3- and 4-residues cyclophanes installed by radical SAM enzymes that constrain the peptide into a β-strand conformation. Phylogenetic genome-mining of the radical SAM enzymes led to the identification of xenorceptides as potential Gram-negative targeting antibiotics. Although the chemical synthesis of previously identified bamabactins have been accomplished,^29,30^ the synthesis of all xenorceptides would be a formidable challenge. From another perspective, while genome mining and synthetic biology have enabled phylogenetic approaches to drive further progress on natural products, discovery of selective antibiotics against Gram-negative bacteria with minimal cytotoxicity against human cells remains a significant challenge and requires further investigation.^31^ We addressed these challenges by developing strategies to access the majority of genetically encoded xenorceptides from various bacterial hosts, eliminating the need to treat each biosynthetic gene cluster (BGC) individually, which typically requires specific manipulation for heterologous expression or pathway activation in the host strains. Our discovery revealed that, unlike many antimicrobial peptides that cause non-specific membrane disruption, xenorceptides, the newest members of the bamabactin antibiotic class - selectively targets the essential protein BamA, reducing the off-target effects commonly observed with non-specific antibiotics.^32^ When assessing the mechanistic insights learned from the three types of bamabactins, we realize that cyclization of side chains is present on all of them, but they do not seem to be conserved neither in position nor in chemical type. It rather seems that their main purpose is the stabilization of the peptide in a rigid β-strand-like structure, with simultaneous formation of a lipid interaction face via the cyclized aromatic side chains^9^. The stabilization of the β-strand significantly reduces the entropic penalty that a flexible peptide would need to outcompete the nascent OMP chain when binding to the β-strand edge of β1. We thus conclude that for antibiotic design and development targeting this region, it will be essential to have a similar stabilization in β-strand conformation. From the pool of identified xenorceptides, **2** demonstrates a potent and selective activity against Enterobacteriaceae with sub-micromolar affinity to BamA which could serve as a promising lead compound in the development of novel antibiotics targeting this protein. The lateral gate of BamA presents a unique and highly adaptable target site for antibiotic intervention. Compared to darobactin and dynobactin, xenorceptide has revealed the enormous plasticity of this region. Given the function of BamA, which can accommodate the augmentation of multiple β-strands to β1, it seems plausible that the conformational space of competitive inhibitors targeting this site is still by far not fully exploited. This flexibility in the lateral gate of BamA may enable the design of a broad range of diverse inhibitors, with mutually exclusive resistance patterns, providing a versatile platform for antibiotic development with fine-tuned spectrum of activity by modulating the differences in the N-terminus of this family of compounds. Furthermore, **2**, with its unique lateral gate insertion mechanism, offers enhanced resistance to mutations compared to darobactin and dynobactin, likely due to its deeper integration into the β-barrel structure. This broader inhibition mechanism positions **2** as a promising candidate for targeting BamA, even in strains that have developed resistance to other bamabactins. Additionally, its selective inhibition of BamA in specific pathogens, while sparing others, holds potential for the development of antibiotics with reduced off-target effects and a lower risk of driving resistance in non-target bacterial populations. Biological profiling of **2** demonstrates that this compound presents several key advantages over last resort colistin and the recently identified macolacin.^31^ The absence of detectable cytotoxicity of **2** against human cell lines highlights a critical improvement in safety over colistin, which is known for its nephrotoxic effects.^33^ *In vivo*, **2** achieves significant bacterial clearance in key organs, demonstrating robust efficacy against resistant *Enterobacteriaceae*. Additionally, slower rate of resistance development against **2** compared to colistin underscores **2** as a promising candidate for further development, offering a safer and more effective alternative to colistin, particularly in treating multidrug-resistant Gram-negative pathogens.

## Materials and Methods

### Materials, equipment, and general experimental procedures

Chemicals and reagents were purchased from the following suppliers: Acetonitrile from Tedia (USA); Isopropanol and methanol from Thermo Fisher Scientific (USA); Kanamycin and spectinomycin from GoldBio; Isopropyl β-_D_-1-thiogalactopyranoside (IPTG) from Combi-Blocks; Strata-X^®^ Polymeric Solid Phase Extraction (SPE) Sorbent (33 µm) from Phenomenex (USA); NMR solvent DMSO-*d*_6_ from Cambridge Isotope Labs (USA); and synthetic peptides from GL Biochem (China). Other chemicals and reagents were purchased from either Sigma (USA) or Bio Basic (Canada). Synthetic genes inserted into expression vectors were purchased from Twist Bioscience (USA). *Escherichia coli* NiCo21(DE3) cells were purchased from New England Biolabs (USA). Electroporation was carried out using mode p2 (2.5 kV, 5.6 ms) on a MicroPulser Electroporator (Bio-Rad, USA). Ultrasonication was carried out using an Ultrasonic Cleaner 142-0307 (VWR, USA). Centrifugation was carried out using either an Eppendorf^®^ Centrifuge 5424R or 5810R (Germany), or an Avanti JXN-26 Ultracentrifuge (Beckman Coulter, USA). SPE was performed using either 12-Position Vacuum Manifold Set (Phenomenex, USA) or Vac-Man^®^ Vacuum Manifold (Promega, USA). Sample solutions were concentrated using either a rotary evaporator (Rotavapor^®^ R-210, Büchi, Switzerland), centrifugal evaporator (Genevac EZ-2 Elite, SP Scientific, UK), or freeze dryer (ScanVac CoolSafe, LaboGene, Denmark). LC-MS experiments were performed on a Waters Acquity UPLC System coupled to Xevo G1 QToF Mass Spectrometer (USA) and data was analyzed using MassLynx v.4.1. Preparative HPLC was carried out on a Shimadzu Nexera Prep System. NMR spectra were acquired at 298 K using a Bruker 400 MHz Avance Neo Nanobay NMR Spectrometer (USA) with a Bruker iProbe 5 mm SmartProbe or a Bruker 800 MHz Avance Neo NMR Spectrometer (USA) with a Bruker 5 mm CPTXI Cryoprobe and data was analyzed using Bruker Topspin v3.6.

### Transformation of plasmids into *E. coli* cells

Plasmids containing precursor genes (*xyeA*), rSAM (*xyeB*), peptidase and transporter (*xyeCDE*) genes in varied constructs were synthesized by Twist Bioscience. Synthesized gene sequences are listed in Supplementary Table S4. The plasmids were reconstituted in autoclaved Milli-Q grade 1 water to a final concentration of 10 ng/µL. For full-length gene cluster expression,1 µL of plasmid DNA was added to 70 µL of *E. coli* electrocompetent NiCo21(DE3) cells and transformed in a 2 mm electroporation cuvette. For coexpression, 1 µL of each plasmid DNA containing the appropriate genes was added to 70 µL of *E. coli* electrocompetent cells and transformed in a 2 mm electroporation cuvette. 1 mL of lysogeny broth (LB) was subsequently added to the transformed cells in an Eppendorf tube and incubated in the shaker at 37 °C, 200 rpm for 1 h. Following this, the bacteria cells were centrifuged at 4,000 rpm for 10 min at 25 °C and the cell pellet obtained by disposing the supernatant. The cell pellet was then resuspended with the residual supernatant and streaked on LB agar supplemented with appropriate antibiotics to be grown overnight at 37 °C.

### Expression and purification of His_6_-precursors

An overnight culture of the transformant was inoculated into LB medium in an Ultra Yield^®^ flask (Thomson) at a ratio of 1:100 v/v with appropriate antibiotics. The flask was shaken at 250 rpm and 37 °C until OD_600_ reaches 1.5–3.0. The culture was cooled in an ice bath for 30 min. Protein expression was induced in the presence of 1 mM IPTG at 16 °C and shaken at 250 rpm for 16 to 24 h. The cells harvested by centrifugation were reconstituted in denaturing lysis buffer (100 mM NaH_2_PO_4_, 10 mM Tris, 9 M urea, 10 mM imidazole, pH 8.0) and then lysed by ultrasonication. The His_6_-precursor in the supernatant was captured on HisPur Ni-NTA resin (Thermo Scientific, 625 mL per 20 mL supernatant) and purified according to the instructions provided by the manufacturer. The protein was eluted using NPI-250 (50 mM NaH_2_PO_4_, 300 mM NaCl, 250 mM imidazole, pH 8.0) and the buffer was exchanged into 50 mM Tris–HCl (pH 7.5) using a PD Minitrap G-10 column (GE Healthcare). When XyeAB were expressed, the purified protein was digested by trypsin (10 µg per 1 mL eluate) at 37 °C for 16 h. Digested precursors were analyzed by LC-MS using the following conditions: column = Phenomenex Kinetex XB-C18, 5 µm, 150 x 4.6 mm; mobile phase/gradient = solvent A: H_2_O (+0.1% formic acid, FA), solvent B: CH_3_CN (+0.1% FA), isocratic 4% B for 2 min, followed by a linear gradient to 60% B over 10 min; flow rate = 0.5 mL/min; column temp. = 50 °C. When XyeAB and XyeCDE were coexpressed, the purified protein was directly analyzed by LC-MS using the following conditions: column = Phenomenex Aeris WIDEPORE C4, 3.6 µm, 150 x 4.6 mm; mobile phase/gradient = solvent A: H_2_O (+0.1% formic acid, FA), solvent B: 1:1 CH_3_CN/*i*-PrOH (+0.1% FA), isocratic 4% B for 2 min, followed by a linear gradient to 60% B over 12 min; flow rate = 0.5 mL/min; column temp. = 50 °C. High-resolution MS data of modified peptide products is summarized in Supplementary Table S1.

### Purification of Xenorceptides

Production of xenorceptides in the AB + CDE and A + BCDE strategies was conducted in the same workflow. Transformants harboring two vectors were inoculated into LB media and protein expression was induced by IPTG as described above. After the overnight cultivation at 16 °C, cells were removed by centrifugation at 4,000 rpm for 15 min at 4 °C. 1 L supernatant was combined with 5.5 g of free-standing Strata-X^®^ resin in a 2 L conical flask and shaken at 16 °C, 160 rpm for 1 h to allow binding of the core peptide to the resin. Peptide-bound resin was poured into a column, then washed twice with H_2_O (55 mL), 60% methanol (55 mL), 100% methanol (55 mL), and finally eluted with 60% acetonitrile with 0.1% FA (55 mL). The elution fraction was concentrated in vacuo, reconstituted in 20% acetonitrile with 0.1% FA, and subjected to purification by preparative HPLC at either of the following conditions: Method A: column = Imtakt, Cadenza 5CD-C18, 5 µm, 250 x 20 mm; mobile phase/gradient = solvent A: H_2_O (+0.1% FA), solvent B: CH_3_CN (+0.1% FA), isocratic 5% B for 1 min, followed by a linear gradient to 25% B over 17 min; flow rate = 21.2 mL/min; UV detection = 220 nm; column temp. = room temperature. Method B: column = Kinetex XB-C18, 5 µm, 250 x 21.2 mm; mobile phase/gradient = solvent A: H_2_O (+0.1% FA), solvent B: CH_3_CN (+0.1% FA), isocratic 10% B for 1 min, followed by a linear gradient to 30% B over 30 min; flow rate = 8 mL/min; UV detection = 220 nm; column temp. = room temperature.

### Yields of Xenorceptides

Xenorceptides A1-A4 (**1**-**4**) were obtained using method A as white or pale-yellow powders. Xenorceptides A5-A16 (**5**-**16**) were obtained using method B as white or pale-yellow powders. Yields of the products were 5.0 mg/L (A1), 4.6 mg/L (A2), 1.0 mg/L (A3), 3.3 mg/L (A4), 0.3 mg/L (A5), 0.1 mg/L (A6), 0.3 mg/L (A7), 0.2 mg/L (A8), <0.1 mg/L (A9), <0.1 mg/L (A10), 0.6 mg/L (A11), 0.4 mg/L (A12), 0.3 mg/L (A13), 1.5 mg/L (A14), 3.0 mg/L (A15), 1.0 mg/L (A16) of culture.

### Minimum inhibitory concentration (MIC) determination

MIC screening of the peptides against a panel of ATCC and clinical strains was performed using broth microdilution method^34^. Briefly, peptides stock solutions in DMSO (0.1% TFA) were diluted into Mueller Hinton Broth (MHB), followed by two-fold serial dilution in a 96-well plate. Bacteria culture in mid-log phase was diluted into MHB to yield 1×10^6^ colony-forming units (CFU)/mL. Equal volume of the starting inoculum was added to the peptide samples, then incubated for 18–20 h (37 °C, 120 rpm). OD_600_ of the samples was then measured using Tecan Infinite M200 (TECAN, Männedorf, Switzerland). MIC is defined as the lowest peptide concentration to achieve more than 90% reduction in OD_600_ relative to the drug-free control. The experiments were repeated three times. Colistin-resistant clinical isolates are a kind gift from Dr. Jeanette Koh (National University Hospital, Singapore). Multidrug-resistant clinical isolates are a kind gift from Dr. Lakshminarayanan Rajamani (Singapore Eye Research Institute, Singapore).

### Resistance development studies

An overnight culture of *E. coli* MG1655 was inoculated in 10 mL MHIIB at a ratio of 1:100 v/v and shaken at 180 rpm and 37 °C until OD_600_ reaches 0.4–0.6. The culture was diluted to 1×10^8^ CFU/mL and then 500 μL culture (5×10^7^ CFU) was spread onto 6 plates of MHIIB containing xenorceptide A2 (**2**) at 4 μg/mL and 1.5% agar. Plates were incubated at 37 °C for 24 h to yield approx. 300 colonies resistant to **2**. MICs of these colonies in liquid culture were confirmed in 96-well plates as described above. 20 clones with a range of MIC values (2× to 16× MIC) were selected for PCR amplification of the *bamA* locus using the following primers: bamA_Fw, GCTGGGATGACAGCGGAAC; bamA_Rv, TTCACAGCAGTCTGGATACGAG. The PCR product was purified by Wizard^®^ SV Gel and PCR Clean-Up System (Promega) and directly sent for sequencing with two additional primers: bamA_SQ1, GGTAAATATAGCGCCAGC; bamA_SQ2, TCGCTAATAAACGGCGTC. Sequence data were analyzed using UGENE 39.0 (https://ugene.net/).

### Construction of recombinant *bamA* mutant in *E. coli* MG1655

The 3,209-bp genomic DNA fragment encoding the mutated *bamA* gene (1285G>T) was amplified from the resistant strain #19 that shows 16× MIC than wild-type MG1655 and used for λ Red recombination.^35,36^ 200 μL of an overnight culture of *E. coli* MG1655 harboring pCas^37^ at 30 °C was inoculated in 10 mL LB in a 50 mL tube and shaken at 180 rpm and 30 °C. After 1.5 h. 0.4% of _L_-arabinose was added and the tube was transferred for shaking at 180 rpm and 37 °C for 1 h. Cells were collected by centrifugation, washed twice by 5 mL of ice-cold 10% glycerol, and resuspended in 500 μL of ice-cold 10% glycerol. The electrocompetent cells were split into 50 μL each in 1.5 mL tubes and stored at -80 °C until use. Approx. 400 ng of the purified DNA fragment was used to transform electrocompetent cells of *E. coli* MG1655-pCas. The recovery step was performed in SOC medium for 3 h at 37 °C with shaking. The cells were streaked onto an LB agar plate containing xenorceptide A2 (**2**) at 8 μg/mL and incubated at 37 °C for 24 h. Positive colonies were cultivated in 2 mL of LB at 42 °C overnight for curing of pCas, and then restreaked onto LB agar plates to confirm resistance to **2** and sensitivity to kanamycin. The *bamA* locus was amplified by PCR and the presence of the mutation (1285G>T leading to G429C) was confirmed by sequencing.

### Advanced Marfey’s analysis

100 µg of xenorceptide A2 (**2**) was hydrolyzed in 6 M HCl (1 mL) at 110 °C for 18 h. The hydrolysate was concentrated using a centrifugal evaporator and reconstituted in water (100 µL), followed by addition of 1 M NaHCO_3_ (40 µL) and 1% w/v of *N*α-(2,4-dinitro-5-fluorophenyl)-_L_-valinamide (_L_-FDVA) in acetone (200 µL). The mixture was incubated at 42 °C for 1 h and quenched with 2 M HCl (20 µL). _L_-Amino acid standards were derivatized in the same manner using _L_- and _D_-FDVA. The sample was diluted with CH_3_CN/H_2_O (1:1 v/v) and analyzed by LC-MS using negative ion mode. Retention times of the derivatized samples and standards are summarized in Supplementary Table S3 with detailed LC conditions.

### BAM complex expression and purification

The BAM complex was expressed from a plasmid containing all five E. coli BamA–E genes, with BamE C-terminally linked to a His6-tag and BamB C-terminally linked to a StrepII-tag.11 E. coli BL21(DE3) C43 cells were grown in LB medium in the presence of ampicillin at 37°C until an OD600 of 0.6, then induced with 0.1 mM isopropyl-β-D-1-thiogalactopyranoside (IPTG) and further cultivated over night at 20°C. Cells were resuspended in ice cold Tris-buffered saline (TBS; 50 mM Tris-HCl pH 8.0, 300 mM NaCl), homogenized with a dounce homogenizer, lysed using a microfluidizer and pelleted by ultracentrifugation with a 45 Ti rotor (220,000 g, 2h, 4°C). The BAM complex was extracted from the pellet with TBS containing 3% (v/v) Elugent (Calbiochem). After ultracentrifugation with a 45 Ti rotor (220,000 g, 30 min, 4°C), the supernatant was passed through a 0.22 µm filter and loaded on a Ni Sepharose FF column, washed with 20 CV of TBS containing 0.1% (w/v) N-Lauryldimethylamine N-oxide (LDAO) and eluted with TBS containing 0.1% (w/v) LDAO and 300 mM imidazole. The eluate was directly loaded on a Strep-Tactin XT column (IBA) and eluted with TBS containing 0.05% DDM and 10 mM biotin. The BAM complex was further purified on a 16/600 Superdex 200 column using 20 mM Tris-HCl pH 8.0, 150 mM NaCl, 0.05% (w/v) n-dodecyl β-D-maltopyranoside (DDM), concentrated to 8 mg/mL and flash frozen in liquid nitrogen.

### Expression and purification of Membrane Scaffold Proteins (MSPs)

MSP1D1 and MSPdH5 were prepared with a similar method. The pET28a plasmid encoding MSP1D1 with N-terminal HisTag and cleavage site was transformed into *E. coli* BL21(DE3) cells. Afterwards, an overnight culture was inoculated into 1L of LB in baffle flasks. Cells were grown at 37 °C until O.D_600_ of 0.7-0.8 and then induced with 0.5 mM of IPTG. 1 hour after induction, the temperature was decreased to 28 °C and expression was continued for another 4 hours. The cells were pelleted by centrifugation at 4000 g for 20 minutes at 4 °C. Next, cells were resuspended in resuspension buffer (20 mM Tris-HCl, 500 mM NaCl, pH 8.0). The suspension was supplemented with 5 mg of DNAse (Applichem) per liter of expression and 1% of Triton X-100, volumetrically. The mixture was incubated and shaken at 4 °C for 1 hour. Cells were then sonicated with a Bronson sonifier for 20 minutes on 20% power of 2 seconds “on” followed by 2 seconds “off” pulses. The sonicated sample was centrifuged at 30’000 g for 30 minutes at 4 °C after which the supernatant was filtered by syringe and non-pyrogenic 0.2 µM sterile filters. The filtered sample was then injected to an AKTA machine for Ni-NTA based purification. Two HisTrap column (Cytiva - 2x5 mL) were connected where the columns were firstly, equilibrated with priming buffer (20 mM Tris-HCl, 500 mM NaCl, pH 8.0, 1% TritonX-100). The sample was applied to the column with a flowrate of 1 mL/min. The columns were then washed again with 100 mL of priming buffer. Next, 100 mL of cholate buffer (20 mM Tris-HCl, 500 mM NaCl, pH 8.0, 50 mM sodium cholate) was introduced to the columns. Afterwards, another 100 mL of resuspension buffer was applied to the column to prepare the column for gradient elution. Unspecific bound impurities were washed out from the column by wash buffer (20 mM Tris-HCl, 500 mM NaCl, pH 8.0, 50 mM Imidazole) and the His tagged MSP was eluted by gradient application of elution buffer (20 mM Tris-HCl, 500 mM NaCl, pH 8.0, 500 mM Imidazole). The eluted fractions were quickly exchanged to MSP final buffer (20 mM Tris-HCl, 100 mM NaCl, pH 7.4). The HisTag was cleaved by introduction of TEV protease in 1:100 ratio while dialyzing at 4 °C. The sample was then subjected to a reverse HisTrap to remove the TEV protease and the cleaved HisTag. A final Size Exclusion Chromatography (SEC) was done in SEC buffer (20 mM Napi, 100 mM NaCl, pH 7.0) as the final step. The protein was then aliquoted into 500 μL with a concentration of 200 μM and stored at −80 °C until further use.

### BAM complex reconstitution into nanodiscs

Protocol for nanodisc reconstitution was adapted from our former efforts^38^. The reconstitution of BAM into nanodiscs was done by 1,2-dimyristoyl-sn-glycero-3-phosphocholine (DMPC 14:0) lipids. For MSP1D1 reconstitution, a molar ratio of BAM: MSP: DMPC: sodium cholate (1: 6: 312: 624) was used resulting in the following molar concentrations: 20 μM: 120 μM: 6.3 mM: 12.6 mM, respectively. For BAM reconstitution into MSPdH5 nanodiscs, 50 µL BAM (83 µM), 210 µL MSPdH5 (200 µM), 11 µL DMPC (100 mM), 11 µL cholate (200 mM) were mixed resulting in molar ratios of approximately 1: 10: 265: 530, respectively. Samples were mixed in nanodisc buffer (25 mM Napi, 100 mM NaCl, pH 7.0) with Bio-beads™ SM-2 Adsorbent Media (Bio-RAD) for cholate and detergent (DDM) removal, mediating the nanodisc reconstitution process at room temperature on an orbital shaker overnight, forming an assembled nanodisc complex. Later on, the supernatant was collected and run on a Size Exclusion Chromatography column (Superdex 16600 s200pg 25mL).

### Electron microscopy sample preparation and data collection

Immediately before grid preparation, a gold quantifoil grid 2/1µm was glow-discharged using a Solarus Plasma Cleaner 950 (Gatan). A 3.5 μl aliquot of BAM complex in MSPdH5-Nanodiscs at a concentration of ∼0.8 mg/mL (2µM) was quickly mixed with a 100-fold molar excess of **2** and directly vitrified by plunging into liquid ethane using a Vitrobot (FEI, Vitrobot Mark IV). Data were collected at a Thermo Scientific Titan Krios electron microscope operated at 300 kV, equipped with a Falcon 4i Detector and an Selectris X imaging filter with a slit width of 10 eV. Images were acquired at -0.6 to -2.0 μm defocus and a nominal magnification of 165,000×, which corresponds to a pixel size of 0.73 Å. 29,988 movies were collected with a total accumulated dose of approximately 40 e−/Å^2^, fractionated over 40 frames, using beam-image shift.

### EM data processing and analysis

Micrographs were corrected for beam-induced drift using patch motion, and the contrast transfer function (CTF) parameters for each micrograph were determined using Patch CTF in cryoSPARC^39^. Particles were picked with the blob picker function in cryoSPARC and subjected to reference-free 2D classification. After an ab-initio reconstruction, heterogeneous classification identified two main populations, the BAM-compound **2** complex in the outward-closed conformation and the apo-state of BAM in the outward-open conformation. A simplified and summed representation of data processing is provided in Extended data Fig. 6. Both states were further refined separately, using non-uniform refinement, followed by local refinement using an automatically determined global mask. Using 3D-classification, we detected slight variability within the compound **2** binding region. The binding of **2** resulted in the opening of the BAM-barrel increasing the flexibility of the region comprising β-strands 1 and 16. We used the class with the best density for model building. Final maps at a global resolution of 3.03 Å (BAM-compound **2**) and 3.06 Å (BAM-Apo) were obtained, respectively. Modelling started by manual fitting of subunits or domains of the BamABCDE crystal structure (7NRI.pdb)^9^ into the EM density map in UCSF ChimeraX^40^. Model building was performed in Coot^41^ and real-space refinement was carried out in PHENIX.^42,43^ Validation was done using the cryo-EM validation MolProbity tools in PHENIX^44^. All structural models were visualized by ChimeraX^40^.

### BamA-β Expression and Purification

BamA-β (BamA residues 421–810 with mutations C690S, C700S) with N-terminal His6-tag were expressed from pET15b-based plasmids.21 E. coli Lemo21(DE3) cells were transformed with the plasmid, grown in LB or isotope-labeled medium to an OD600 of 0.6 and then induced with 1 mM IPTG. For [U-2H, 15N]-labelling, M9 minimal medium containing15NH4Cl in D2O was used. After overnight expression at 37 °C, cells were harvested by centrifugation (5000 g, 20 minutes). Cell pellets were resuspended and incubated in resuspension buffer (50 mM Tris-HCl, 300 mM NaCl, pH 8.0) supplemented with DNase 1 (Applichem) and lysozyme for 30 minutes at 4 °C. Cells were lysed by sonication and centrifuged (26’000 g, 60 minutes). The pellet was dissolved in denaturation buffer (50 mM Tris-HCl, 300 mM NaCl, 6 M GuHCl, pH 8.0) and centrifuged at 26’000 g for 30 minutes. Denatured cell extracts were subjected to Immobilized Metal Affinity Chromatography (IMAC) with a gravity column containing Ni-NTA charged beads being washed with running buffer (50 mM Tris-HCl, 300 mM NaCl, 6 M GuHCl, 20 mM imidazole, pH 8.0) and eluted with elution buffer (50 mM Tris-HCl, 300 mM NaCl, 6 M GuHCl, 200 mM imidazole, pH 8.0). Dissolved and unfolded BamA-β in the supernatant was refolded by dropwise addition into refolding buffer (50 mM Tris-HCl, 300 mM NaCl, 0.5% LDAO, 500 mM L-arginine, pH 8.0). The mixture was then dialyzed overnight against 5 liters of dialysis buffer (20 mM Tris, pH 8.0) followed by another dialysis round in fresh dialysis buffer, next day for 3 hours. Correctly folded protein fraction was purified by ion exchange chromatography on Hi-Trap Q HP (2 x 5 mL) columns, using buffer A (20 mM Tris-HCl, 0.1% LDAO, pH 8.0) and buffer B (20 mM Tris-HCl, 0.1% LDAO, 500 mM NaCl, pH 8.0). The protein was eluted with a gradient flow of buffer B. The pooled elution fractions were concentrated and applied to a HiLoad 16/600 Superdex 200pg (125 mL) size exclusion column in SEC buffer (HEPES-NaOH 20 mM, NaCl 150 mM, 0.1% LDAO, pH 7.5) for final purification. Purified protein was aliquoted, flash frozen in liquid nitrogen and stored at –80 °C to be used for SPR and NMR experiments.

### Surface plasmon resonance

SPR measurements were conducted on a Biacore T100 (T200 Sensitivity Enhanced, GE Healthcare Life Sciences). BamA-β with an N-terminal Avi-tag (GLNDIFEAQKIEWHE) was biotinylated via BirA and was captured via biotin on a sensor chip CAP (Cytiva) resulting in capture levels of 1600 RU after 7 blank injections. All runs were conducted with PBS buffer (10 mM phosphate, 150 mM NaCl, 0.1% (m/v) LDAO, pH 7.4). **2** was injected in increasing concentrations in triplicate for a multi cycle experiment with an increasing concentration profile equal to 0.0195, 0.039, 0.078, 0.156, 0.313, 0.625, 1.25, 2.5, 5, 10, 20 µM and a contact time of 120 sec and dissociation time of 500 sec at a flowrate of 30 µL/min. Reference and blank subtracted data were analyzed with the BiacoreT200 Evaluation software 3.0. The dissociation constant was fitted using the Steady State Affinity model and the kinetic parameters were fitted using the 1:1 binding model with RI jump deactivated.

### Thermostability analysis of protein samples

Thermal denaturation experiments were performed using a Nanotemper Prometheus NT.48 (Nano DSF) instrument. Samples were held in high sensitivity capillaries and subjected to a continuous thermal ramp from 20 – 95°C at a rate of 1°C/min. 10 µM of BAM complex reconstituted in nanodiscs in 20 mM Tris-HCl, pH 7.0 and 100 mM NaCl as buffer was mixed and briefly incubated with 20 µM of **2**. To account for dilution effect, volumetrically equal value of buffer was introduced to the control sample. Denaturation was monitored by intrinsic protein fluorescence using excitation at 280 nm and measuring emission at 330 and 350 nm. Denaturation curves were obtained by plotting the ratio of emission intensity at 350 nm to emission intensity at 330 nm as a function of temperature. The midpoint of thermal denaturation was obtained from the peak in the first-derivative of the curve using the automated algorithm in the instrument software.

### Solution NMR Spectroscopy

Protein samples for NMR spectroscopy were expressed in M9 medium, supplemented with perdeuterated water and 1.0 g/L 15N-ammonium chloride. BamA-β sample purified in 20 mM Hepes, pH 7.5, NaCl 150 mM, LDAO 0.1% w/v were concentrated to a final concentration of 300 µM. 2.0 equivalents of either dynobactin A or **2** were added to the sample and two-dimensional [^15^N, ^1^H]-TROSY-HSQC experiments were measured on a 700 MHz Bruker spectrometer equipped with a cryogenic probe. 128 transients with 1024 and 256 complex points in the ^1^H and ^15^N dimensions, respectively, were accumulated at a sample temperature of 37 °C for the initial apo sample, where the number of scans were increased accordingly upon each titration step to re-cover the protein dilution effect. The data were processed using Topspin 3.6.2 and analysed using CcpNmr version 3.0. Chemical shift perturbation of the amide backbone was calculated as:

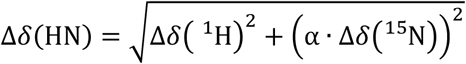

Titration of the drug candidate with the protein samples was performed in three steps accounting for 0.5, 1.0 and 2.0 equivalent of the drug candidate in molar equivalent with respect to the apo sample concentration. CSPs of each candidate were calculated and plotted using Microsoft Excel and matplotlib library of python 3.

### PISA assay sample preparation

PISA assay was performed in 4 replicates for both cell and lysate experiments. *E. coli* K12-MG1655 cells carrying an ampicillin resistance plasmid were plated on ampicillin-containing plates. A single colony was picked and cultured overnight in LB media. The following day, a fresh LB culture supplemented with ampicillin was prepared and cells were inoculated on a volumetric ratio of 1:1000 and grown until the optical density at 600 nm (OD_600_) reached 0.5. Cells were then pelleted by centrifugation at 3000 g for 20 minutes and divided into two groups for lysate and cell experiments. For lysate experiment, lysis of the cells was performed through five freeze-thaw cycles in liquid nitrogen using a lysis buffer composed of 20 mL PBS at pH 6.8 containing 1x HALT protease inhibitor (Thermo Scientific), 40 µL lysozyme and 4 µL benzonase (E1014-5KU, Sigma-Aldrich). The lysates were centrifuged at 7000x g for 5 minutes to remove cell debris. Both lysate and cell samples were then treated with ligands: Dynobactin A at 10 µM was added to the lysates and incubated for 15 minutes, and **2** at 10 µM was added to both lysate and cell conditions also for 15 minutes. An equal volume of only PBS at pH 6.8 was also prepared for the control set. The samples were then aliquoted into 96-well PCR plates at 40 µL per well and were individually exposed to a temperature gradient of 44.0, 45.5, 47.3, 50.0, 53.0, 56.0, 59.0, 62.0, 65.0, 67.7, 69.5, and 70.1 °C using a qPCR machine (Analytik Jena). After 3 min at room temperature, cells were snap-frozen in liquid nitrogen. Upon thawing, the aliquots heated to the different temperature points within each replicate were combined.

For the cell experiments, cells were first treated with compounds, aliquoted and then individually heated to the above temperatures, kept 3 min at room temperature and then snap-frozen in liquid nitrogen. Upon thawing, the aliquots heated to the different temperature points within each replicate were combined, and lysis buffer containing 80 µL NP-40 was added to each sample, followed by 4 more rounds of free-thawing. The lysate and cell samples were then centrifuged at 100,000g for 20 min at 4 °C, and supernatants were taken into new tubes. The protein concentration in each sample was measured and 20 µg of the protein mixture was subjected to sample preparation for LC-MS/MS analysis S-Trap™ micro spin columns (Protifi) as described below.

### Sample digest and proteomics analysis

20 µg of protein extract from the PISA assay were collected and resuspended in lysis buffer (5% SDS, 10mM TCEP, 0.1 M TEAB) and lysed by sonication using a PIXUL Multi-Sample Sonicator (Active Motif) with Pulse set to 50, PRF to 1, Process Time to 10 min and Burst Rate to 20 Hz. Lysates were incubated for 10 min at 95°C, cleared by centrifugation, alkylated in 20 mM iodoacetamide for 30 min at 25°C and proteins digested using S-Trap™ micro spin columns (Protifi) according to the manufacturer’s instructions. Shortly, 12 % phosphoric acid was added to each sample (final concentration of phosphoric acid 1.2%) followed by the addition of S-trap buffer (90% methanol, 100 mM TEAB pH 7.1) at a ratio of 6:1. Samples were mixed by vortexing and loaded onto S-trap columns by centrifugation at 4000 g for 1 min followed by three washes with S-trap buffer. Digestion buffer (50 mM TEAB pH 8.0) containing sequencing-grade modified trypsin (1/25, w/w; Promega, Madison, Wisconsin) was added to the S-trap column and incubate for 2h at 47 °C. Peptides were eluted by the consecutive addition and collection by centrifugation at 4000 g for 1 min of 40 ul digestion buffer, 40 μL of 0.2% formic acid and finally 35 μL 50% acetonitrile, 0.2% formic acid. Samples were dried under vacuum and stored at -20 °C until further use.

Dried peptides were resuspended in 0.1% aqueous formic acid and subjected to LC–MS/MS analysis using a Orbitrap Fusion Lumos Mass Spectrometer fitted with an EASY-nLC 1200 (both Thermo Fisher Scientific) and a custom-made column heater set to 60°C. Peptides were resolved using a RP-HPLC column (75μm × 36cm) packed in-house with C18 resin (ReproSil-Pur C18– AQ, 1.9 μm resin; Dr. Maisch GmbH) at a flow rate of 0.2 μL/min. The following gradient was used for peptide separation: from 5% B to 12% B over 5 min to 35% B over 40 min to 50% B over 15 min to 95% B over 2 min followed by 18 min at 95% B. Buffer A was 0.1% formic acid in water and buffer B was 80% acetonitrile, 0.1% formic acid in water.

The mass spectrometer was operated in DIA mode. MS1 scans were acquired in the Orbitrap in centroid mode at a resolution of 120,000 FWHM (at 200 m/z), a scan range from 390 to 910 m/z, AGC target set to 250 % and a maximum ion injection time of 50 ms. MS2 scans were acquired in the Orbitrap in centroid mode at a resolution of 15,000 FWHM (at 200 m/z), precursor mass range of 400 to 900, quadrupole isolation window of 16 m/z with 1 m/z window overlap, a scan range from 145 to 1450 m/z, normalized AGC target set to 2000 % and a maximum ion injection time of 22 ms. Peptides were fragmented by HCD (Higher-energy collisional dissociation) with collision energy set to 33 % and one microscan was acquired for each spectrum.

The acquired raw-files were searched using the Spectronaut (Biognosys v18.5.231110.55695) directDIA workflow against an E coli database (consisting of 4390 protein sequences downloaded from Uniprot on 20220222) using default factory settings. Quantitative data was exported from Spectronaut and analyzed using the MSstats R package v.4.8.6. (https://doi.org/10.1093/bioinformatics/btu305).

### PISA assay data analysis

The raw abundances were normalized to the total sample intensity. Fold changes were then calculated by dividing the normalized intensities in the treated samples vs. control. Two-sided unpaired t-test was applied to calculate p-values between the normalized intensities of treated and control samples.

### Growth-inhibition assay

Overnight cultures of *E. coli* K12 MG1655 wild-type and *ΔbamB* were used to inoculate 3 mL fresh LB supplemented with 1 μg/mL **2** or 4 μg/mL dynobactin A or no antibiotic to a starting optical density at 600 nm (OD_600_) of 0.1. Cells were incubated in these different conditions at 37°C. After 0, 2, 5 and 7 hours, 40 μL of each condition were used for a serial dilution (10^0^ to 10^-7^) with LB and 5 μL of each dilution were spotted on a LB plate. CFU were counted after incubating the plates 16 hours at 37°C. The experiment was performed in triplicates.

### AlphaFold predictions

Protein structure predictions were conducted using AlphaFold version 3.0, implemented on the AlphaFold webserver. The FASTA file for bamA of *Escherichia coli* K12 (P0A940) was sourced from UniProt and uploaded to the sciCORE environment. For the resistant strains, the *bamA* gene was first annotated using Geneious Prime 2025.0.2 (https://www.geneious.com) from Whole Genome Sequencing data of *Acinetobacter baumannii* (ATCC 19606) and *Pseudomonas aeruginosa* (ATCC 9027). The annotated *bamA* sequences were then utilized for AlphaFold predictions.

### AlphaFold data analysis

To calculate and visualize pLDDT scores, the output JSON files, containing keys such as atom_plddts (pLDDT scores) and token_res_ids (residue IDs), were loaded into Python using the json library. The atom_plddts key provided a list of pLDDT scores, while the token_res_ids key provided a list of corresponding residue IDs. A dictionary was created to map each residue ID to its corresponding pLDDT scores. By iterating through the token_res_ids and atom_plddts lists simultaneously, each pLDDT score was appended to the list of scores for the corresponding residue ID in the dictionary. For each residue ID in the dictionary, the average pLDDT score was computed by calculating the mean of the list of pLDDT scores for that residue. Two lists were created: one for residue IDs and one for their corresponding average pLDDT scores, which were used for plotting and exporting data. A line plot was created with residue IDs on the y-axis and average pLDDT scores on the x-axis using the matplotlib library.

### Data availability

Proteomics data have been deposited to the ProteomeXchange Consortium (https://www.proteomexchange.org/) via the MassIVE partner repository with MassIVE data set identifier MSV000095669 and ProteomeXchange identifier PXD055028. Python codes for data analysis are available upon request. PDB deposition, PDB ID: 9HE1. EMDB deposition, EMD-52074 (for Electron Microscopy Database)

### Killing kinetics determination

Peptides stock solutions were diluted into MHB to desired concentrations. *E. coli* M6 culture in mid-log phase was diluted into MHB to yield 10^6^ CFU/mL. The mixture was incubated at 37 °C with shaking. At each time point, 10 µL of the sample was drawn out and subjected to ten-fold serial dilution. 20 µL of relevant dilutions was dropped onto MHA plate using the drop plate method. The plate was incubated for 18–20 h at 37 °C. Colony number was counted, and used for calculating the CFU/mL according to the equation: CFU/mL = Colony count × 50 × dilution factor

### Field-emission scanning electron microscopy (FE-SEM) microscopy

*E. coli* M6 culture at mid-log phase was diluted to an OD_600_ of 0.1. After incubating the bacteria with the peptide at 8×MIC for 1 h, 2 h, or 4 h at 37 °C with shaking, the samples were washed thrice in PBS. After overnight fixation with 2.5% glutaraldehyde (in PBS) at 4 °C, the samples were washed twice in PBS, and then re-suspended in 500 µL of PBS. Sample was dropped onto cover slips pre-treated with poly-l-lysine. After 30 min, unbound cells were washed away with PBS. Following post-fixation with 1% OsO_4_ for 30 min, OsO_4_ was removed, and the cover slips were washed twice with distilled water. Samples were dehydrated using a series of ethanol solutions (50%, 75%, 95%, 3 × 100%). They were then subjected to critical point drying using Leica EM CPD300 (Wetzlar, Germany), followed by sputter gold coating using Leica EM ACE200 (Wetzlar, Germany). Viewing of the samples was performed using JEOL JSM-6701F (Tokyo, Japan). Images were processed using ImageJ (National Institutes of Health, Bethesda, MD).

### Serial passage

Resistance development of *E. coli* M6 against **2** was assessed by serial passaging of the bacteria in MHB containing subinhibitory concentrations of the peptide. In brief, bacteria culture at mid-log phase was diluted to 10^5^-10^6^ CFU/mL in MHB containing 0.25×, 0.5×, 1×, 2×, and 4× MIC of the peptide. After 24h of incubation (37 °C, 120 rpm shaking), the new visually determined MIC value was recorded, and the culture at highest peptide concentration showing visible growth was diluted to 10^5^-10^6^ CFU/mL in MHB. A new set of peptide concentration range was added to the cultures based on the latest MIC. This process was repeated over 14 days for three independent starting cultures.

### *In vivo* efficacy in peritonitis model

All animal procedures were performed in accordance with protocols approved by the Institutional Animal Care and Use Committee (IACUC) at National University of Singapore (Singapore). Female C57BL/6NTac mice aged 6-8 weeks were acquired from InVivos Pte Ltd (Singapore, Singapore). Solutions for injections were prepared fresh in pharmaceutical grade saline and filter-sterilized. Murine peritonitis model was established according to the literature^45^. Briefly, healthy mice were rendered neutropenic by administering i.p. injection (0.5 mL) of cyclophosphamide on day -4 (150 mg/kg) and day -1 (100 mg/kg). On day 0, mice were infected with *E. coli* M6 (10^9^ CFU/mL) through i.p. injection (0.1 mL). At 30 min post-inoculation, mice were given i.p. injection (0.5 mL) of a single dose of Smc (5 or 50 mg/kg), colistin (5 mg/kg), or saline control (n = 5 mice per treatment group). At 2 h post-treatment, mice were humanely euthanized by carbon dioxide asphyxiation and cervical dislocation. Sterile PBS (3 mL) was injected into the peritoneal cavity, followed by abdominal massage and collection of peritoneal fluid (1-2 mL). Blood (0.3-0.5 mL) was collected through cardiac puncture. Liver, spleen, and kidney were surgically removed and stored in 0.1% Triton X-100 (in PBS). Tissue homogenization was performed using gentleMACS dissociator (Miltenyi Biotec, Germany) by following a published protocol^46^. Cell aggregates were removed using a 30 µm mesh MACS SmartStrainer (Miltenyi Biotec). Blood, peritoneal fluid, and tissue homogenates were plated on LB agar and incubated overnight for colony counting.

## Author Contributions

SMM, RS, SH, and BIM designed the study; RS, YM, J L., C-S P, PSYL, ZHL, YH, JYL, and BIM designed and carried out experiments for production and characterization of natural products; RS, NDTT, XJ, SYHL, PWLC, AYJY, KTSK, WYG, MY, PLRE, BIM designed and carried out experiments for biological evaluation and resistance of natural products.; SMM and SH designed and carried out experiments on the mode of action of natural products. SMM, RS, YM, NDTT, PLRE, SH and BIM wrote the paper.

## Declaration of Interests

RS, YM, NDTT, JL, PLRE, and BIM are inventors on SG Patent Application No. 10202250612J

## Acknowledgments

The authors acknowledge financial support from the Singapore Ministry of Education (A-0004623- 00-00, A-0004318-00-00, A-0004629-00-00, A-0008472-00-00, and A-8001694-00-00 to B. I. M., and A-0004347-00-00 to P.L.R.E.), Singapore National Research Foundation SINERGY Grant (A-0008472-00-00 to B. I. M.), WuXi AppTec - NUS Grant to B. I. M., the Human Frontier Science Program to R. S., the Naito Foundation to Y. M. and the Japan Society for Promotion of Science (2019-60269 to R. S., 2021-60291 to Y. M., and 22H05124 to M. Y.). We thank Dr. Jingsong Fan (NUS Department of Biological Sciences) for assistance with NMR measurements, and Dr Lakshminarayanan Rajamani (Singapore Eye Research Institute) and Dr Jeanette Teo (National University Hospital of Singapore) for providing the clinical bacterial strains. We thank BioEM lab of the University of Basel for support with cryo-EM data acquisition as well as Proteomics Core Facility (PCF) of the Biozentrum, University of Basel. We thank Dr. Alexander Harms for *bamB* knock out cells. Calculations were performed at sciCORE (http://scicore.unibas.ch/) scientific computing core facility at the University of Basel. This project was supported by Swiss National Science Foundation via the National Center of Competence in Research AntiResist (180541). A.A.S. acknowledges the Swiss National Science Foundation (Ambizione Fellowship, SNSF grant number: PZ00P3_216203) for their support. We thank Prof. Kim Lewis (Northeastern University) for a sample of dynobactin and darobactin- and dynobactin-resistant strains of *E. coli*.

## EXTENDED DATA FIGURES AND TABLES

**Extended Data Figure 1.**
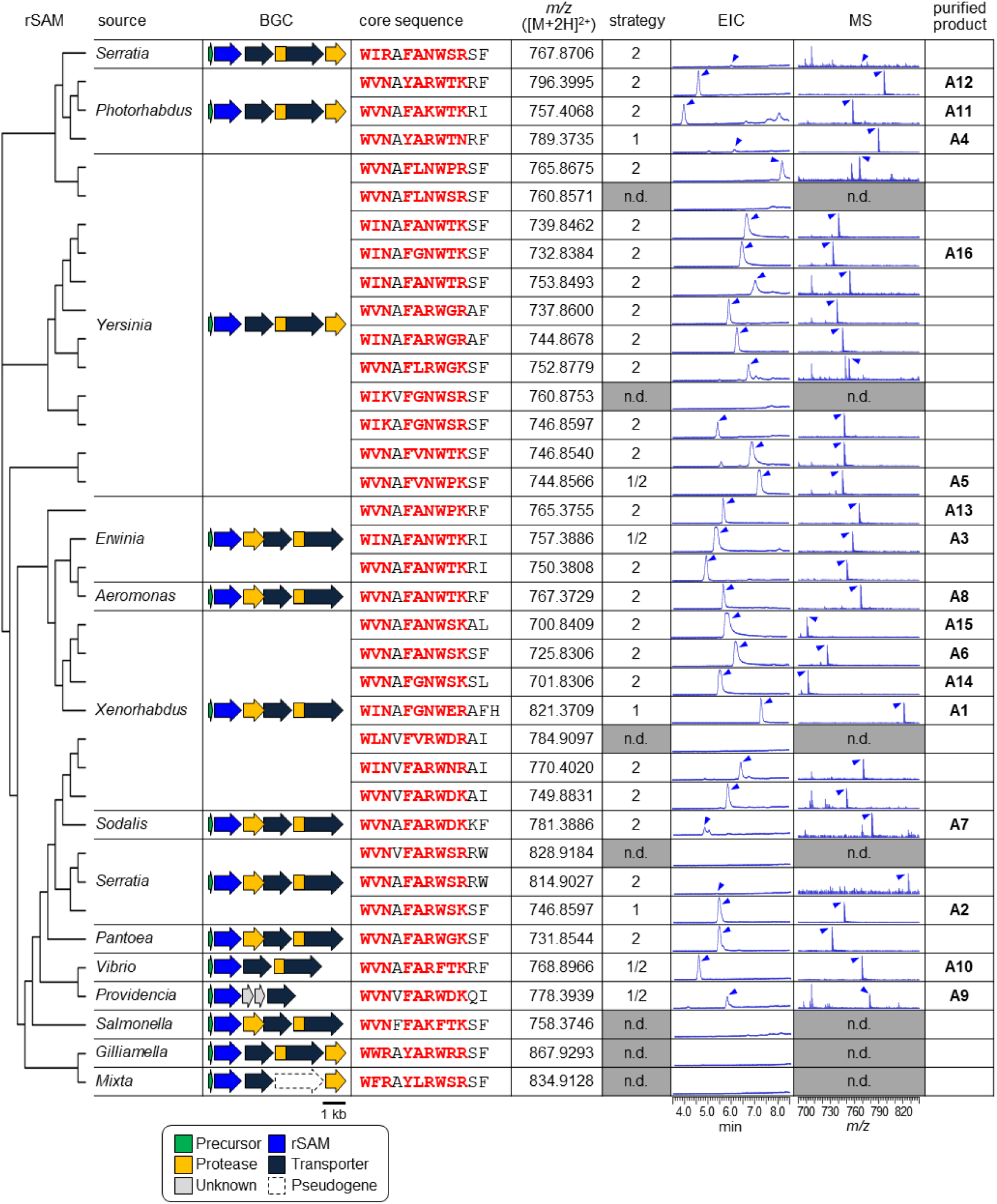
Comprehensive production of Type A xenorceptides. Production of Type A xenorceptides with 37 unique core sequences were produced in two strategies. The table summarizes the genera of source bacteria, representative BGC diagrams with core sequences, theoretical m/z values of natural products corresponding to -6 Da mass loss from those of unmodified core peptides, and their extracted ion chromatogram (EIC) and mass spectra (MS). The phylogenetic tree made by Clustal Omega summarizes gene sequences encoding rSAM/SPASM XyeB proteins associated with a XyeA precursor peptide with a unique core. Potential cyclized three-residue motifs in the core sequences are indicated in red. 16 products named xenorceptides A1- A16 were purified and subjected to subsequent experiments.

**Extended Data Figure 2.**
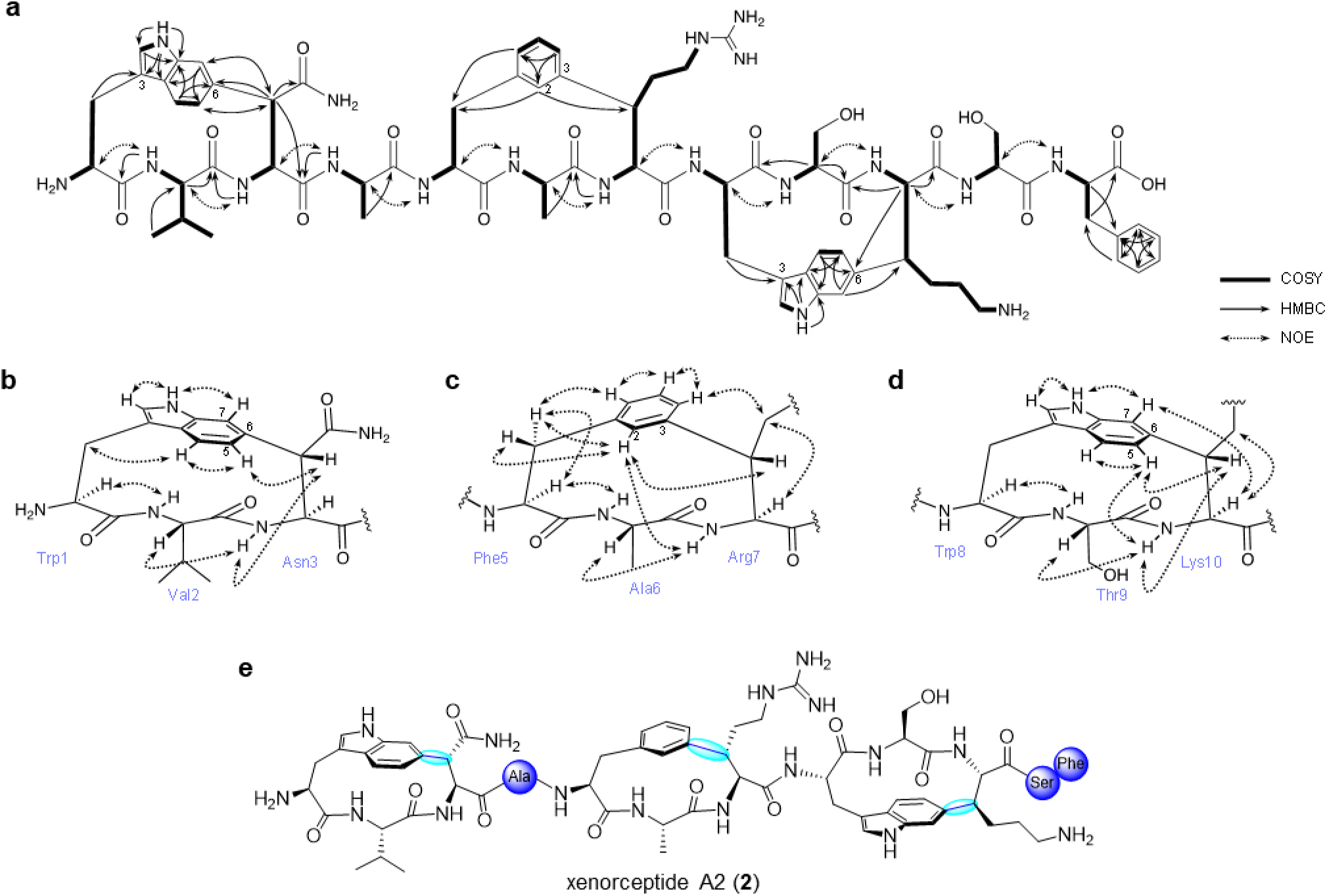
Structure elucidation of xenorceptide A2 (2). (a) Key 2D NMR correlations of **2**. (b-d) Conformational analysis with NOE correlations for WVN (b), FAR (c), and WTK (d) motifs. (e) The deduced chemical structure of **2**.

**Extended Data Figure 3.**
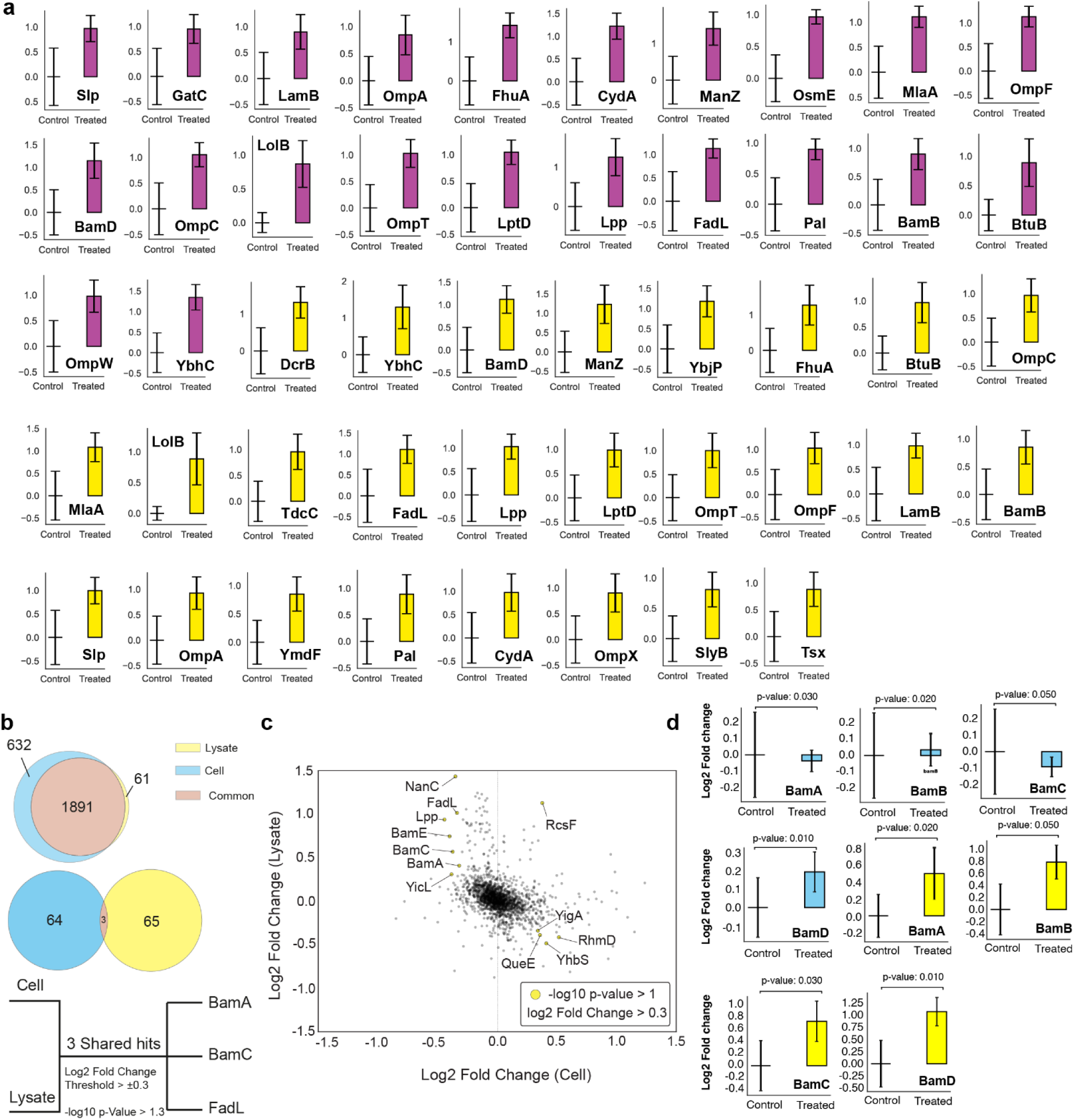
Protein Integral Solubility Alteration Assay of bamabactins performed on *E. coli* K12 MG1655 Cells and Lysates. (a) Bar plots showing log2 fold changes (y-axis) in measured intensity of detected peptides for dynobactin A (magenta) and xenorceptide A2 (yellow) treated samples compared to controls. (b) Pair-wise Venn diagram illustrating the total proteins detected in cell and lysate datasets, as well as shared hits with the most significant changes in solubility between the two datasets. (c) Scatter plot depicting observed log2 fold changes in cell and lysate PISA experiments. Hits passing defined filter (cutoff: -log 10 p-value > 1, arbitrary, log2FC > 0.3) are shown in yellow circles (d) Bar plots showing log2 fold changes in BAM complex components along with their corresponding p-values.

**Extended Data Figure 4.**
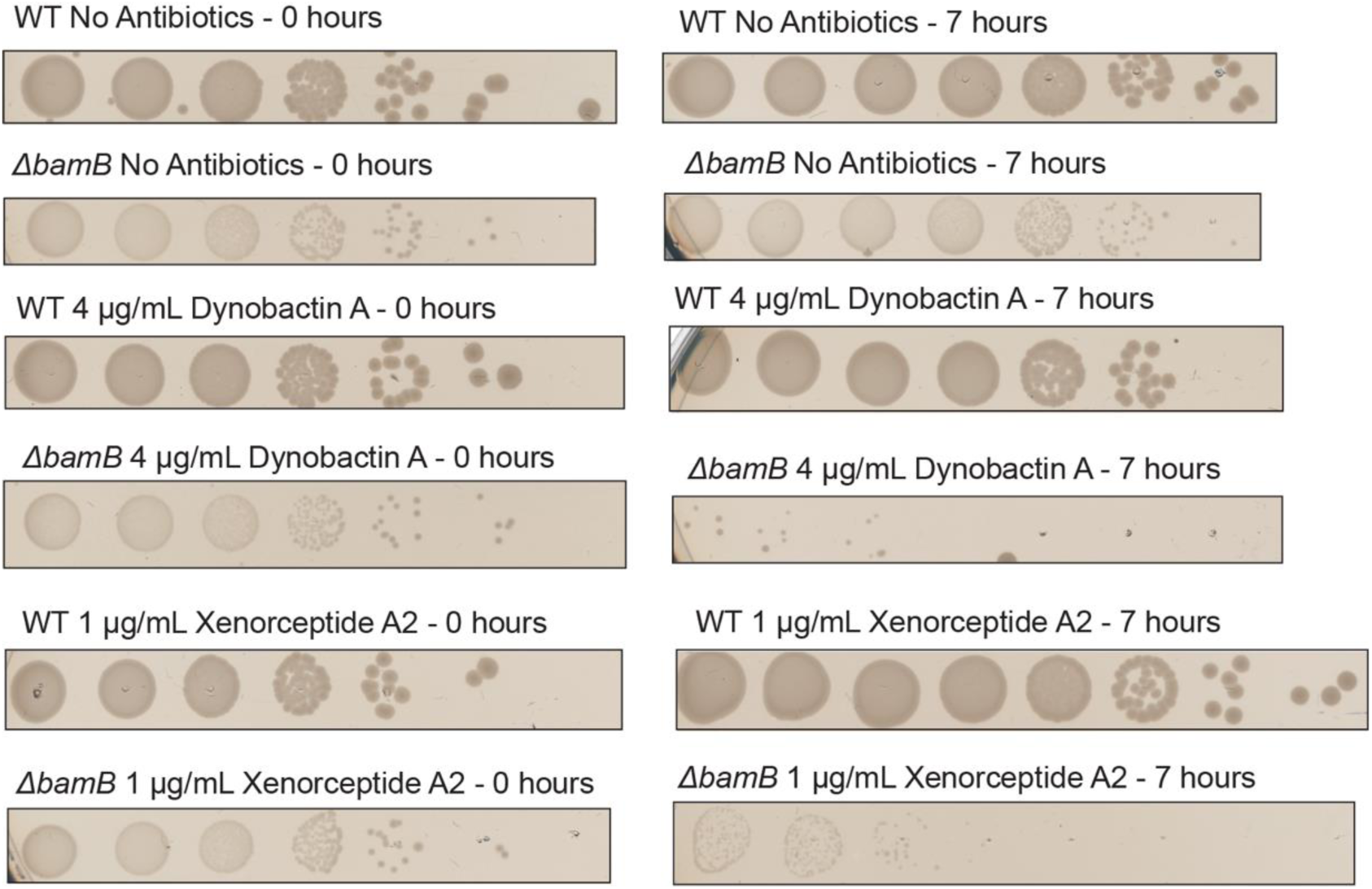
Growth Inhibition assay (Colony Forming Unit) Assay for *E. coli* K12 MG1655 and *ΔbamB* Mutant Strains. Colony formation of wild-type (WT) and *ΔbamB* mutant *E. coli* K12 MG1655 strains under different conditions at 0 and 7 hours as readout points. Two first row: WT and *ΔbamB* cells without antibiotics at 0 and 7 hours. Third and fourth row: WT and *ΔbamB* cells treated with 4 µg/mL dynobactin A at 0 and 7 hours. Fifth and sixth row: WT and *ΔbamB* cells treated with 1 µg/mL xenorceptide A2 at 0 and 7 hours. Each condition shows a series of dilution spots to indicate the number of viable colonies. Sub-MIC treatment of *ΔbamB* cells show higher susceptibility compared to WT cells to both dynobactin A and xenorceptide A2.

**Extended Data Figure 5.**
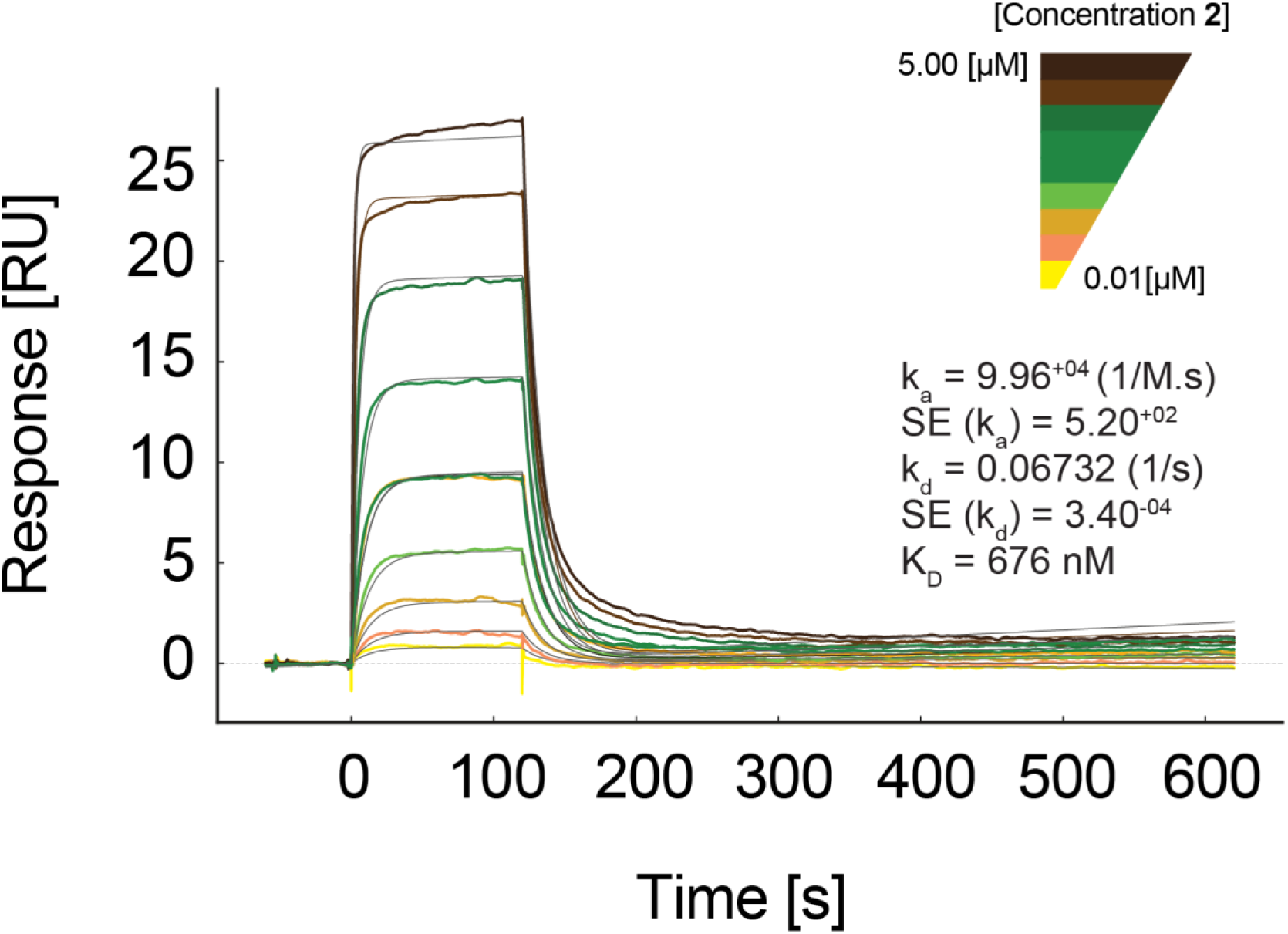
Surface Plasmon Resonance (SPR) sensorgrams (colored) and fitted curves from the 1:1 binding model (black). Kinetics parameters for the binding of **2** to BamA-β in LDAO micelles were determined using the 1:1: binding model. Injections were performed at concentrations of 0.01, 0.03, 0.07, 0.15, 0.31, 0.62, 1.25, 2.50, and 5.00 µM. Kinetics parameters of the binding events are plotted on the figure and the extracted dissociation constant (K_D_=676 nM) is consistent with the steady-state measurements (K_D_=800 nM). k_a_ and k_d_ stands for association and dissociation rates, respectively and SE stands for standard error.

**Extended Data Figure 6.**
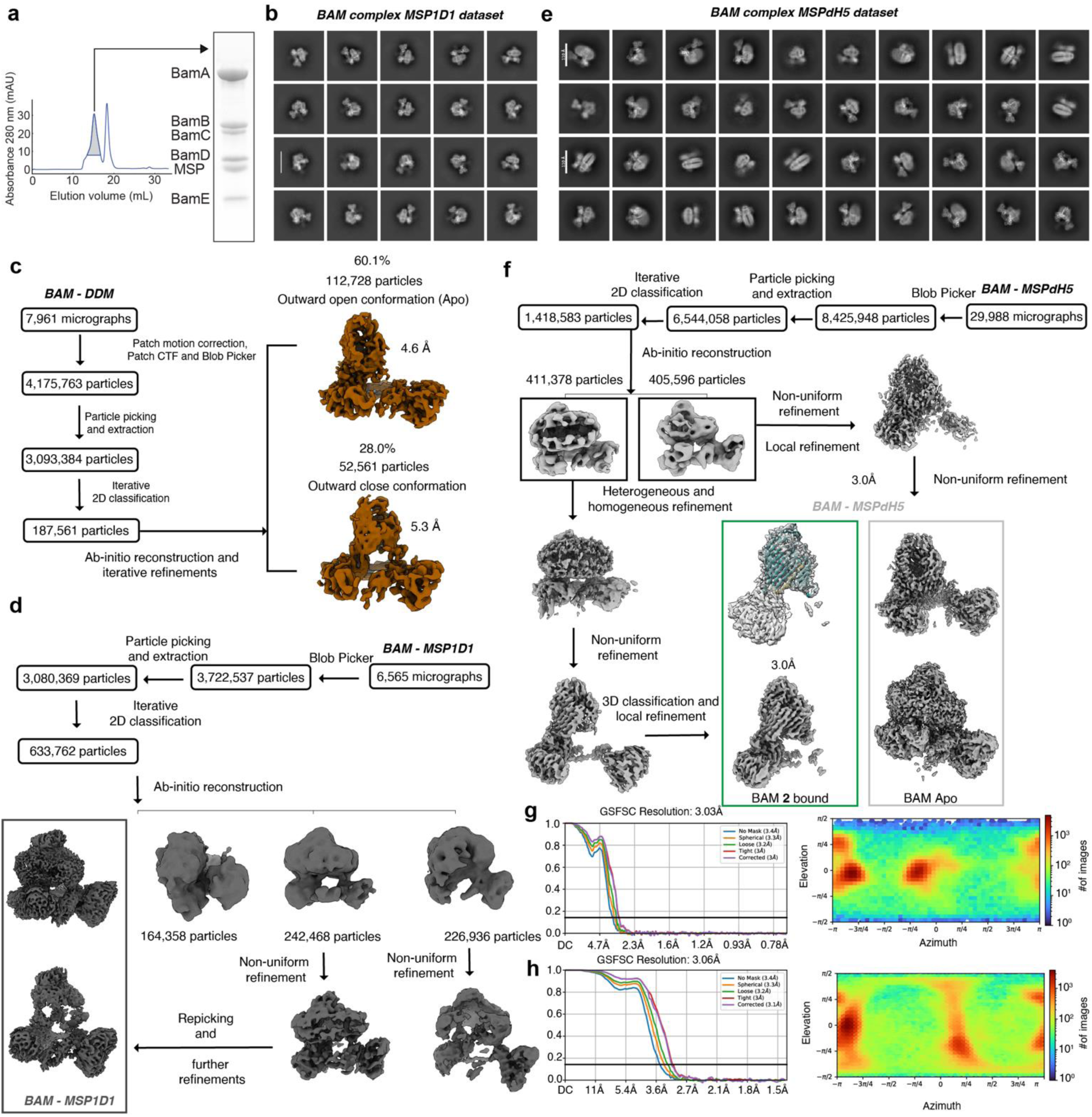
Cryo-EM workflow. (a) Size exclusion chromatogram of the BAM complex reconstituted in nanodiscs. The first peak corresponds to the assembled proteo-nanodisc complex, where its larger size leads to reduced dispersion and earlier elution from the column as well as SDS-PAGE (Sodium Dodecyl Sulfate–Polyacrylamide Gel Electrophoresis) of BAM complex reconstituted in nanodisc, 6 bands corresponding to penta-heteromeric BAM reconstituted in DMPC (14:0) lipid bilayers via MSP (Membrane Scaffold Protein). (b) Two-dimensional class averages of BAM in MSP1D1 (c) Flowchart for data processing to generate the cryo-EM map of BAM in DDM micelles, leading toward maps from 28.0 % of particles bound to **2** in 5.3 Å resolution and 60.1 % (majority) of the protein in the apo state and (d) Same for BAM complex reconstituted in MSP1D1 nanodiscs, where the gate region was not well-resolved. (e) Two-dimensional class averages of BAM in MSPdH5 nanodiscs – bar 110 Å. (f) Data processing flowchart of BAM in MSPdH5 nanodiscs giving cryo-EM map of BAM bound to **2** in 3.0 Å resolution. (g) Fourier Shell Correlation (FSC) plot versus spatial frequency and view direction distribution for conformation BAM bound to **2** and (h) same for apo BAM in MSPdH5 nanodiscs.

**Extended Data Figure 7.**
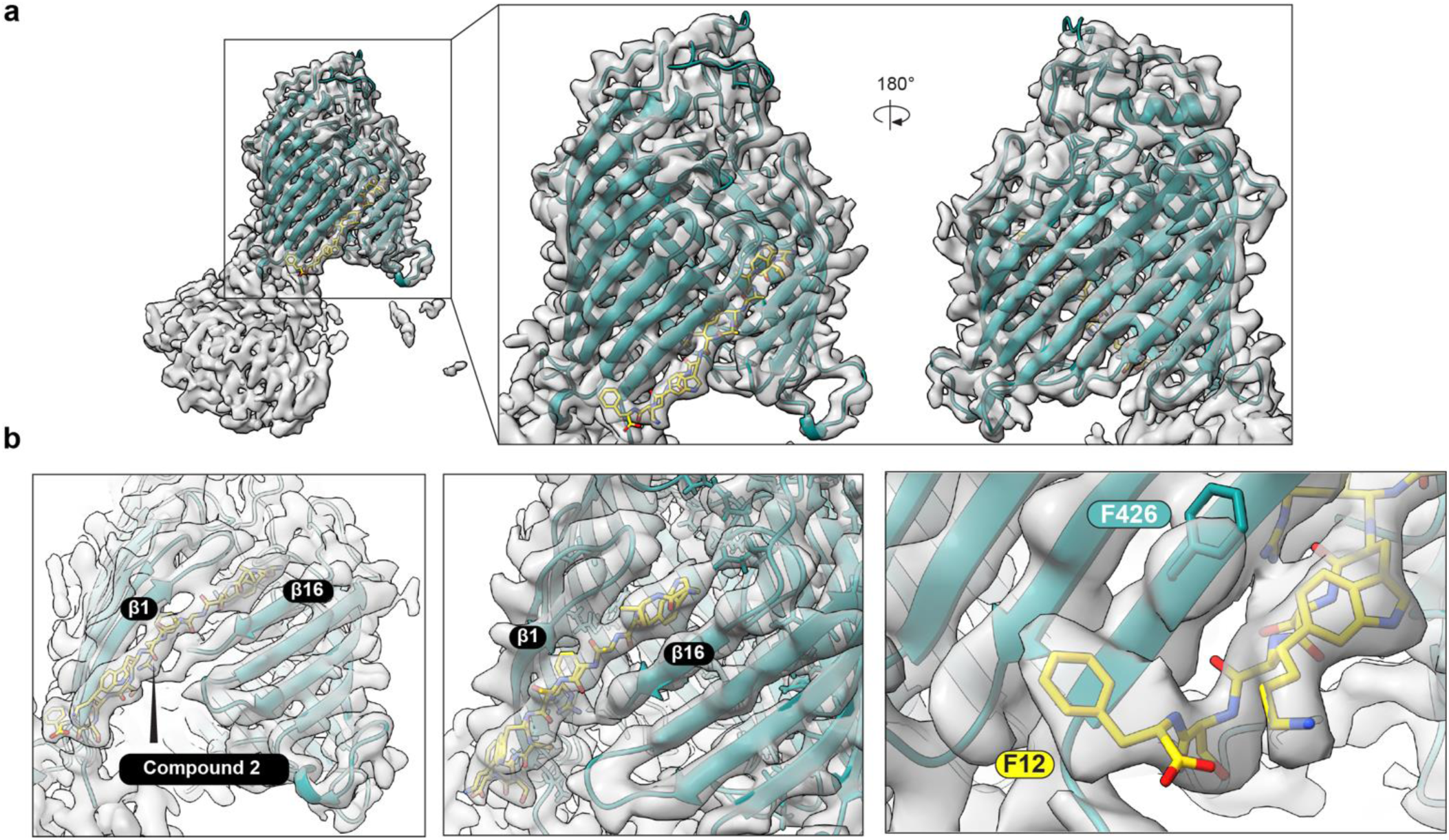
Structural details from the columb potential map of cryo-EM data. (a) Overview of the cryo-EM reconstruction of the BAM-**2** complex. Structural model of BAM is illustrated as ribbon and colored in teal, while the coulomb potential map is shown as gray transparent surfaces. (b) Side and lateral view of BamA bound to **2** as well as density and constructed structural model of phenylalanine 12 of **2** fitted in the recognition binding pocket located at the lateral gate of BamA.

**Extended Data Figure 8.**
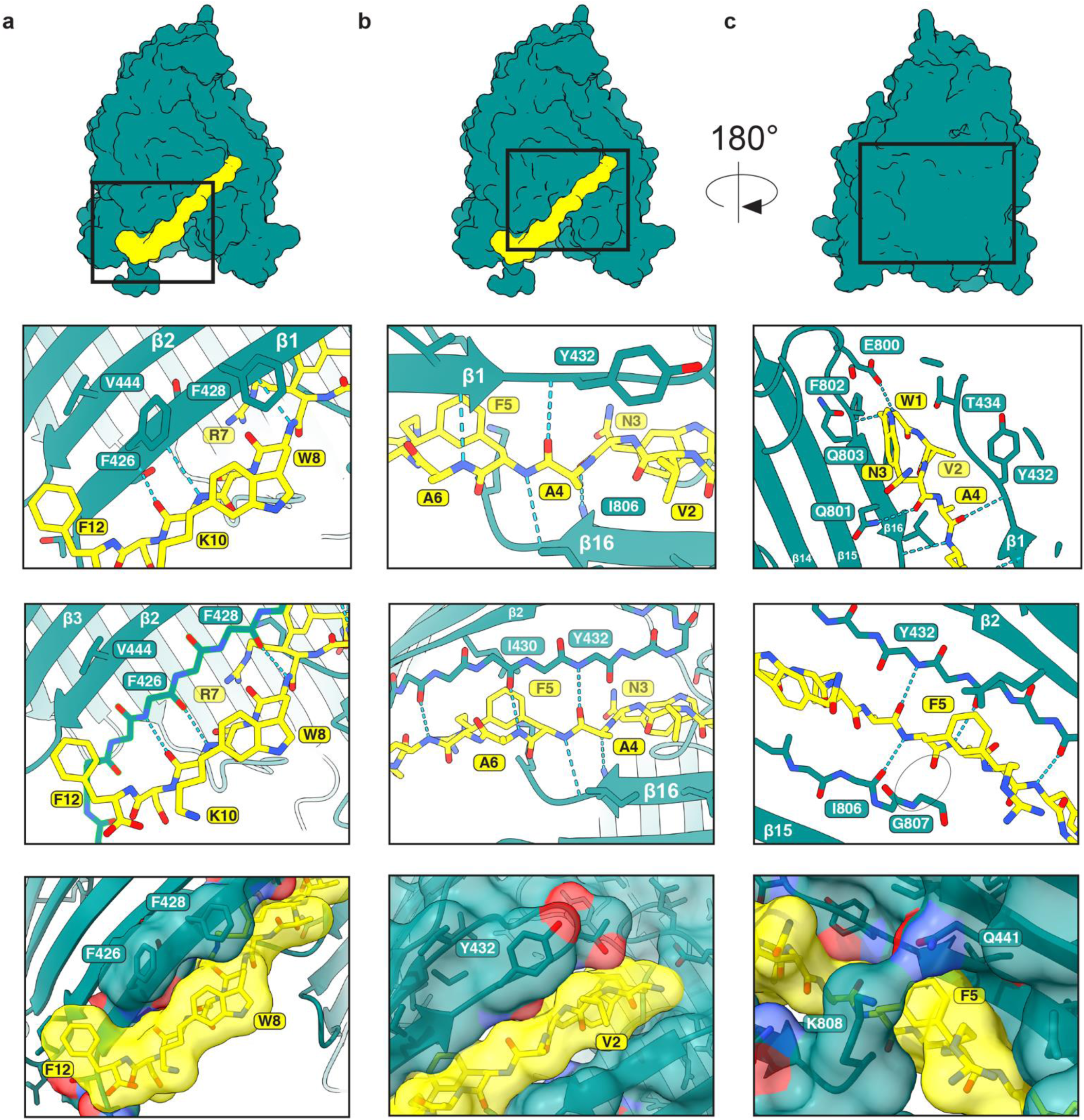
Binding interface details. Cryo-EM structure BAM complex bound to **2**. Panels represent details of BamA-β with bound **2** at 3.0 Å resolution. Zoomed-in panels highlight specific residues involved in the interaction

**Extended Data Figure 9.**
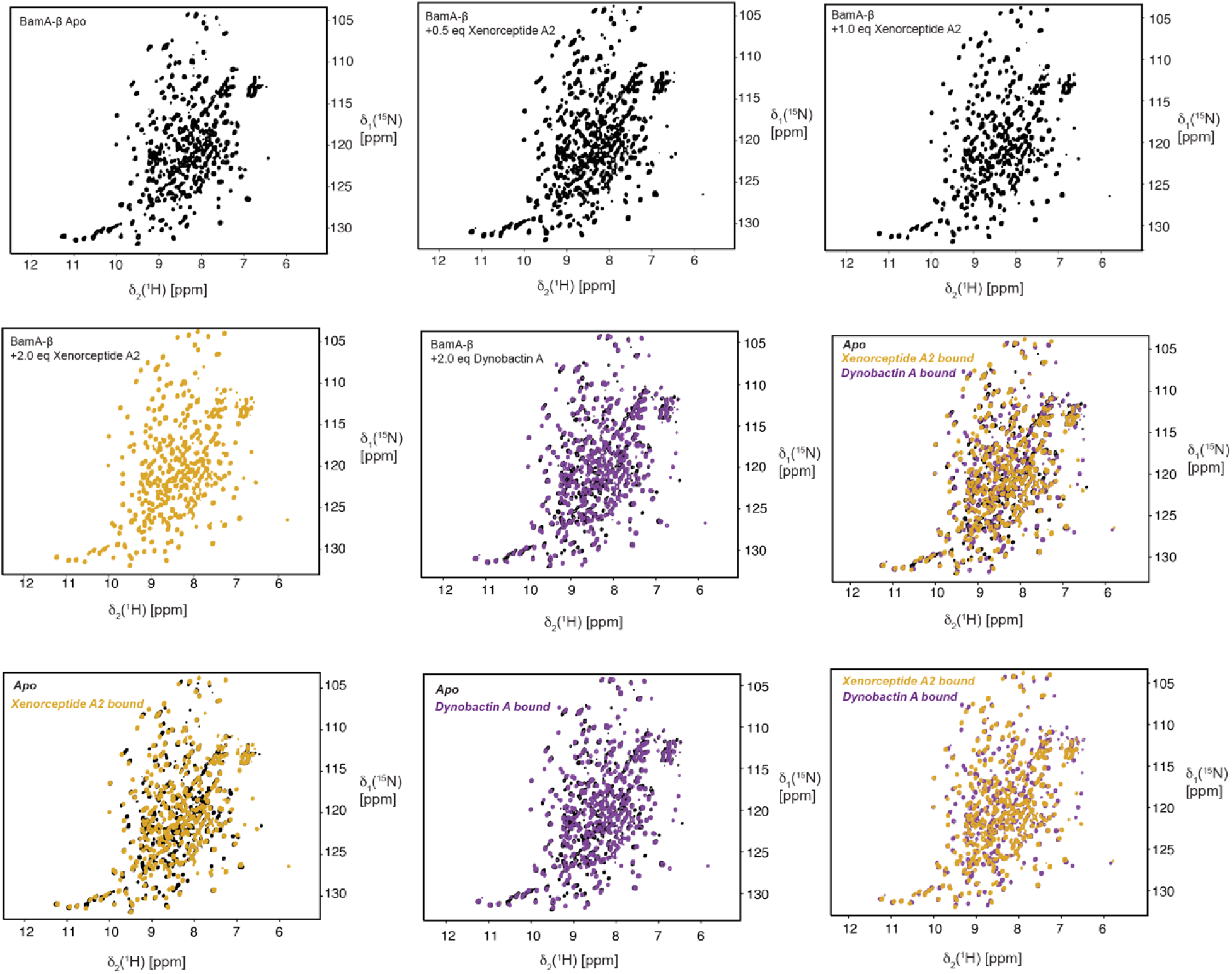
Bamabactin binding monitored by solution NMR spectroscopy. 2D [^15^N,^1^H]- TROSY spectra of apo BamA-β in LDAO micelles (black) overlaid with BamA-β with 2.0 eq of **2** (Xenorceptide A2) (yellow) and 2.0 eq of dynobactin A (magenta). A combination across various spectra is made visualizing the unique conformational changes in BamA-β upon interaction with each natural peptide.

**Extended Data Figure 10.**
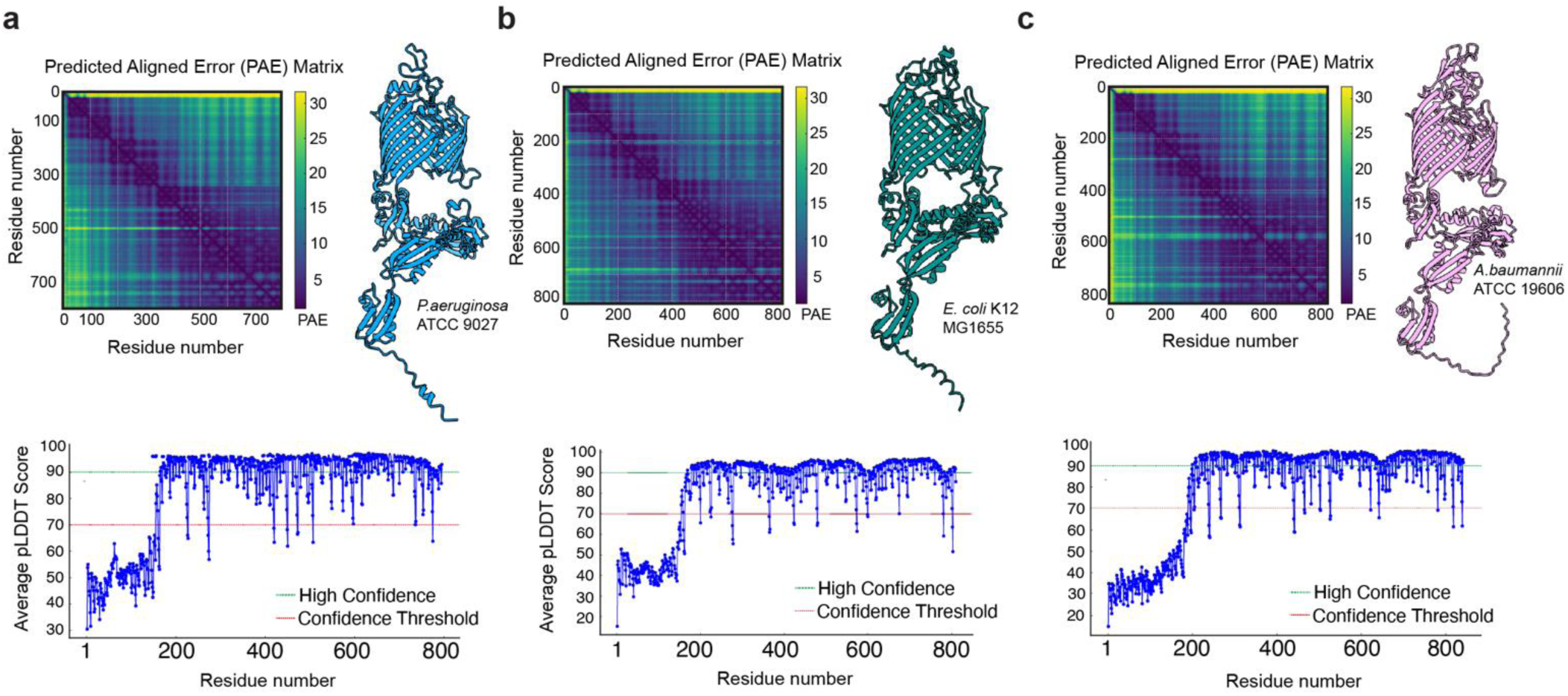
AlphaFold 3.0 predictions of 2-resistant BamA mutants. Predicted structures of full-length BamA from (a) *Pseudomonas aeruginosa* ATCC 9027, (b) *Escherichia coli* K12 MG1655, and (c) *Acinetobacter baumannii* ATCC 19606, along with their respective pLDDT scores plotted against the amino acid sequence of BamA. AlphaFold 3.0 provides highly accurate predictions for the BamA β-barrel, as indicated by elevated pLDDT values.

**Extended Data Figure 11.**
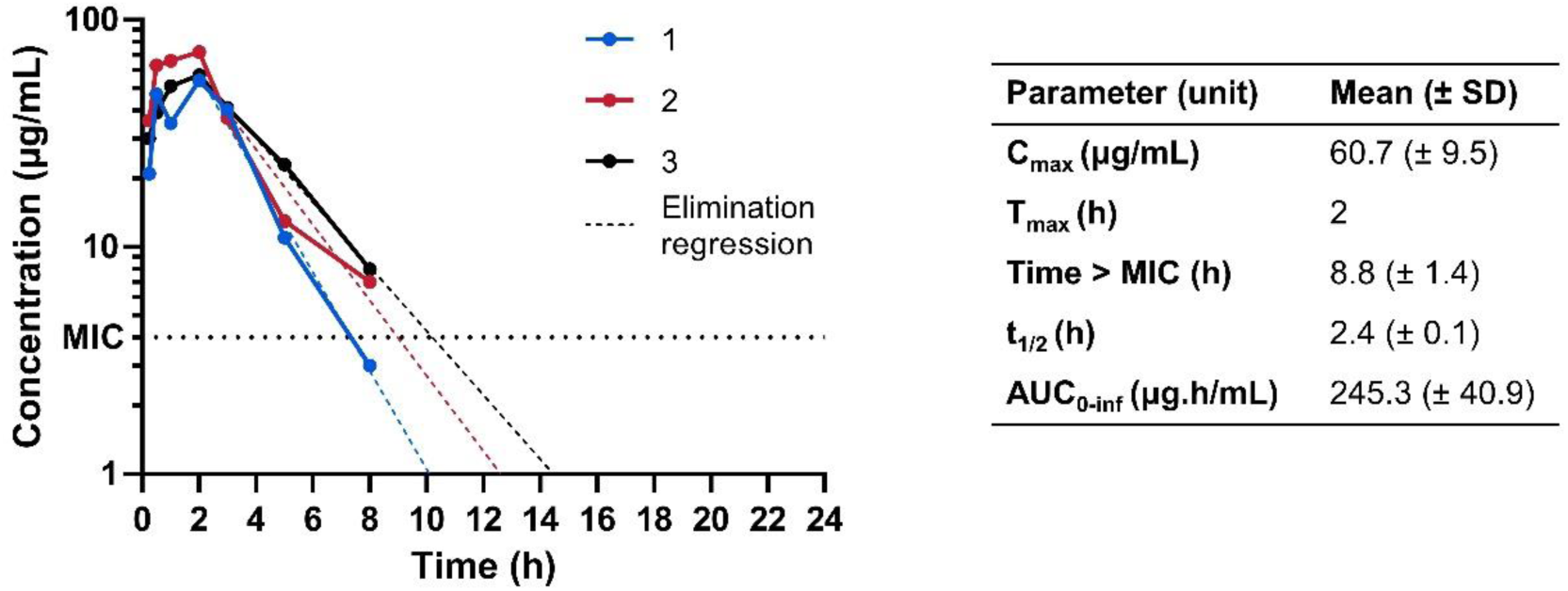
Three mice were intraperitoneally injected with 50 mg/kg **2**, and blood samples were collected by tail snip over 24 h. Samples (n = 1 per time point and mouse) were analysed for **2** content by LC-MS/MS, and concentrations were calculated using a standard curve created by linear regression on the log (area under the curve peak) to log(concentration) of standards. Pharmacokinetic values were calculated in Excel; t_1/2_ and time > MIC assuming first-order elimination and using linear regression on time points 0-8 h; AUC (0–16 h) using the trapezoid rule. The limit of detection (LOD) was 10 ng/ml.

**Extended Data Table 1.** MIC values against Gram-negative and Gram-positive bacteria.

| Extended Data Table 1. MIC Values against Gram-positive and Gram-negative bacteria |  |  |  |  |  |  |  |  |  |  |  |  |  |  |
| --- | --- | --- | --- | --- | --- | --- | --- | --- | --- | --- | --- | --- | --- | --- |
| Strain | MIC (µg/mL) |  |  |  |  |  |  |  |  |  |  |  |  |  |
|  | 1 | 2 | 3 | 4 | 5 | 6 | 7 | 8 | 11 | 12 | 13 | 14 | 15 | 16 |
| Gram-negative bacteria |  |  |  |  |  |  |  |  |  |  |  |  |  |  |
| *Escherichia coli* ATCC 25922 | 64 | 2 | 8 | 8 | 16 | 64 | 64 | 2 | 4 | 4 | 8 | 64 | 64 | >64 |
| *Klebsiella pneumoniae* ATCC 700603 | 64 | 8 | 8 | 16 | 32 | >64 | >64 | 4 | 8 | 16 | 32 | >64 | >64 | >64 |
| *Morganella morganii* ATCC 25830 | >64 | 32 | 64 | 64 | - | - | - | - | - | - | - | - | - | - |
| *Pseudomonas aeruginosa* ATCC 9027 | >64 | 64 | 64 | >64 | >64 | >64 | >64 | 64 | - | - | - | - | - | - |
| *Acinetobacter baumannii* ATCC 19606 | >64 | >64 | >64 | >64 | >64 | >64 | >64 | >64 | >64 | >64 | >64 | >64 | >64 | >64 |
| *Enterobacter cloacae* ATCC 13047 | - | - | - | - | - | - | - | - | 8 | >64 | >64 | 64 | >64 | >64 |
| *Enterococcus faecalis* ATCC 29212 | - | - | - | - | - | - | - | - | >64 | >64 | >64 | >64 | >64 | >64 |
| Gram-positive bacteria |  |  |  |  |  |  |  |  |  |  |  |  |  |  |
| *Bacillus subtilis* ATCC 6633 | >64 | >64 | >64 | >64 | - | - | - | - | - | - | - | - | - | - |
| *Staphylococcus aureus* ATCC 29737 | >64 | >64 | >64 | >64 | >64 | >64 | >64 | >64 | >64 | >64 | >64 | >64 | >64 | >64 |

**Extended Data Table 2.** Cryo-EM data collection and refinement statistics of BAM-2 complex in MSPdH5 nanodiscs.

|  | BAM-Xenorceptide A2 (2)<br>(EMDB- 52074)<br>(PDB 9HE1) |
| --- | --- |
| Data collection and processing |  |
| Magnification | 165,000 |
| Voltage (kV) | 300 |
| Electron exposure (e<sup>-</sup>/Å<sup>2</sup>) | 40 |
| Defocus range (μm) | -0.6 to -2.0 |
| Pixel size (Å) | 0.73 |
| Symmetry imposed | C1 |
| Initial particle images (no.) | 8,425,948 |
| Final particle images (no.) | 93,541 |
| Map resolution (Å) | 3.03 |
| FSC threshold | 0.143 |
| Map resolution range (Å) | 2.4-8.4 |
| Refinement |  |
| Initial model used (PDB code) | *ab initio* |
| Model resolution (Å) | 3.3 |
| FSC threshold | 0.5 |
| Model resolution range (Å) | 2.8-6.5 |
| Map sharpening B factor (Å<sup>2</sup>) | 85.3 |
| Model composition |  |
| Non-hydrogen atoms | 11837 |
| Protein residues | 1499 |
| Ligands | 1 |
| B factors (Å<sup>2</sup>) |  |
| Protein | 189.8 |
| Ligand | 107.6 |
| R.m.s. deviations |  |
| Bond lengths (Å) | 0.003 |
| Bond angles (°) | 0.63 |
| Validation |  |
| MolProbity score | 1.27 |
| Clashscore | 3.33 |
| Poor rotamers (%) | 0.64 |
| Ramachandran plot |  |
| Favored (%) | 97.2 |
| Allowed (%) | 2.8 |
| Disallowed (%) | 0.0 |

