## Supplementary Info for "Antibiotics that Kill Gram-negative Bacteria by Restructuring the Outer Membrane Protein BamA"

| | | Page No |
|---|---|---|
| Figure S1 | Production of xenorceptide A2 (**2**) | 2 |
| Figure S2 | Production of xenorceptides using strategy 1 | 3 |
| Figure S3 | Tandem mass spectrometry (MSMS) data of xenorceptides A2–A4 (**2**–**4**) | 4 |
| Figure S4 | MSMS data of xenorceptides A5–A7 (**5**–**7**) | 5 |
| Figure S5 | MSMS data of xenorceptides A8–A10 (**8**–**10**) | 6 |
| Figure S6 | MSMS data of xenorceptides A11–A13 (**11**–**13**) | 7 |
| Figure S7 | MSMS data of xenorceptides A14–A16 (**14**–**16**) | 8 |
| Figure S8 | $^{1}$H NMR spectrum of xenorceptide A2 (**2**) | 9 |
| Figure S9 | TOCSY spectrum of xenorceptide A2 (**2**) | 10 |
| Figure S10 | Phase-sensitive NOESY spectrum of xenorceptide A2 (**2**) | 11 |
| Figure S11 | HSQC spectrum of xenorceptide A2 (**2**) | 12 |
| Figure S12 | HMBC spectrum of xenorceptide A2 (**2**) | 13 |
| Table S1 | High-resolution MS data of modified peptide products identified in this study | 14 |
| Table S2 | NMR data for xenorceptide A2 (**2**) | 15-18 |
| Table S3 | Marfey's analysis of xenorceptide A2 (**2**) | 19 |

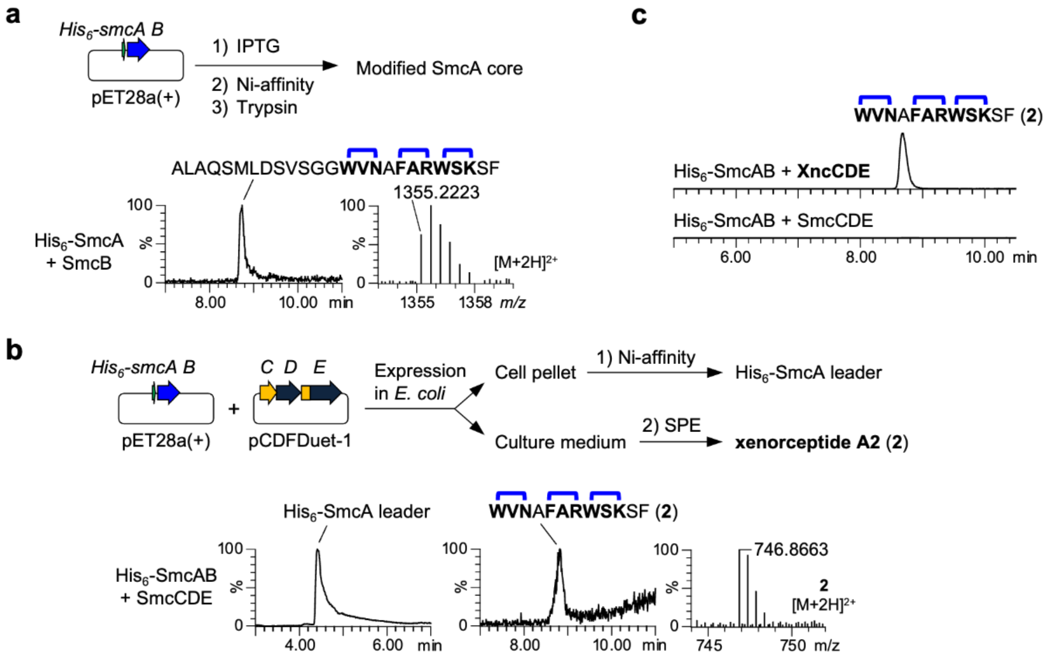

**Fig. S1. Production of xenorceptide A2 (2). a**, Coexpression of His$_6$-SmcA+SmcB. **b**, Production of natural product using a 2-vector system, His$_6$-*smcAB*/pET28a(+) + *smcCDE*/pCDFDuet-1. Extracted ion chromatograms (EICs) show cleaved leader (left) and natural product (right) detected only when coexpressed with SmcCDE. HR-MS for **2** is shown. **c**, Enhanced production of **2** by coexpressing *His$_6$-smcAB*/pET28 with *xncCDE*/pCDFDuet-1.

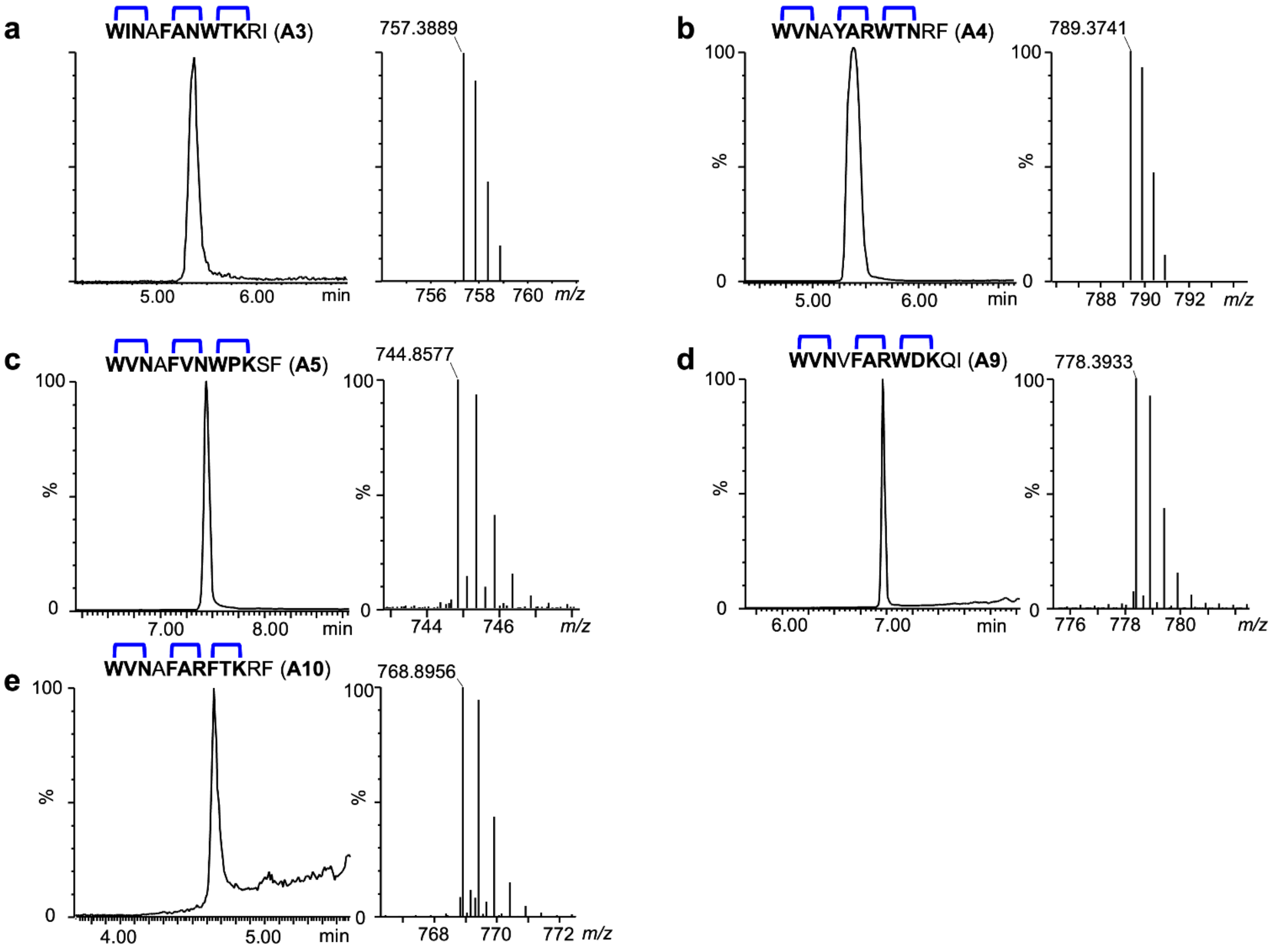

**Fig. S2. Production of xenorceptides using strategy 1. a**–**e**, Production of xenorceptides A3–A5, A9, and A10 (**3**–**5, 9**, **10**) by the coexpression of His$_6$-*xyeAB*/pET28a(+) and *xncCDE*/pCDFDuet-1 vectors. EIC (left) and MS (right) corresponding to the end products are shown.

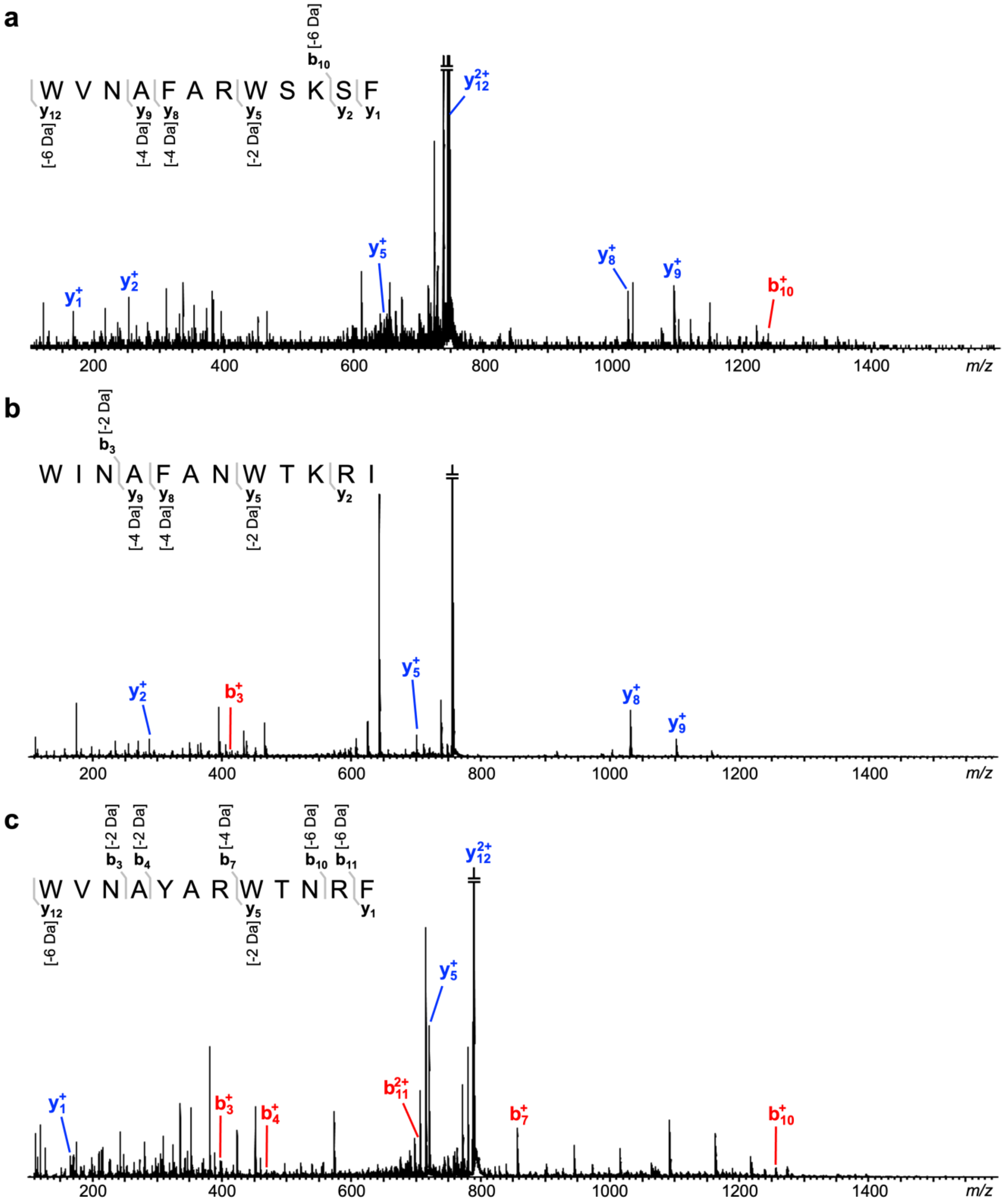

**Fig. S3. Tandem mass spectrometry (MSMS) data of xenorceptides A2–A4 (2–4).** MSMS data of **2** (a), **3** (b), **4** (c) localize the -2 Da mass losses to respective three-residue motifs. Mass range = *m/z* 100–1600, Collision energy = 23 V, 30 V, and 30 V for **2**, **3,** and **4**, respectively.

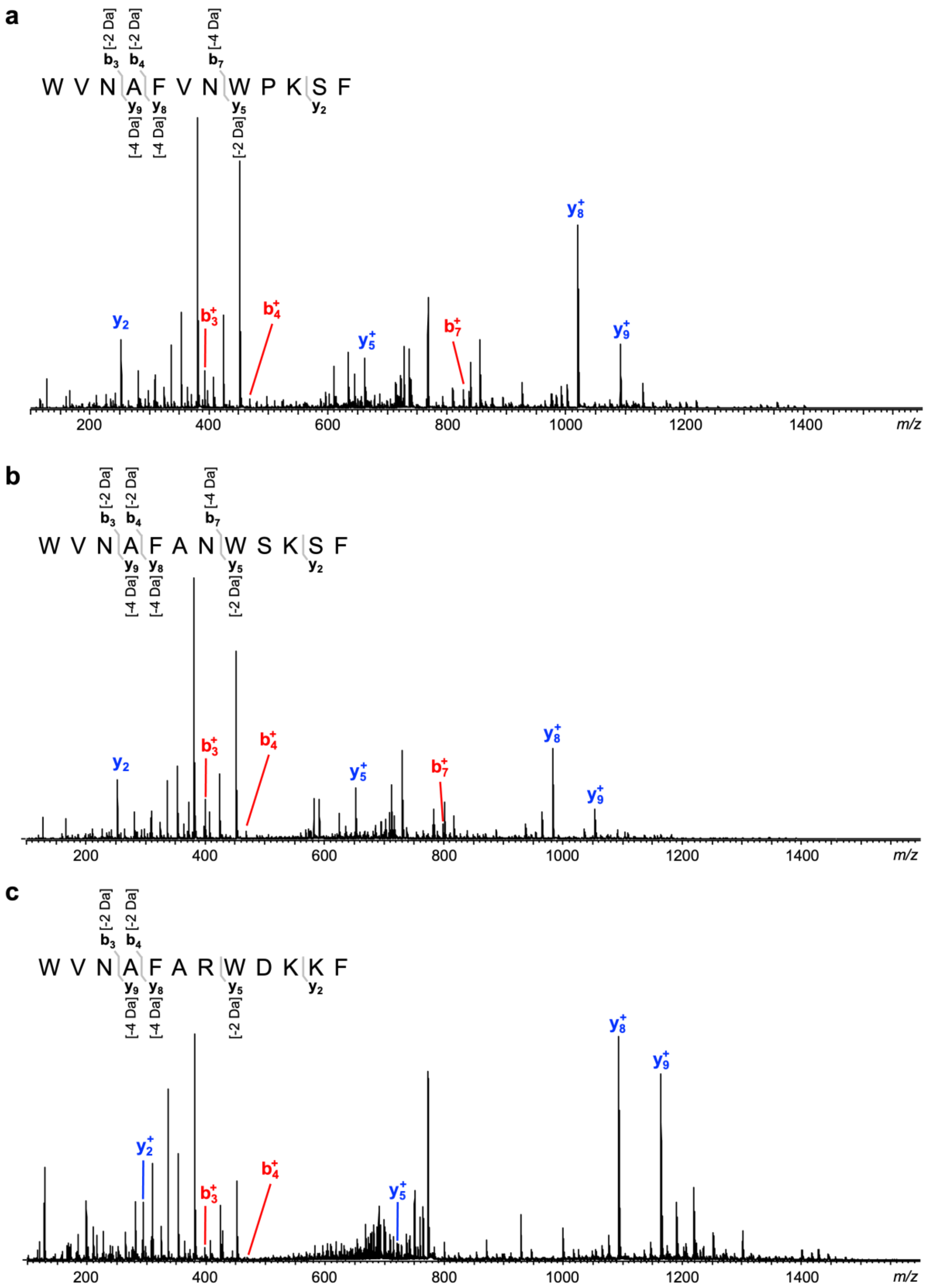

**Fig. S4. MSMS data of xenorceptides A5–A7 (5–7).** MSMS data of **5** (a), **6** (b), **7** (c) localize the -2 Da mass losses to respective three-residue motifs. Mass range = *m/z* 100–1600, Collision energy = 30 V, 30 V, and 35 V for **5**, **6,** and **7**, respectively.

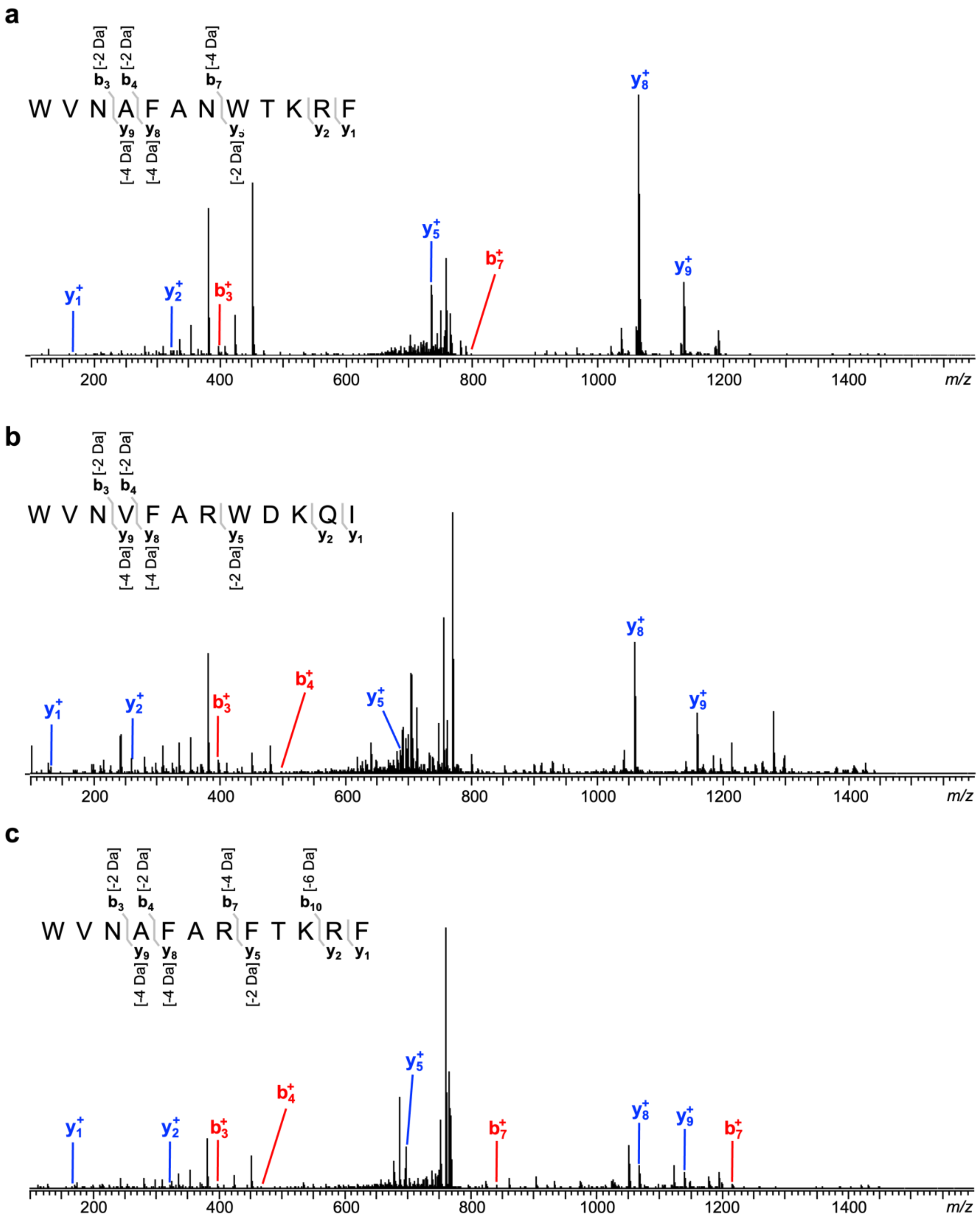

**Fig. S5. MSMS data of xenorceptides A8–A10 (8–10).** MSMS data of **8** (a), **9** (b), **10** (c) localize the -2 Da mass losses to respective three-residue motifs. Mass range = *m/z* 100–1600, Collision energy = 30 V, 34 V, and 34 V for **8**, **9,** and **10**, respectively.

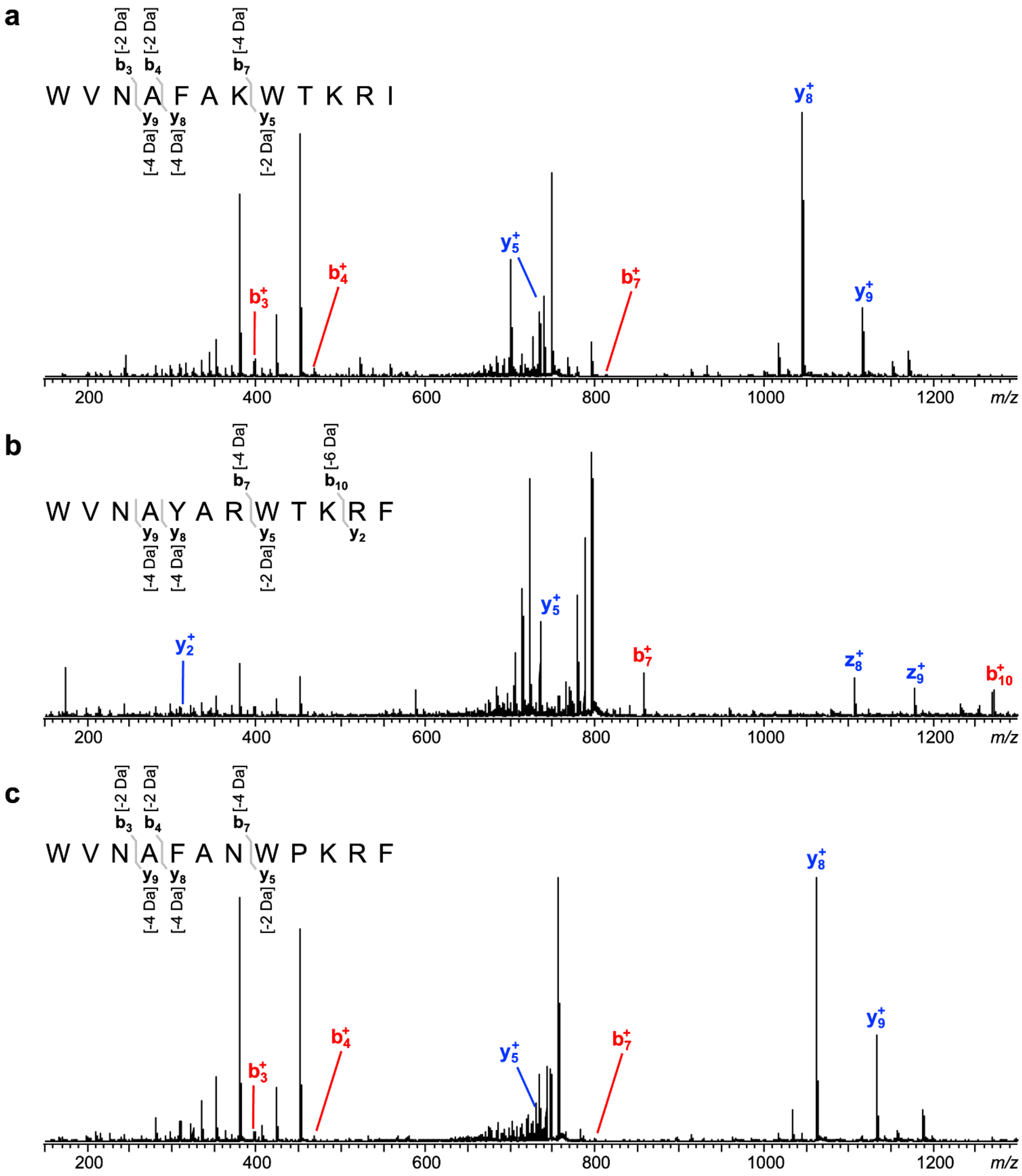

**Fig. S6. MSMS data of xenorceptides A11–A13 (11–13).** MSMS data of **11** (a), **12** (b), **13** (c) localize the -2 Da mass losses to respective three-residue motifs. Mass range = *m/z* 100–1600, Collision energy = 30 V, 35 V, and 30 V for **11**, **12,** and **13**, respectively.

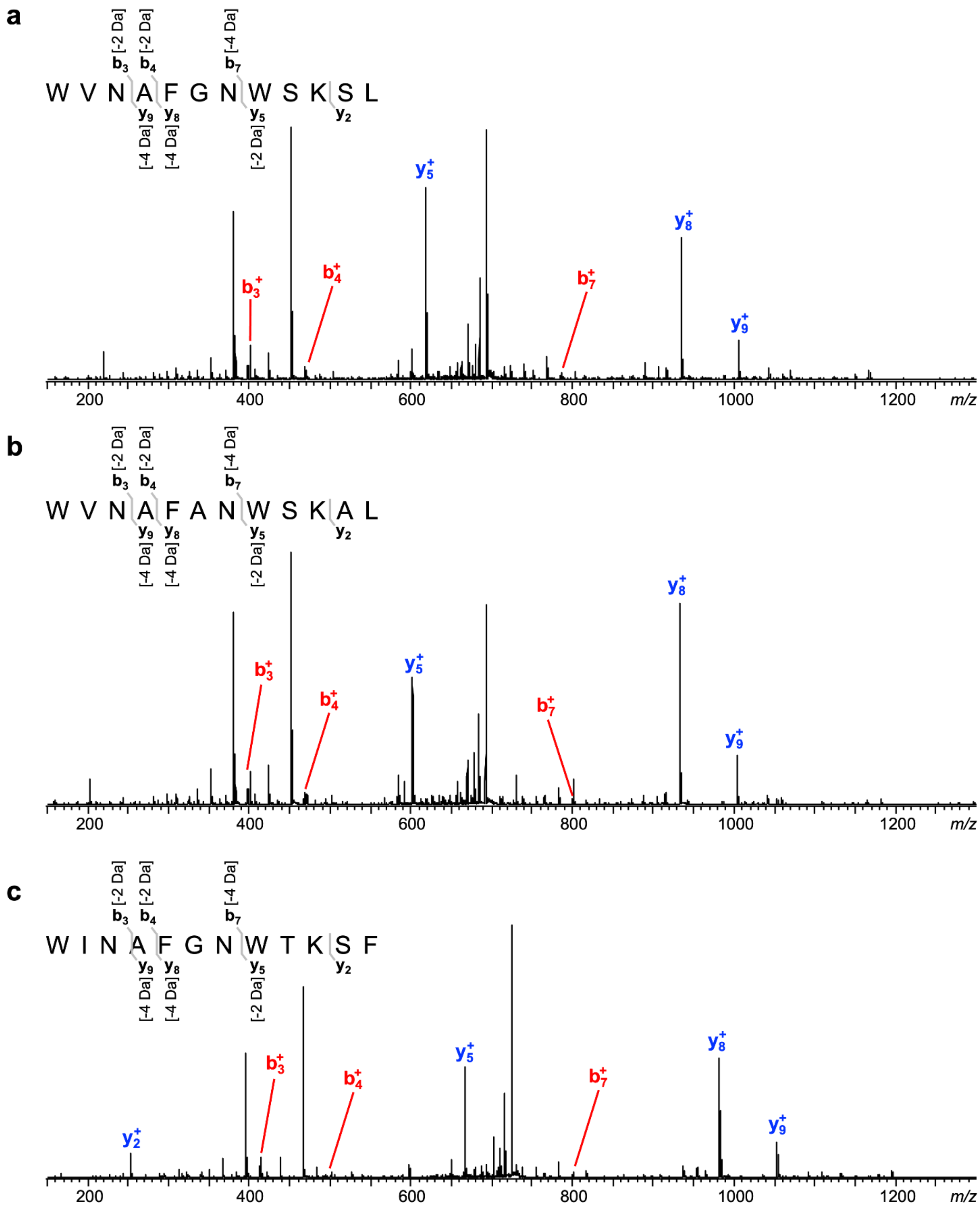

**Fig. S7. MSMS data of xenorceptides A14–A16 (14–16).** MSMS data of **14** (a), **15** (b), **16** (c) localize the -2 Da mass losses to respective three-residue motifs. Mass range = *m/z* 100–1600, Collision energy = 25 V for **14–16**.

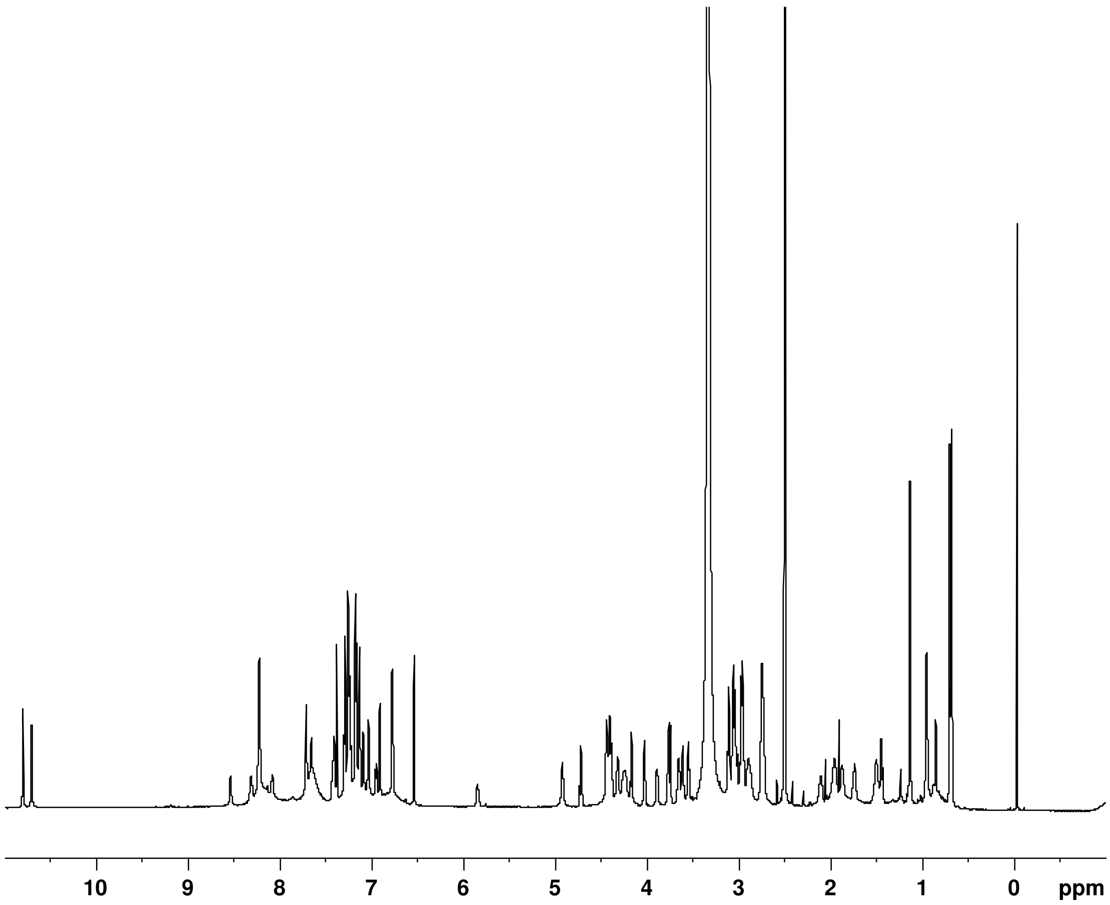

**Fig. S8. [1]H NMR spectrum of 2.** Acquired at 800 MHz in DMSO-$d_6$ at 298 K.

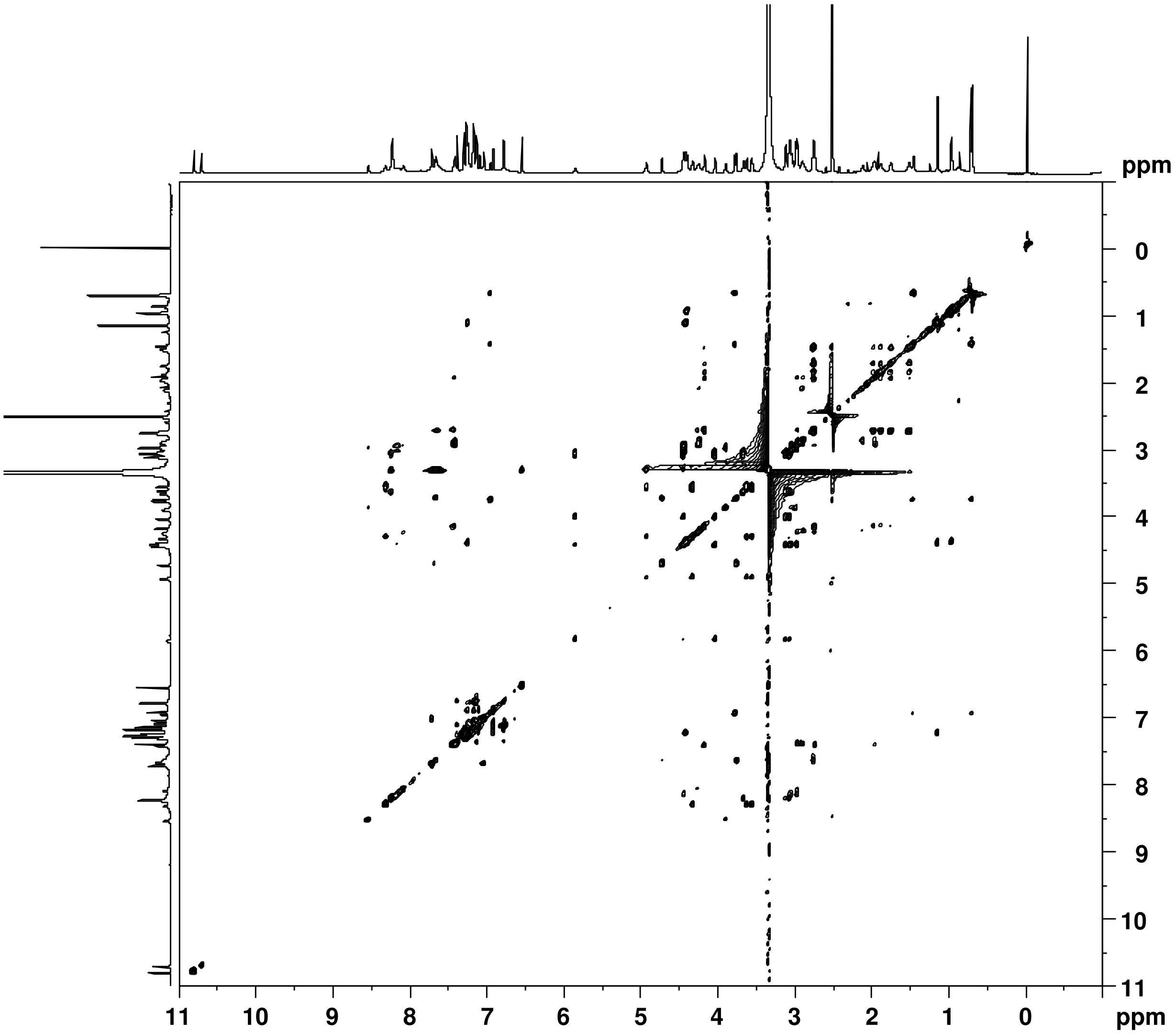

**Fig. S9. TOCSY spectrum of 2.** Acquired at 800 MHz in DMSO-$d_6$ at 298 K.

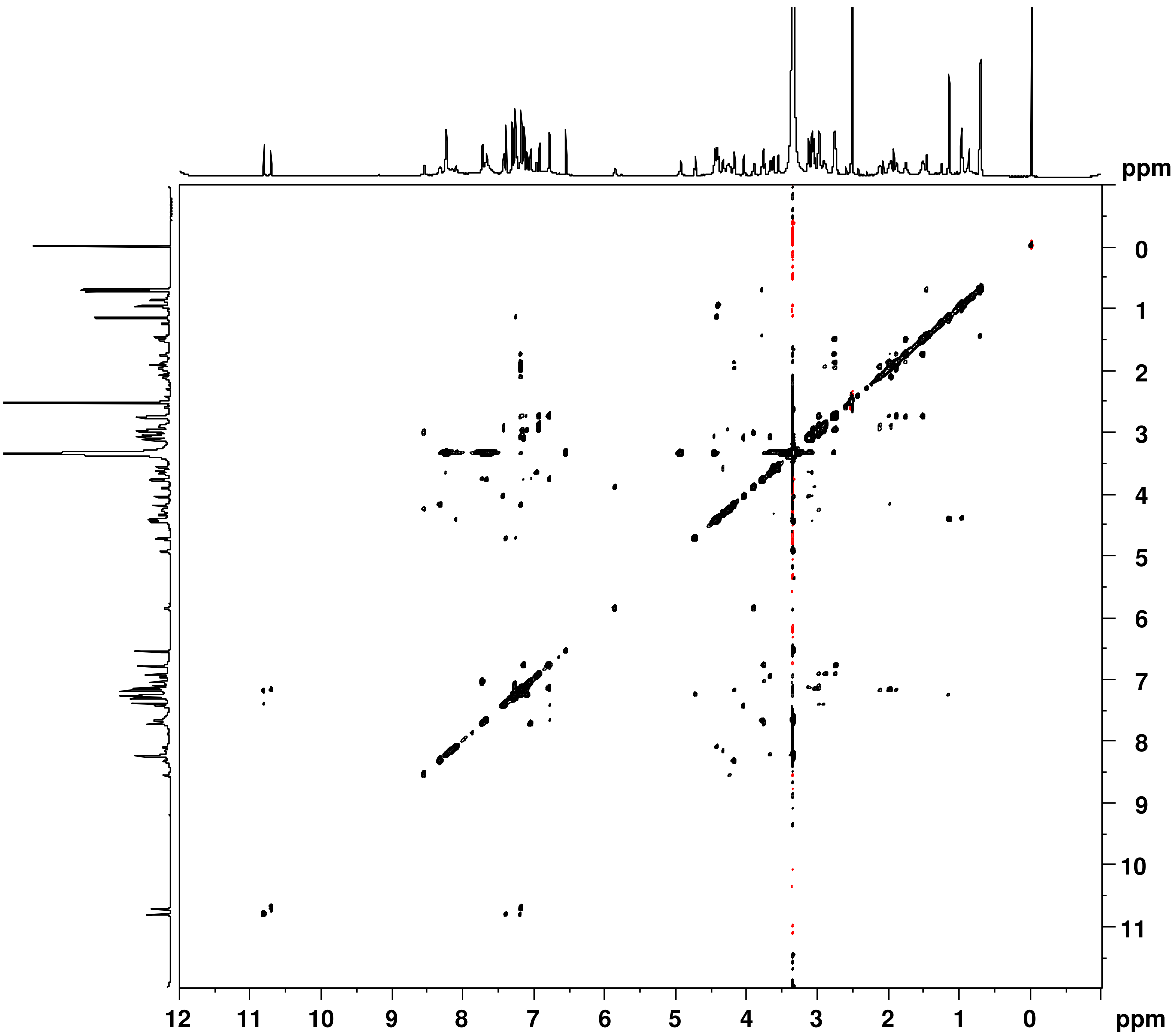

**Fig. S10. Phase-sensitive NOESY spectrum of 2.** Acquired at 800 MHz in DMSO-$d_6$ at 298 K.

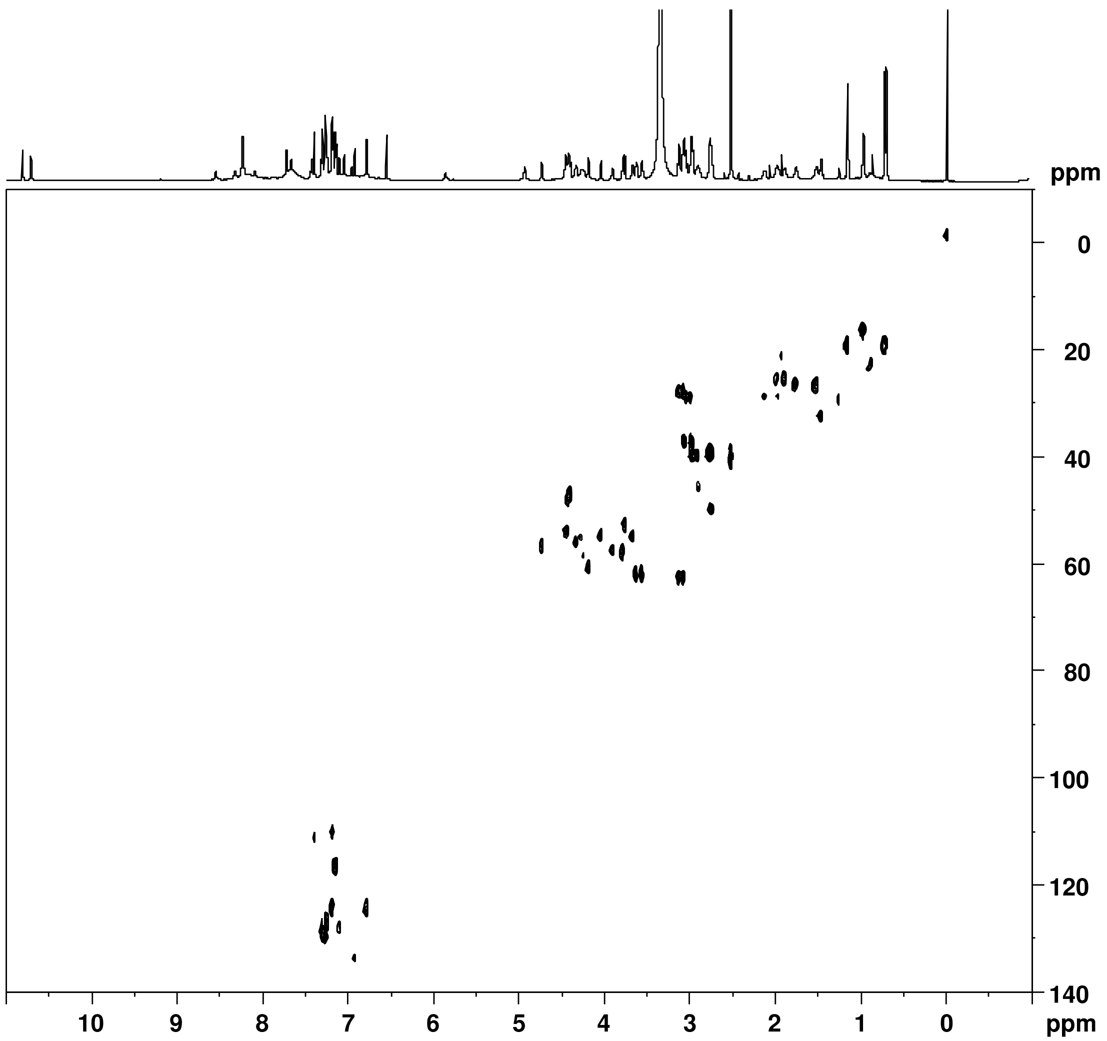

**Fig. S11. HSQC spectrum of 2.** Acquired at 800 MHz in DMSO-$d_6$ at 298 K.

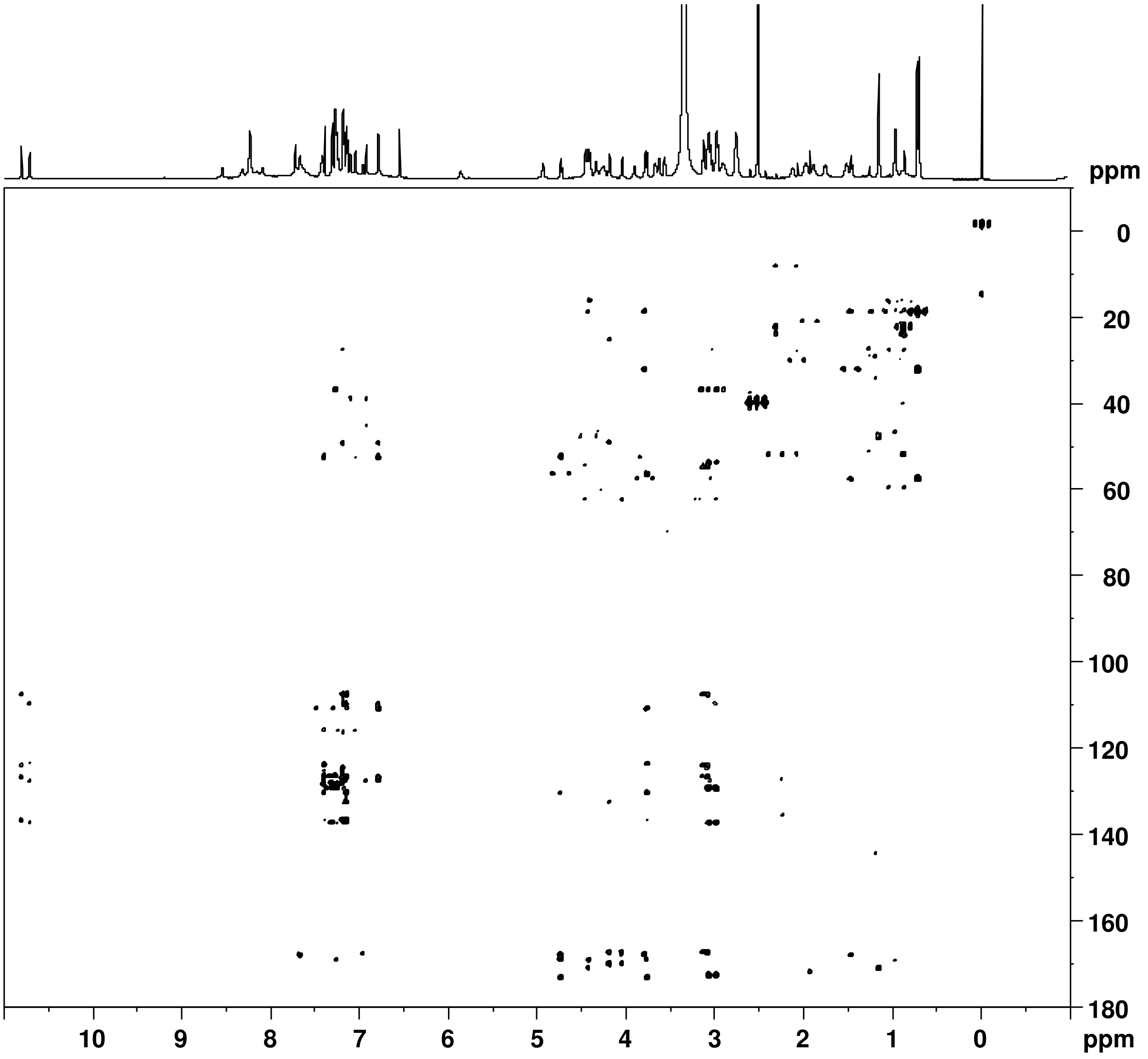

**Fig. S12. HMBC spectrum of 2.** Acquired at 800 MHz in DMSO-$d_6$ at 298 K.

**Table S1.** High-resolution MS data of modified peptide products identified in this study.

| Compound # | Sequence[a] | Charge State | Calculated mass (monoisotopic) | Observed mass (monoisotopic) | Δppm |
|---|---|---|---|---|---|
| **1** | WINAFGNWERAFH | $[M+2H]^{2+}$ | 821.3709 | 821.3721 | 1.5 |
| **2** | WVNAFARWSKSF | $[M+2H]^{2+}$ | 746.8597 | 746.8602 | 0.7 |
| **3** | WINAFANWTKRI | $[M+2H]^{2+}$ | 757.3886 | 757.3889 | 0.4 |
| **4** | WVNAYARWTNRF | $[M+2H]^{2+}$ | 789.3735 | 789.3741 | 0.8 |
| **5** | WVNAFVNWPKSF | $[M+2H]^{2+}$ | 744.8566 | 744.8577 | 1.5 |
| **6** | WVNAFANWSKSF | $[M+2H]^{2+}$ | 725.8306 | 725.8322 | 2.2 |
| **7** | WVNAFARWDKKF | $[M+2H]^{2+}$ | 781.3886 | 781.3898 | 1.5 |
| **8** | WVNAFANWTKRF | $[M+2H]^{2+}$ | 767.3729 | 767.3740 | 1.4 |
| **9** | WVNVFARWDKQI | $[M+2H]^{2+}$ | 778.3939 | 778.3933 | -0.8 |
| **10** | WVNAFARFTKRF | $[M+2H]^{2+}$ | 768.8966 | 768.8956 | -1.3 |
| **11** | WVNAFAKWTKRI | $[M+2H]^{2+}$ | 757.4068 | 757.4058 | -1.3 |
| **12** | WVNAYARWTKRF | $[M+2H]^{2+}$ | 796.3995 | 796.4008 | 1.7 |
| **13** | WVNAFANWPKRF | $[M+2H]^{2+}$ | 765.3755 | 765.3766 | 1.5 |
| **14** | WVNAFGNWSKSL | $[M+2H]^{2+}$ | 701.8306 | 701.8297 | -1.2 |
| **15** | WVNAFANWSKAL | $[M+2H]^{2+}$ | 700.8409 | 700.8425 | 2.2 |
| **16** | WINAFGNWTKSF | $[M+2H]^{2+}$ | 732.8384 | 732.8412 | 3.8 |
| **S1** | ALAQSMLDSVSGGWVNAFA RWSKSF | $[M+3H]^{3+}$ | 903.7675 | 903.7661 | -1.5 |

[a]Cyclized three-residue motifs are indicated in red.

**Table S2.** NMR data for xenorceptide A2 (**2**).

| Residue | Position | $^{1}H^{a}$ | $^{13}C^{a,b}$ | COSY | HMBC (H to C) | NOESY |
|---|---|---|---|---|---|---|
| Trp1 | C=O | | 168.3 | | | |
| | $NH_2$ | 8.22 | | Hα | | Trp1-Hα |
| | α | 3.65 | 54.5 | $NH_2$, Hβ | | Trp1-$NH_2$, Trp1-Hβa, Tryp1-Hβb, Val2-NH |
| | β | 3.10 (Ha) | 27.0 | Hα | Trp1-Ca, Trp1-C2, Trp1-C3, Trp1-C3a | Trp1-Hα, Trp1-H4 |
| | | 3.06 (Hb) | | | | Trp1-Hα, Trp1-H2 |
| | 1 | 10.80 | | H2 | Trp1-C2, Trp1-C3, Trp1-C3a, Trp1-C7a | Trp1-H2, Trp1-H7 |
| | 2 | 7.18 | 124.6 | H1 | Trp1-C3a, Trp1-C7a | Trp1-H1, Tryp1-Hβb |
| | 3 | | 108.0 | | | |
| | 3a | | 127.2 | | | |
| | 4 | 7.13 | 116.4 | H5 | Trp1-C3, Trp1-C3a, Trp1-C6, Trp1-C7a | Trp1-Hβa, Trp1-H5 |
| | 5 | 6.77 | 124.2 | H4, H7 | Trp1-C3a, Trp1-C7 | Trp1-H4, Asn3-NH, Asn3-Hβ |
| | 6 | | 130.9 | | | |
| | 7 | 7.38 | 110.7 | H5 | Trp1-C3a, Trp1-C5, Asn3-Cb | Trp1-H1 |
| | 7a | | 137.1 | | | |
| Val2 | C=O | | 168.5 | | | |
| | NH | 6.94 | | Hα | Trp1-C=O | Trp1-Hα, Val2-Hβ |
| | α | 3.77 | 57.0 | NH, Hβ | Val2-C=O, Val2-Cβ, Val2-Cγ-M1 | Val2-Hβ, Val2-Hγ-M1, Asn3-NH |
| | β | 1.45 | 31.9 | Hα, Hγ, Hγ–M1, Hγ–M2 | Val2-C=O, Val2-Cα, Val2-Cγ-M1 | Val2-Hγ-M1, Val2-Hγ–M2 |
| | γ-M1 | 0.70 | 18.4 | Hβ | Val2-Cα, Val2-Cβ | Val2-Hβ |
| | γ-M2 | 0.68 | 18.4 | Hβ | Val2-Cα, Val2-Cβ | Val2-Hβ |
| Asn3 | C=O | | 169.6 | | | |
| | NH | 7.67 | | Hα | Val2-C=O | Trp-H5, Val2-Hα |
| | α | 4.71 | 55.9 | NH, Hβ | Val2-C=O, Asn3-Cβ, Asn3-$CONH_2$, Asn3-C=O | Ala4-NH |
| | β | 3.74 | 52.0 | Hα | Trp1-C5, Trp1-C6, Trp1-C7, Asn3-$CONH_2$, Asn3-Cα, Asn3-C=O | Trp1-H5 |
| | $CONH_2$ | | 173.8 | | | |
| Ala4 | C=O | | 171.7 | | | |

| | | | | | | |
|---|---|---|---|---|---|---|
| | NH | 7.24 | | Hα | Asn3-C=O | Asn3-Hα, Ala4-Hα, Ala4-Hβ |
| | α | 4.40 | 48.1 | NH, Hβ | Ala4-Cβ | Ala4-NH, Ala4-Hβ, Phe5-NH |
| | β | 1.13 | 18.4 | Hα, Hγ | Ala4-Cα, Ala4-C=O | Ala4-NH, Ala4-Hα Phe5-NH |
| Phe5 | C=O | | n.d.[c] | | | |
| | NH | 8.08 | | Hα | | Ala4-Hα, Ala4-Hβ, Phe5-Hα, Phe5-Hβ |
| | α | 4.26 | 54.5 | NH, Hβ | | Phe5-Hα, Phe5-Hβ, Phe5-H6, Ala6-NH |
| | β | 2.96 (Ha) | 39.5 | Hα | | Phe5-NH, Phe5-H2, Phe5-H6 |
| | | 2.73 (Hb) | | | | Phe5-NH, Phe5-H2 |
| | 1 | | n.d.[c] | | | |
| | 2 | 6.91 | 133.3 | H5 | Phe5-Cβ, Phe2-C6, Arg7-Cβ | Phe5-Hβa, Phe5-Hβb, Arg7-NH, Arg7-Hβ |
| | 3 | | n.d.[c] | | | |
| | 4 | 7.17 | 123.4 | H6 | Phe2-C2, Phe2-C6 | Arg7-Hγ |
| | 5 | 7.25 | 129.1 | H2 | | Phe5-H4, Phe5-H6 |
| | 6 | 7.09 | 127.6 | H3 | | Phe5-H5, Phe5-Hα, Phe5-Hβa |
| Ala6 | C=O | | 169.9 | | | |
| | NH | 7.86 | | Hα | | Phe5-Hα |
| | α | 4.38 | 46.4 | NH, Hβ | Ala6-Cβ | Ala6-Hβ, Arg7-NH |
| | β | 0.95 | 15.8 | Hα | Ala6-Cα, Ala6-C=O | Ala6-Hα |
| Arg7 | C=O | | n.d.[c] | | | |
| | NH | 7.58 | | Hα | | Phe5-H2, Ala6-Hα |
| | α | 4.23 | 58.3 | NH, Hβ | | Arg7-Hβ, Arg7-Hγ, Trp8-NH |
| | β | 2.87 | 45.7 | Hα | Arg7-Cδ | Phe5-H2, Arg7-Hα, Trp8-NH |
| | γ | 2.10 (Ha) | 28.3 | | | Phe5-H4, Arg7-Hα |
| | | 1.94 (Hb) | | | | Phe5-H4, Arg7-Hα |
| | δ | 2.96 | 37.2 | | | |
| | C (guanidine) | | n.d.[c] | | | |
| Trp8 | C=O | | 170.6 | | | |
| | NH | 8.53 | | Hα | | Arg7-Hα, Arg7-Hβ, Trp8-Hβ |
| | α | 3.89 | 57.0 | NH, Hβ | | Trp8-Hβ, Thr9-NH |

| | | | | | | |
|---|---|---|---|---|---|---|
| | β | 3.02 (Ha) | 28.3 | Hα | Trp8-C3 | Trp8-NH, Trp8-Hα |
| | | 2.98 (Hb) | | | | |
| | 1 | 10.70 | | H2 | Trp8-C2, Trp8-C3, Trp8-C3a, Trp8-C7a | Trp8-H2, Trp8-H7 |
| | 2 | 7.16 | 123.9 | H1 | Trp8-C7a | Trp8-NH |
| | 3 | | 110.3 | | | |
| | 3a | | 128.2 | | | |
| | 4 | 7.14 | 115.9 | H5 | Trp8-C6, Trp8-C7α | Trp8-H5 |
| | 5 | 6.77 | 124.6 | H4 | Trp8-C3a, Trp8-C7 | Trp8-H4, Lys10-NH, Lys10-Hβ |
| | 6 | | 132.9 | | | |
| | 7 | 7.17 | 110.4 | | Arg10-Cβ | Trp8-H1, Lys10-Hα |
| | 7a | | 137.8 | | | |
| Ser9 | C=O | | 167.9 | | | |
| | NH | 5.84 | | Hα | | Trp8-Hβ |
| | α | 4.03 | 54.5 | NH, Hβ | Trp8-C=O, Ser9-Cβ, Ser9-C=O | Ser9-Hβ, Lys10-NH |
| | β | 3.09 | 62.0 | Hα | Ser9-C=O | Ser9-NH, Lys10-NH |
| Lys10 | C=O | | 170.7 | | | |
| | NH | 7.42 | | Hα | | Trp8-H5, Ser9-Hα, Lys10-Hα, Lys10-Hβ |
| | α | 4.16 | 60.7 | NH, Hβ | Trp8-C6, Ser9-C=O, Lys10-C=O, Lys10-Cβ, Lys10-Cγ | Trp8-H7, Lys10-NH, Lys10-Hγa, Lys10-Hγb, Ser11-NH |
| | β | 2.73 | 49.5 | Hα, Hγ | | Trp8-H5, Lys10-Hα, Lys10-Hγa, Lys10-Hgb, Lys10-Hδa, Lys10-Hδb |
| | γ | 1.97 (Ha) | 24.5 | Hβ, Hδ | | Lys10-Hα, Lys10-Hβ |
| | | 1.86 (Hb) | | | | Lys10-Hα, Lys10-Hβ |
| | δ | 1.74 (Ha) | 25.7 | Hγ, Hε | | Lys10-Hβ |
| | | 1.50 (Hb) | | | | Lys10-Hβ |
| | ε | 2.75 | 39.4 | $NH_2$, Hδ | | Lys10-$NH_2$ |
| | $NH_2$ | 7.64 | | Hε | | Lys10-Hε |
| Ser11 | C=O | | n.d.[c] | | | |
| | NH | 8.31 | | Hα | | Lys10-Cα, Ser11-Hβ |
| | α | 4.32 | 55.7 | NH, Hβ | | Ser11-Hβ, Phe12-NH |

| | | | | | | |
|---|---|---|---|---|---|---|
| | β | 3.58 | 61.9 | Hα, Hγ | | Ser11-NH |
| Phe12 | C=O | | 173.2 | | | |
| | NH | 8.15 | | Hα | | Ser11-Hα, Phe12-Hβb |
| | α | 4.42 | 53.3 | NH, Hβ | | Phe12-NH |
| | β | 3.05 | 36.9 | | Phe12-Cα, Phe12-C1, Phe12-C2, Phe12-C=O | |
| | | 2.96 | | | | Phe12-NH |
| | 1 | | 137.3 | Hα, Hγ | | |
| | 2 | 7.26 | 129.2 | Hβ, Hδ | Phe12-Cβ, Phe12-C4, Phe12-C6 | |
| | 3 | 7.29 | 128.8 | Hβ | Phe12-C1, Phe12-C5 | |
| | 4 | 7.24 | 127.0 | Hγ | Phe12-C2, Phe12-C6 | |
| | 5 | 7.29 | 128.7 | | Phe12-C1, Phe12-C5 | |
| | 6 | 7.26 | 129.2 | | Phe12-Cβ, Phe12-C4, Phe12-C6 | |

[a]800 MHz in DMSO-$d_6$ at 298 K. [b]Assigned by HSQC and HMBC. [c]Not detected.

**Table S3.** Marfey's analysis of xenorceptide A2 (**2**).

| | Retention time (min)[a] | | |
|---|---|---|---|
| Amino acid | L-DVA-std | D-DVA-std | Hydrolysate of **2**[b] |
| L-Ala | 9.13 | 10.57 | 9.13 |
| L-Arg | 4.28 | 3.92 | n.d.[c] |
| L-Asp | 7.63 | 7.98 | n.d.[c] |
| L-Lys | 4.01 | 3.64 | n.d.[c] |
| L-Phe | 11.93 | 13.87 | 11.93 |
| L-Ser | 7.31 | 7.66 | 7.31 |
| L-Trp | 11.53 | 12.77 | n.d.[c] |
| L-Val | 10.60 | 13.04 | n.d.[c] |

[a]Analytical condition: MS polarity = negative; column: Kinetex XB-C18, 2.6 μm, 150 x 4.6 mm; flow rate: 0.50 mL/min; column temperature: 50 °C; mobile phase/gradient: 30% $H_2O/CH_3CN$ + 0.1% formic acid isocratic for 2 min followed by linear gradient to 70% $H_2O/CH_3CN$ + 0.1% formic acid over 17 min.
[b]Derivatized with L-FDVA.
[c]Not detected.
